# Voltage-gated ion channels mediate the electrotaxis of glioblastoma cells in a hybrid PMMA/PDMS microdevice

**DOI:** 10.1101/2020.02.14.948638

**Authors:** Hsieh-Fu Tsai, Camilo IJspeert, Amy Q. Shen

## Abstract

Transformed astrocytes in the most aggressive form cause glioblastoma, the most common cancer in central nervous system with high mortality. The physiological electric field by neuronal local field potentials and tissue polarity may guide the infiltration of glioblastoma cells through the electrotaxis process. However, microenvironments with multiplex gradients are difficult to create. In this work, we have developed a hybrid microfluidic platform to study glioblastoma electrotaxis in controlled microenvironments with high throughput quantitative analysis by a machine learning-powered single cell tracking software. By equalizing the hydrostatic pressure difference between inlets and outlets of the microchannel, uniform single cells can be seeded reliably inside the microdevice. The electrotaxis of two glioblastoma models, T98G and U-251MG, require optimal laminin-containing extracellular matrix and exhibits opposite directional and electro-alignment tendencies. Calcium signaling is a key contributor in glioblastoma pathophysiology but its role in glioblastoma electrotaxis is still an open question. Anodal T98G electrotaxis and cathodal U-251MG electrotaxis require the presence of extracellular calcium cations. U-251MG electrotaxis is dependent on the P/Q-type voltage-gated calcium channel (VGCC) and T98G is dependent on the R-type VGCC. U-251MG and T98G electrotaxis are also mediated by A-type (rapidly inactivating) voltage-gated potassium channels and acid-sensing sodium channels. The involvement of multiple ion channels suggests that the glioblastoma electrotaxis is complex and patient-specific ion channel expression can be critical to develop personalized therapeutics to fight against cancer metastasis. The hybrid microfluidic design and machine learning-powered single cell analysis provide a simple and flexible platform for quantitative investigation of complicated biological systems.

## I. INTRODUCTION

Glioma is one of the most common types of brain cancer and the aggressive form of it, glioblastoma, contributes to poor prognosis, high mortality, and high probability of recurrence^1,2^, due to the infiltration nature of the disease. The highly infiltrative ability of glioblastoma originates from the invasive/migratory ability of glioma stem cells or brain tumor initiating cells^3,4^. Not only glioma cells are important, but also the microenvironment in the brain helps shaping the heterogeneity of the glioma^5^. The glioma cells interact with the extracellular matrix (ECM), glial cells, and immune cells in the brain and mediate the formation of peri-vascular, peri-necrotic, and invasive tumor microenvironments^6–9^. Understanding the molecular mechanisms of the invasiveness in glioma cells with respect to the tumor microenvironment is vital for developing new therapeutic options and improving patient outcome^10,11^.

In the brain, glial cells are immersed in an electric field created by tissue polarity from brain macrostructures as well as the local field potentials which are established from the action potentials fired by the neurons^12^. A weak endogenous electric current has been shown to serve as a guidance cue for neuroblast migration from the sub-ventricular zone in mouse^13^, a region speculated as the origin of glioma tumorigenesis^14^. Thus, the physiological electric fields in the brain may play an important role in mediating the glioma tumorigenesis and invasion^15–19^. Cells sense the electric field by bioelectrical activation of voltage-sensitive proteins, mechanosensing due to electrokinetic phenomena, or activated chemical signaling due to electrokinetically polarized membrane receptors (Supplementary FIG. S.1). The voltage gradient creates a large voltage drop at cellular membrane which can directly activate voltage sensitive proteins such as voltage-gated ion channels that are most commonly expressed on excitable membranes at neuronal synapses and neuro-muscular junctions^20^. Among the voltagegated ion channels in the brain, calcium channels are especially important as calcium influx plays a pivotal role in cellular signaling^21,22^. Calcium signaling is also important in glioma cell proliferation, resistance to therapy, and metastasis^23–27^. Whether or not the calcium signaling in glioma is mediated by electric field is still an open question.

Conventional *in vitro* electrical stimulation systems for studying cell responses in electrical microenvironments are bulky and the experimental throughput is limited^28,29^. To overcome these limitations, a robust high-throughput *in vitro* platform that creates stable electrical stimulation of cells and interfaces with automated microscopy is a prerequisite for rapid screening of targets and identifying molecular mechanisms. To this end, we have developed a hybrid poly(methylmethacrylate)/poly(dimethylsiloxane) (PMMA/PDMS) microfluidic platform to reliably study glioblastoma single cell migration under high-throughput dcEF stimulations that multiple antagonists can be tested simultaneously to identify molecular mechanisms. Quantitative single cell migration analysis is carried out by extracting cell migration metrics such as the directedness, orientation, or speed using a robust machine learning-powered cell segmentation/tracking/analysis software with stain-free phase contrast microscopy^30^. Using the hybrid microfluidic platform, the role of voltage-gated calcium channels in calcium signaling pathways of glioblastoma electrotaxis are investigated.

## II. RESULTS

### A. Uniform single cell seeding by submerged manipulation in hybrid multiple electric field chip (HMEFC)

To analyze single cell migration in microchannels, cells must be seeded sparsely and allowed to adhere and culture reliably^31,32^. However, it is known that uneven distribution of cells due to fluid flow, convection in suspension and vessel movement after seeding can cause aggregation and differentiation of cells^33^. Different cell loading methods affecting cell distribution in single cell migration experiments are investigated, such as tip loading method, tip injection method, and a pressure-balanced submerged cell seeding, as shown in FIG. 1.

**FIG. 1.**
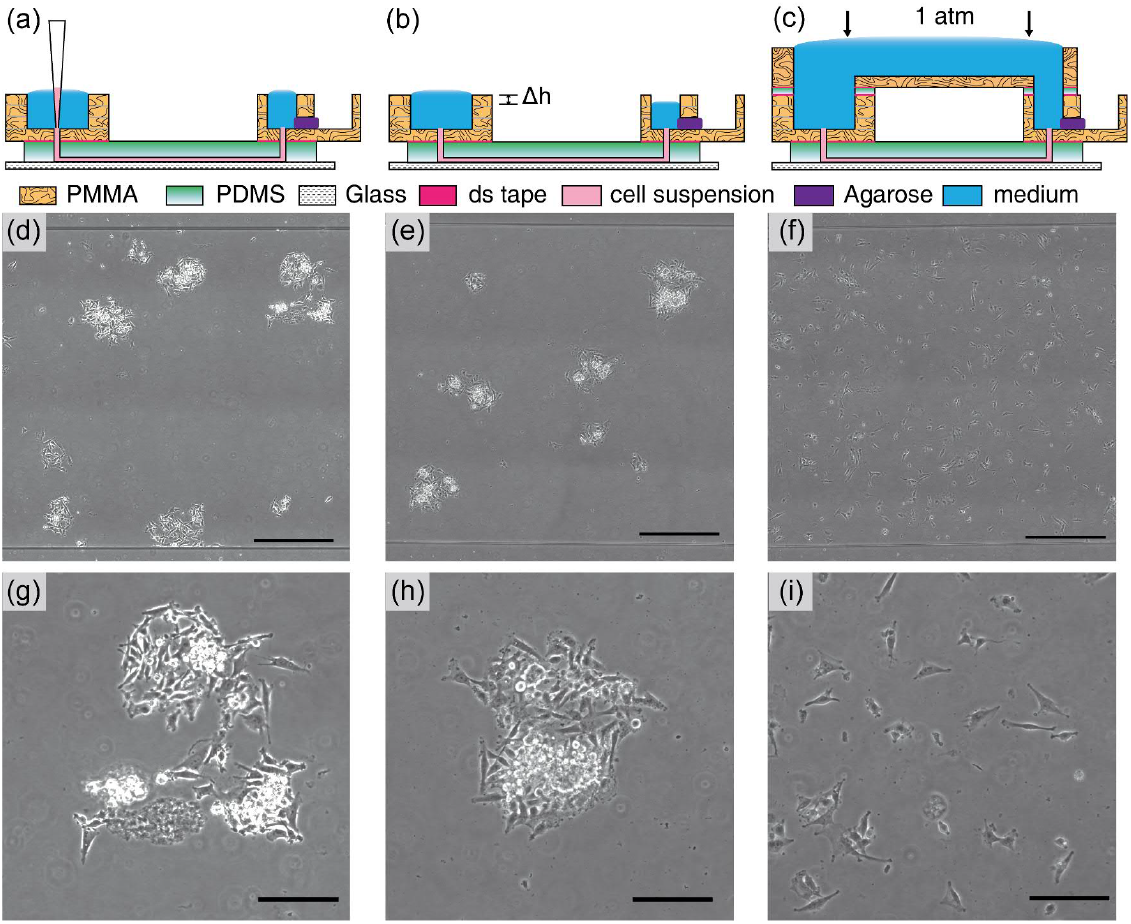
The results of U-251MG cell seeding inside microchannels by various methods. (a, d, g, j) In tip loading method, cells are introduced by using gravitational flow with micropipette tips. The cells can flocculate inside the tips and in microchannels as illustrated in (d). The microscopy image of seeded cells is shown in (g) and magnified in (j); (b, e, h, k) In tip injection method, cells are injected into the channels and tips are removed. The small hydrostatic pressure differences between the inlet/outlet (shown as Δh) will contribute to hydrodynamic flow and disturb the cell distribution, causing non-uniform cell distribution and aggregates as shown in (e). The microscopy image of seeded cells is shown in (h) and magnified in (k); (c, f, i, l) In our pressure-balanced sub-merged cell seeding method, the hydrostatic pressure difference is eliminated. The injected cells remain uniform through-out the channel as shown in (f). The microscopy image of seeded cells is shown in (i) and magnified in (l). The uniform and sparse cell seeding method is suitable for many different applications from single cell tracking, ensembled cell studies to cell assembly. The scale bars in (g, h, i) represent 500 *μ*m. The scale bars in (j, k, l) represent 200 *μ*m.

In tip loading method (FIG. 1.a), the cells flocculate in the small pipet tip and cannot be dispersed uniformly in the microchannel (FIG. 1.d, g, j). In tip injection method (FIG. 1.b), the cells are originally injected in the channels with uniform cell distribution. However, with-out balancing the microchannel inlet/outlet pressure, the minute hydrostatic pressure difference between inlets and outlets generates a small pressure-driven flow that displaces cells which lead to cell aggregates (FIG. 1.e, h, k). Furthermore, in tip injection method, due to the small dimension of the punched holes at inlet/outlet interfaces, bubbles are easily trapped and may be introduced into microchannels, disrupting fluid advection and chemical transport.

By submerging inlets and outlets underwater using a reversibly bonded top reservoir and balancing the pressure between inlets and outlets (FIG. 1.c), air bubbles can be avoided and pressure-driven flow is prevented from affecting cell distribution. Moreover, using this cell seeding method, only minute amount of cells is needed (the volume of the microchannel). Uniformly distributed single cell seeding across the entire microchannel is obtained for single cell migration experiments (FIG. 1.f, i, l). The top reservoir in our submerged cell seeding setup can be easily removed after cells are seeded and can be adapted to a wide range of microfluidic chips.

### B. Glioblastoma electrotaxis requires optimal laminin-containing ECM

An effective ECM coating on the substrate is essential for cell adhesion and formation of focal adhesions for cell migration^34,35^. Glioblastoma can be molecularly classified into proneural, neural, classical, and mesenchymal types according to The Cancer Genome Atlas (TCGA)^36^. We use two glioblastoma cell models, T98G and U-251MG, which are both of caucasian male origin and classified as mesenchymal type with p53 mutant genotype^37,38^. The adhesion and electrotaxis of T98G and U-251MG glioblastoma cell lines on various ECMs are tested in a double-layer hybrid multiple electric field chip (HMEFC) based on the hybrid PMMA/PDMS design approach^17^ (Supplementary TABLE. S.1). The migration directedness, speed, and morphology of glioblastoma cells (see details in section IV.D) are quantitatively analyzed by a machine learning-based single cell segmentation and tracking software from stain-free phase contrast microscopy^30^. The cell morphologies of the two cell lines on various ECMs are shown in Supplementary FIG. S.2.

While standard poly(D-lysine) (PDL) and various combinations of poly(L-ornithine) (PLO) and laminin have been used for glioblastoma electrotaxis^16,39^, the adhesion and electrotaxis of T98G and U-251MG are not al-ways are not always consistent and reproducible as shown in Supplementary FIG. S.2 and FIG. S.3. T98G and U-251MG electrotactic responses are also not stably reproduced on collagen I, collagen IV, vitronectin, and fibronectin coatings.

Both T98G and U-251MG cells adhere well and demonstrate lamellipodia structures on substrates containing laminins, such as pure laminin coating, or Geltrex™. Geltrex™ is a growth factor-reduced complex basement membrane extract purified from murine Engelbreth-Holm-Swarm tumors containing laminin, collagen IV, entactin, and heparin sulfate proteoglycan^40^. Cells interact with laminins through various integrins including *α*1*β*1, *α*2*β*1, *α*3*β*1, *α*6*β*1, *α*7*β*1^41,42^. The integrins are believed to participate in the initiation of electrotaxis through mechanosensitive pathways^43–45^.

Discussed in details in section C, under electrical stimulation, the electrotaxis of both T98G and U-251MG are more prominent and reproducible on Geltrex™ coatings, hence all the studies in the following sections are based on Geltrex™ coatings. The detailed data of T98G and U-251MG electrotaxis on various ECMs is shown in Supplementary TABLE S.2.

An interesting observation is found in U-251MG cells on iMatrix-511-coated substrates. U-251MG cells demonstrate large lamellipodia associated with high migratory speed (15.45 *μ*m hr^−1^ under 300 V m^−1^, P < .0001) but with diminished directedness (0.01, P < .0001). Note that iMatrix-511 is a recombinant truncated laminin with *α*5*β*1*γ*1 subunits and interacts with cells through the *α*6*β*1 integrin^46,47^. This suggests that the specific molecular configuration of laminins in ECM may be vital for electrotaxis.

While it is understood that ECMs in tumor microenvironment are important^48^, glioblastoma cells demonstrate preference for adherence and electrotaxis on laminin-coated surfaces. Within the brain microenvironment, laminin expressions are restricted to the basement membrane of neural vasculature^49^ and perivascular tumor microenvironment is especially vital for glioblastoma metastasis^10,50^. Therefore, the correlation among laminin, glioblastoma electrotaxis, and the perivascular invasion process may be important in glioblastoma cancer biology that requires further elucidation.

### C. Electrotaxis behavior may reflect the heterogeneity of glioblastoma

To further analyze how T98G and U-251MG cells migrate under dcEF stimulation, the glioblastoma electrotaxis in serum-containing (FBS) and serum-free media were examined.

#### a. T98G and U-251MG cells migrate toward opposite directions under dcEF stimulation

FIG. 2 shows the directedness and speed of T98G and U-251MG electrotaxis. Upon 300 V m^−1^ dcEF stimulation, T98G cell migrate toward anode (positive electrode) while the U-251MG migrate toward cathode (negative electrode). Furthermore, the directedness in the electrotaxis of T98G cells does not depend on the presence of fetal bovine serum (FBS) (P = .45) but the speed is significantly decreased (P < .0001). The directedness of U-251MG cell electro-taxis however is highly dependent on the presence of FBS (P < .0001) and lack of FBS also decreases the speed of U-251MG electrotaxis (P < .0001). The FBS serum is rich in growth factors, proteins, and ions, which can enhance the chemical signaling in electrotaxis (Supplementary FIG. S.1). The electrotaxis and random migration of T98G and U-251MG with or without electrical stimulation is shown in Supplementary VIDEO. S.1 – Supplementary VIDEO. S.4.

**FIG. 2.**
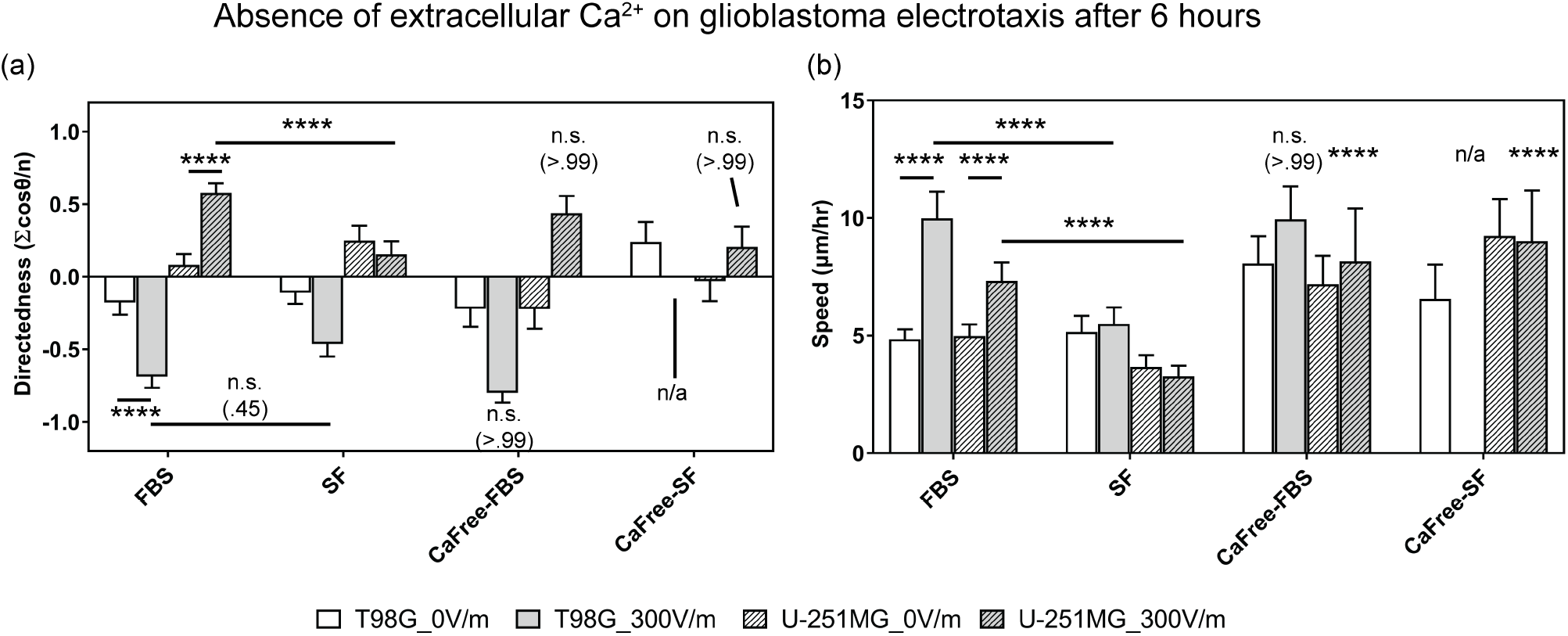
The electrotaxis of T98G & U-251MG glioblastoma cells in dependence of the extracellular serum and calcium by varying the medium recipe. (a) The electrotaxis directedness; (b) The electrotaxis speed; n.s. indicates not significant; **** indicate P < .0001; The numbers in parentheses indicate actual P-values. FBS: DMEM with serum; SF: serum-free DMEM; CaFree-FBS: 0 mM calcium DMEM with 10% FBS; CaFree-SF: 0 mM calcium serum-free DMEM.

Both T98G cells and U-251MG cells are categorized as mesenchymal type glioblastoma^37,38^, however, their electrotactic responses are completely different. Similar results are reported in the electrotaxis of glioblastoma cells and spheroids^15,16,19,39^ and lung adenocarcinoma^51^, showing that although molecular and surface marker makeups of the cell lines are similar, their electrotaxis responses can be completely different. The opposite electrotaxis results may reflect the fundamental heterogeneity among glioblastoma cells which has been speculated to contribute to the recurrence and therapeutic resistance after anti-tumor therapy^52–56^. Further elucidation of the correlation among electrotactic responses, metastatic properties of glioblastoma, and *in situ* electric field around the lesion may be beneficial to evaluate electrotaxis response as a predictive tool for glioblastoma metastasis.

#### b. Only T98G cells demonstrate prominent electroalignment behavior under electrical stimulation

Aside from directional migration in the dcEF, cells may also demonstrate long axial alignment in perpendicular to the dcEF vector. While this phenomenon is commonly observed among many cell types^51,57–60^, the molecular mechanism and the biological role are not clear. Electro-alignment may participate in the cytoskeletal restructuring in tissue morphogenesis^61,62^, but biophysical studies show that *in vitro* microtubules align in parallel to electric field vectors rather than perpendicular^63,64^.

Supplementary FIG. S.4 shows orientation of the cells with respect to time in electrically stimulated T98G and U-251MG cells over 6 hours. The orientation index is defined as the average cosine of two times the angle between the long axis of a cell and the electric field vector (See more details in Supplementary FIG. S.11 & Supplementary TABLE. S.5). For a group of perpendicularly oriented cells, the average orientation index is −1 and for a group of parallely orientated cells, the average orientation index is 1. For a group of randomly arranged cells, the average orientation index should be 0. Only T98G cells show prominent perpendicular alignment after electrical stimulation (Index_orientation_ at 0 hr v.s. 6 hr is −0.11 v.s. −0.83, P < .0001). FBS deprivation slightly decreases the alignment tendency and delays the onset but does not abolish it (Index_orientation_ of serum free v.s. 10% FBS at 6 hr is −0.48 v.s. −0.83, P < .0001). However, U-251MG does not show any perpendicular alignment. These results further illustrate the heterogeneity of glioblastoma cells. The cell electroalignment phenotypes of T98G and U-251MG after dcEF stimulation are shown in Supplementary FIG. S.5.

### D. Glioblastoma electrotaxis requires extracellular calcium

Calcium ion flux is known to be involved in the electrotaxis signaling of multiple cell types including mouse fibroblasts, human prostate cancer cell, and neural crest cells^65–69^. Deregulation of calcium influx in the cells reduces actin polymerization and affects cell motility speed but its effect on electrotactic directedness vary depending on cell types^65,69^. The hypothesis that glioblastoma electrotaxis is dependent of extracellular calcium cations is tested in this work (FIG. 2).

First, a calcium-free, serum-free cell culture media (CaFree SF) is used to test if glioblastoma electrotaxis requires extracellular calcium. T98G cells lose viability both with and without dcEF stimulation. Lack of calcium ion in cell culture media may impact calcium homeostasis significantly but the loss of viability is rescued by the addition of 10% FBS which contains calcium and growth factors. The electrotaxis of U-251MG in calcium-free, serum-free media is not affected compared to those in serum-free media (P > .99). Interestingly, the electrotactic speed of U-251MG in calcium-free serum-free media further increases (P < .0001).

Second, to validate that calcium cations are important, cation chelators EDTA and EGTA are used to chelate the free calcium in the cell culture media with 10% FBS. Treatment of 2 mM EDTA significantly represses the directedness of only U-251MG cells (P < .0001, Supplementary FIG. S.6.a). To further confirm the electrotaxis inhibition is due to extracellular calcium cations, EGTA, a divalent cation chelator with increased affinity towards Ca^2+^, is used. At 1 mM EGTA, the electrotactic directednesses of neither cell lines are affected but the electrotactic speeds of them become more dispersed (P < .0001). Under 2 mM EGTA treatments, the electrotactic directedness of U-251MG cells are reduced (P < .0001). When 5 mM EGTA is used, the electrotaxis of both cell lines are further repressed in both directedness and speed and the cells detach from the substrate. These results suggest that glioblastoma electrotaxis requires extracellular calcium cations and calcium influx may be important for electrotaxis, particularly that of U-251MG cells.

### E. Glioblastoma electrotaxis is mediated by voltage-gated ion channels

Ions channels expressed on glioblastoma cells including various potassium, calcium, sodium, and chloride ion channels are believed to facilitate pathogenesis of glioblastoma^27,70–72^. Although the expressions of numerous ion channels vary among clinical glioma samples^25^, ion channel expression profiles have been suggested to predict survival in glioma patients^73,74^. Glioblastoma cells are immersed in the local field potentials within the brain, and extracellular calcium is required for glioblastoma electrotaxis which can flow into the cells through voltage-gated calcium channels (VGCCs). Whether and how VGCCs participate in the electrotaxis of glioblastoma may shed new insights for inhibiting glioblastoma infiltration.

VGCCs are known to play important roles in glioma biology such as cell proliferation, apoptosis, and sensitization to ionizing radiation^27,75,76^. VGCCs can be categorized as high voltage activated (HVA) or low voltage activated (LVA) types^77–79^. Among the HVA VGCCs, four subtypes can be categorized by electrophysiological property and genetic phylogeny, including L-type (long-lasting, Ca_v_1.1–1.4), P/Q-type (purkinje/unknown, Ca_v_2.1), N-type (neural, Ca_v_2.2), and R-type (residual, Ca_v_2.3) VGCCs. The LVA VGCCs are composed of T-type channels (transient, Ca_v_3.1–3.3). Although the involvement of VGCCs in the electrotaxis of glioblastoma is not yet elucidated, VGCCs’ role in electrotaxis of other cell types has been proposed. Trollinger *et al.*, find that a stretch-sensitive VGCC mediates the electrotaxis of human keratinocytes^80^. Aonuma *et al.*, report that T-type VGCC mediates the electrotaxis of green paramecia^81^. L-type VGCCs also regulate the chondrogenesis during early limb development which is known to be a bioelectricity process^82^.

Another class of membrane proteins that are bioelectrically activated are voltage-gated potassium channels (VGKCs), which are represented by 12 families (K_v_1–K_v_12). VGKCs are involved in diverse physiological and pathological processess regulating the re-polarization of neuromuscular action potential, calcium homeostasis, cellular proliferation, migration, and cancer proliferation^83–89^. Voltage-gated potassium channel K_v_1.2^90^ and non voltage-gated inwardly-rectifying potassium channel K_ir_4.2^91^ have been shown to be involved in the sensing of electric field and signaling of cell electrotaxis. The potassium ion transporters confer biophysical signals that are key for regulating stem cells and tumor cells behavior in microenvironment^92^. In prostate cancer cells, VGKC expressions are linked to the increased metastatic potential^93^. Furthermore, inhibition of K_v_1.3 VGKC has been shown to induce apoptosis of glioblastoma cells *in vitro*^94^.

The role of VGCCs in glioblastoma cell electrotaxis is investigated by using pharmacological inhibitors (Supplementary TABLE. S.3). Many of these inhibitors are short peptides purified or recombinant engineered from venoms of poisonous species. When testing inhibition, cytochrome C was added to the culture media to prevent non-specific adsorption of peptide inhibitors to microfluidic chips^95^. The detailed results of pharmacological VGCC inhibition on the electrotaxis of T98G and U-251MG cells are shown in FIG. 3, Supplementary FIG. S.7, Supplementary FIG. S.8, and Supplementary TABLE. S.4.

**FIG. 3.**
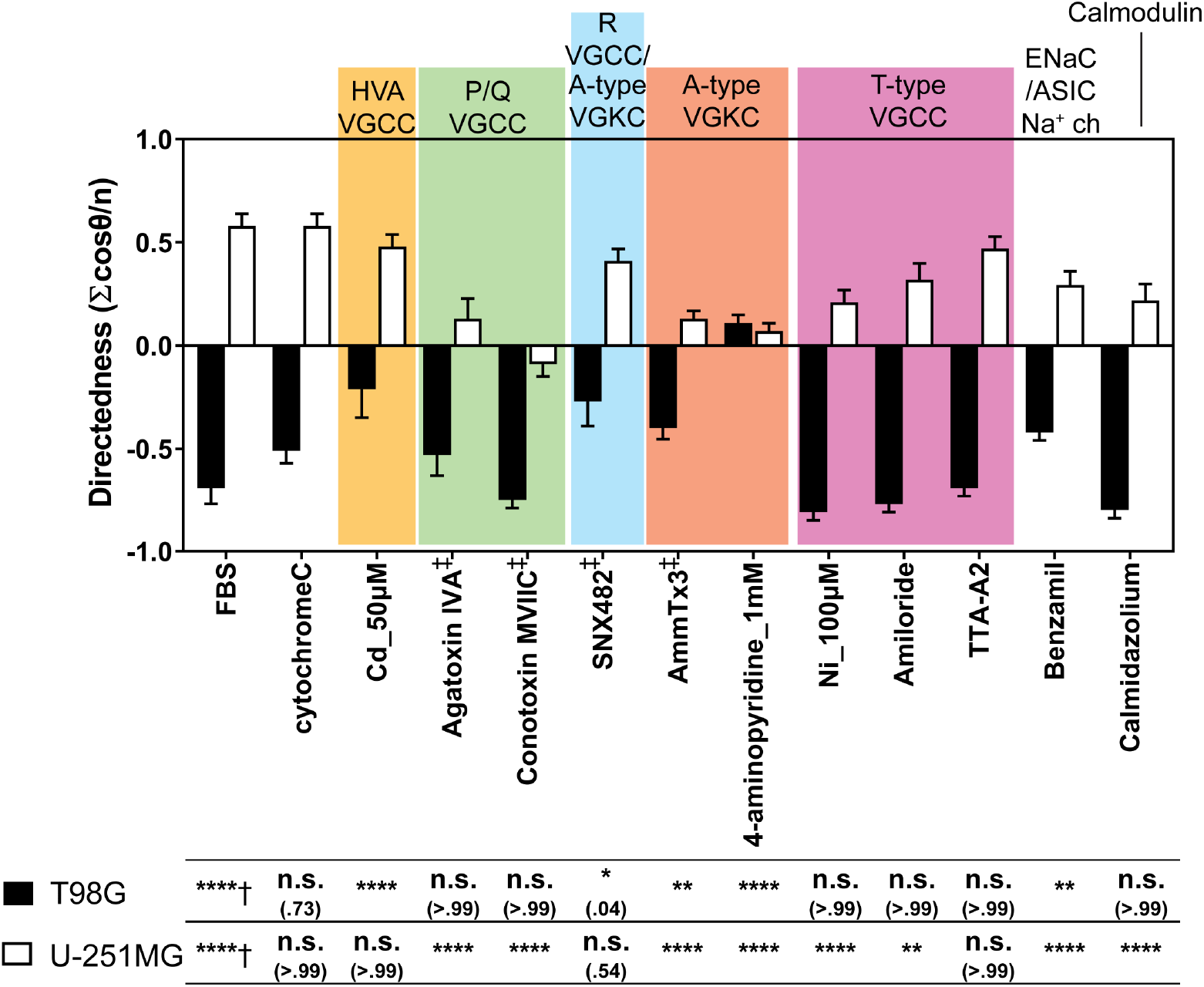
The electrotactic directedness of T98G & U-251MG glioblastoma cells under 300 V m^−1^ dcEF after 6 hours with pharmacological inhibition on various ion channels. † indicates the electrotaxis tested against those without EF stimulation; ‡ indicate the electrotaxis group tested against those with cytochrome C which prevents adsorption of short peptides to experimental apparatus; All other groups are statistically compared to their respective controls in cell culture media with 10% FBS; n.s. indicates not significant; * indicates P < .05; ** indicate P < .01; *** indicate P < .001; **** indicate P < .0001; The numbers in parentheses indicate actual P-values.

In both glioblastoma cell lines, inhibition of L-type HVA VGCC with gadolinium^96^ or nicardipine^97^ exhibit no effect on electrotactic directedness (P > .99) nor on speed (P > .15) (Supplementary FIG. S.7). Inhibition of N-type HVA VGCCs with *ω*-Conotoxin GVIA^98,99^ also has no effect on the electrotaxis of either cell type (P > .92).

#### a. T98G electrotaxis is mediated by R-type HVA VGCC

The electrotaxis of T98G is repressed when treated with cadmium which is a broad spectrum HVA VGCC inhibitor at 50 *μ*M and 100 *μ*M (P < .0001, FIG. 3)^100,101^. Upon further identification, the directedness in T98G electrotaxis is repressed by use of SNX-482, an R-type VGCC inhibitor^102,103^ (P = .049). The electrotaxis of T98G cells repressed with SNX-482 is shown in Supplementary VIDEO. S.5.

Calmodulin, a calcium binding protein, mediates many of the Ca^2+^ dependent-signaling by interacting with VGCCs and maintaining intracellular calcium homeostasis^104,105^. However, the electrotaxis of T98G is not dependent on calmodulin by inhibition with calmidazolium (P > .99, FIG. 3) and Ni^2+^ treatment has no inhibition on T98G cells (P > .99) which has partial inhibition on R-type VGCC^103,106^. These results imply an alternative mechanism might be at play.

#### b. U-251MG electrotaxis is mediated by P/Q-type HVA VGCCs

U-251MG electrotaxis has exhibited its dependency on HVA VGCCs. Decreased directedness and speed are observed in U-251MG cells treated with 100 *μ*M cadmium (P = .0372, Supplementary TABLE S.4). Upon further identification, U-251MG electrotaxis directedness is repressed when treated with P/Q-type HVA VGCC inhibitor using agatoxin IVA and conotoxin MVIIC (P < .0001) (FIG. 3). However, the electrotactic speed is not affected by agatoxin IVA (P > .99) but decreased by conotoxin MVIIC (P < .0001) (Supplementary FIG. S.8). The electrotactic directedness of U-251MG is dependent on calmodulin (P < .0001). The electrotaxis of U-251MG is repressed by the treatment of agatoxin IVA, shown in Supplementary VIDEO. S.6.

Furthermore, the electrotactic directedness of U251MG cells is repressed with nickel (P < .0001) and amiloride (P < .01) as well as the electrotactic speed (P < .0001) (FIG. 3). These results suggest a possible involvement of T-type VGCC^107^ in U-251MG electrotaxis. However, U-251MG electrotaxis is not affected when tested using another potent T-type VGCC inhibitor, TTA-A2^108,109^.

#### c. T98G and U-251MG electrotaxis are also mediated by A-type VGKCs and acid-sensing sodium channels

In T98G electrotaxis, cadmium and SNX-482 inhibit the Rtype HVA VGCC and decreases electrotaxis directedness. However, cadmium and SNX-482 have been reported to also block a rapid inactivating (A-type) transient outward VGKC (K_v_4.3) and experimental results of SNX482 should be interpreted carefully^110–112^. Using 5 *μ*M AmmTx3, a member of the *α*-KTX15 family of scorpion toxins, to block A-type VGKCs (K_v_4.2 & K_v_4.3)^113–115^, T98G electrotaxis directedness (FBS v.s. AmmTx3 = −0.69 v.s. −0.39, P = .0025) and speed (FBS v.s. AmmTx3 = 9.99 v.s. 4.42, P < .0001) are repressed but not completely abolished (Supplementary TABLE S.4). Furthermore the inhibition on T98G directeness caused by SNX482 is stronger than AmmTx3 (SNX-482 v.s. AmmTx3 = −0.27 v.s. −0.39, P = .039, two-tailed t test). Another broad spectrum transient VGKC inhibitor, 4aminopyridine (4-AP), was used to confirm the results from AmmTx3^116–118^. Under 1 mM 4-AP, the directedness and speed in T98G electrotaxis were both repressed (P < .0001, FIG. 3). Increasing of 4-AP to 4 mM, though results suggest that electrotactic directedness has not changed (P > .99) compared to control, but the migration speed is decreased (P < .0001). This is likely an artifact from part of T98G cells starting to detach from the surface rather than actual electrotaxis (Supplementary FIG. S.9). These results suggest that T98G electrotaxis may be mediated by A-type VGKCs (K_v_4.3), but the involvement of R-type VGCCs cannot be completely ruled out. Further molecular studies into the roles of VGCCs and VGKCs in glioblastoma electrotaxis are required.

Similar inhibition of A-type VGKC also represses U251MG electrotaxis. When U-251MG cells were treated with AmmTx3 and 4-AP, the electrotaxis directedness is inhibited by both compounds (P < .0001) and the speed is repressed in only 4-AP (P <.0001) but not AmmTx3 (P = .09) (Supplementary FIG. S.8). The electrotaxis of T98G and U-251MG is suppressed when A-type VGKC is inhibited through 1 mM 4-AP are shown in Supplementary VIDEO. S.7 & VIDEO. S.8).

Furthermore, nickel and amiloride repress the directedness of U-251MG electrotaxis but not through T-type VGCC. At high concentration of nickel and amiloride, the compounds may inhibit acid-sensing ion channel (ASIC) and epithelial sodium channel (ENaC), which are members of a superfamily of voltage-insensitive mechanosensitive sodium channels^119,120^. Furthermore, ASIC sodium channels are specifically expressed in the high-grade glioma cells but not in normal brain tissues or low grade glial cells^121,122^. Sodium ion flux is known to mediate electrotaxis in keratinocytes through ENaC sodium channels^123^ and prostate cancer cells through voltagegated sodium channels^124–126^.

To confirm the involvement of ASIC sodium channels in U-251MG electrotaxis, U-251MG cells are treated with 5 *μ*M benzamil hydrochloride (Alomone labs, USA)^127–129^. The directedness of U-251MG electrotaxis is significantly repressed by benzamil treatment (FBS v.s. benzamil = 0.58 v.s. 0.29, P < .0001, FIG. 3) but not the speed (P = .42) (more details in Supplementary FIG. S.8). Benzamil is also tested on T98G electrotaxis and found to inhibit its directedness (P < .01) and speed (P < .0001).

#### d. Various ion channels participate in the electrotaxis of glioblastoma cells of different origins

The pharmacological studies on the ion channels in T98G and U251MG electrotaxis suggest that multiple ion channels, which may be voltage-sensitive or not, can mediate the sensing of endogenous electric field and initiate the migratory response^13^. The proposed mechanism is highlighted in FIG. 4.

**FIG. 4.**
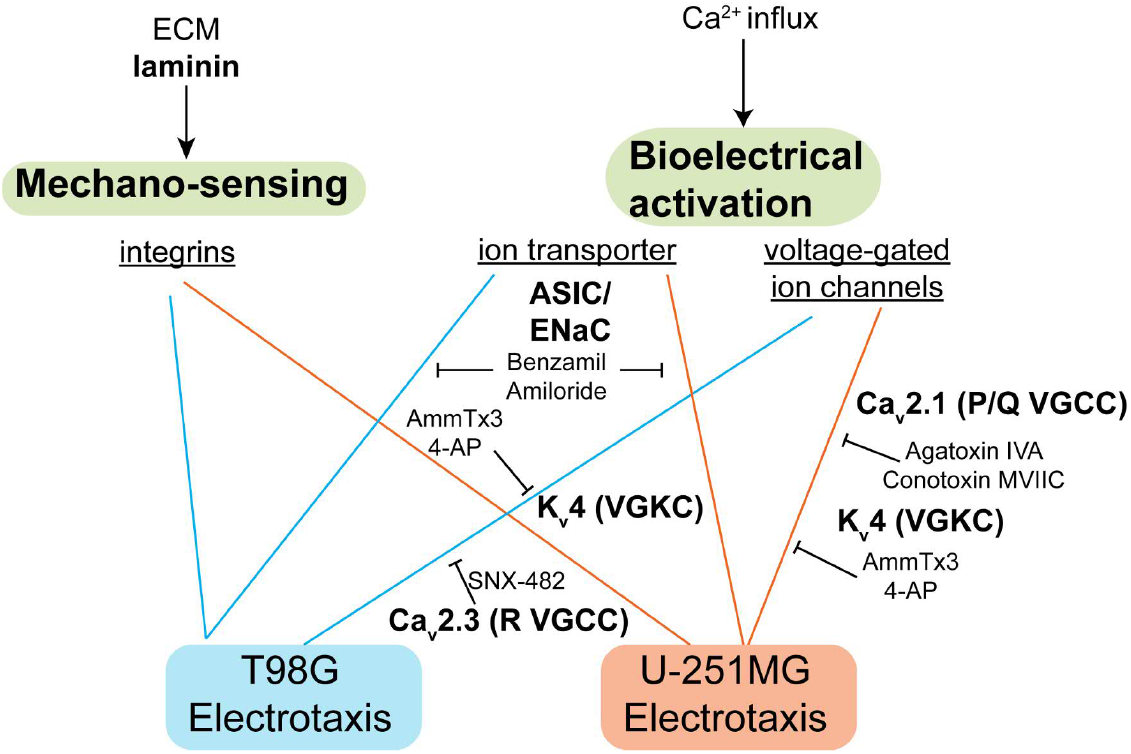
The signaling mechanism identified in this study. Laminin-based ECMs is necessary for glioblastoma electrotaxis, suggesting that integrins may play a role. The voltagegated ion channels and ion transporters also mediate glioblastoma electrotaxis that requires extracellular calcium.

A-type VGKC, R-type VGCC, and ASIC sodium channels mediate the electrotaxis of T98G cells while P/Qtype VGCCs, A-type VGKC, and ASIC sodium channels mediate the electrotaxis of U-251MG cells. These results suggest that ion channel expression profiles are cell line specific and correlating ion channel expressions with electrotactic phenotypes of cancer cells may be beneficial to provide new insights of metastasis-aimed therapeutics by inhibiting electrotaxis^130,131^. If glioblastoma infiltration can be inhibited by targeted therapeutics, the quality-of-life and prognosis of glioblastoma patients could be improved. The downstream molecular signaling of VGCCs, VGKCs, and ASIC sodium channels in glioblastoma electrotaxis is an interesting future direction to investigate. Recently, glutamatergic receptors have also been shown to mediate neuron-glioma interaction and glioma progression through calcium signaling^132–134^. Therefore, a systematic screening of potassium channel, sodium channels, and glutamate receptor ion channels’ ability to mediate glioblastoma electrotaxis is necessary to map the signaling network that may contribute to glioblastoma metastasis. The high-throughput hybrid microdevices and machine learning-assisted quantitative analysis developed in this work can be very useful for systematic phenotype profiling and identification of molecular mechanisms underlying cell electrotaxis.

## III CONCLUSION

The hybrid PMMA/PDMS microfluidic chip is a robust platform for high throughput electrotaxis studies. Cell migration in multiple dcEFs under multiplex conditions can be studied in a single experiment in combination with automated microscopy. The submerged operation balancing the inlet/outlet hydrostatic pressures guarantees a stable microenvironment that avoids microbubbles, ensures uniform cell seeding, and minimizes required number of cells. Uniformly distributed single cells can be reliably seeded in microfluidic chip that further increases the robustness for high throughput and more reproducible experiments. Use of machine learning-enabled single cell migration analysis automates the cell migration data analysis workflow for reliable quantitative data at high throughput.

Geltrex™ coating has been identified to support reproducible electrotaxis model of T98G and U-251MG glioblastoma cells. The heterogeneity responses of T98G and U-251MG electrotaxis and the importance of calcium signaling are identified. By further inhibitorial study, the electrotaxis of T98G may depend on R-type HVA VGCCs, A-type VGKCs, and ASIC sodium channels. The electrotaxis of U-251MG depends on P/Q-type HVA VGCCs, A-type VGKCS, and ASIC sodium channels. Multiple ion channels, which may be voltage-sensitive or not, can mediate the sensing of electric field and electrotaxis in different glioblastoma models, suggesting that glioblastoma infiltration can be amplified by endogenous electric field in a tumor sample-dependent manner. The roles of ion channels on glioma metastasis and survival with regard to physiological electric field require further systematic studies and *in vivo* validation.

As proof of principle, the hybrid PMMA/PDMS microfluidic design demonstrates robustness and versatility for high throughput experiments. The microfluidic chip design can be tailor-made for specific biological study. By using a robust, flexible, and high-throughput microfluidic platform together with machine learning software, the bottleneck of data analysis in high throughput experiments can be resolved, opening new opportunities for quantitatively studying cell responses in well-controlled microenvironments.

## IV. METHODS

### A. Hybrid multiple electric field device (HMEFC) design, simulation, and fabrication

To phenotypically test cell electrotaxis and elucidate molecular mechanism, a reliable *in vitro* platform for high throughput study is necessary. The HMEFC is designed by the hybrid PMMA/PDMS approach^17^ (FIG. 5.a). Using the hybrid PMMA/PDMS approach, the prototyping disadvantages in both materials can be mitigated while the advantages can be combined. In PMMA, complex 3D structures for fluidic routing or reservoir and worldto-chip interface can be quickly prototyped by CO_2_ laser cutting and thermal bonding. However, spatial resolution using this approach is not sufficiently high to create reliable microfluidic environments. In contrast, precise quasi-two dimensional microstructures can be fabricated using the soft lithography technique in PDMS microfluidic chips. But standard soft lithogarphy for PDMS fluidics is limited in the 3D design and worldto-chip interface. By using a dual-energy double-sided tape, PMMA and PDMS substrates can be easily and reversibly bonded^17,135^, enabling broad experimental flexibility.

**FIG. 5.**
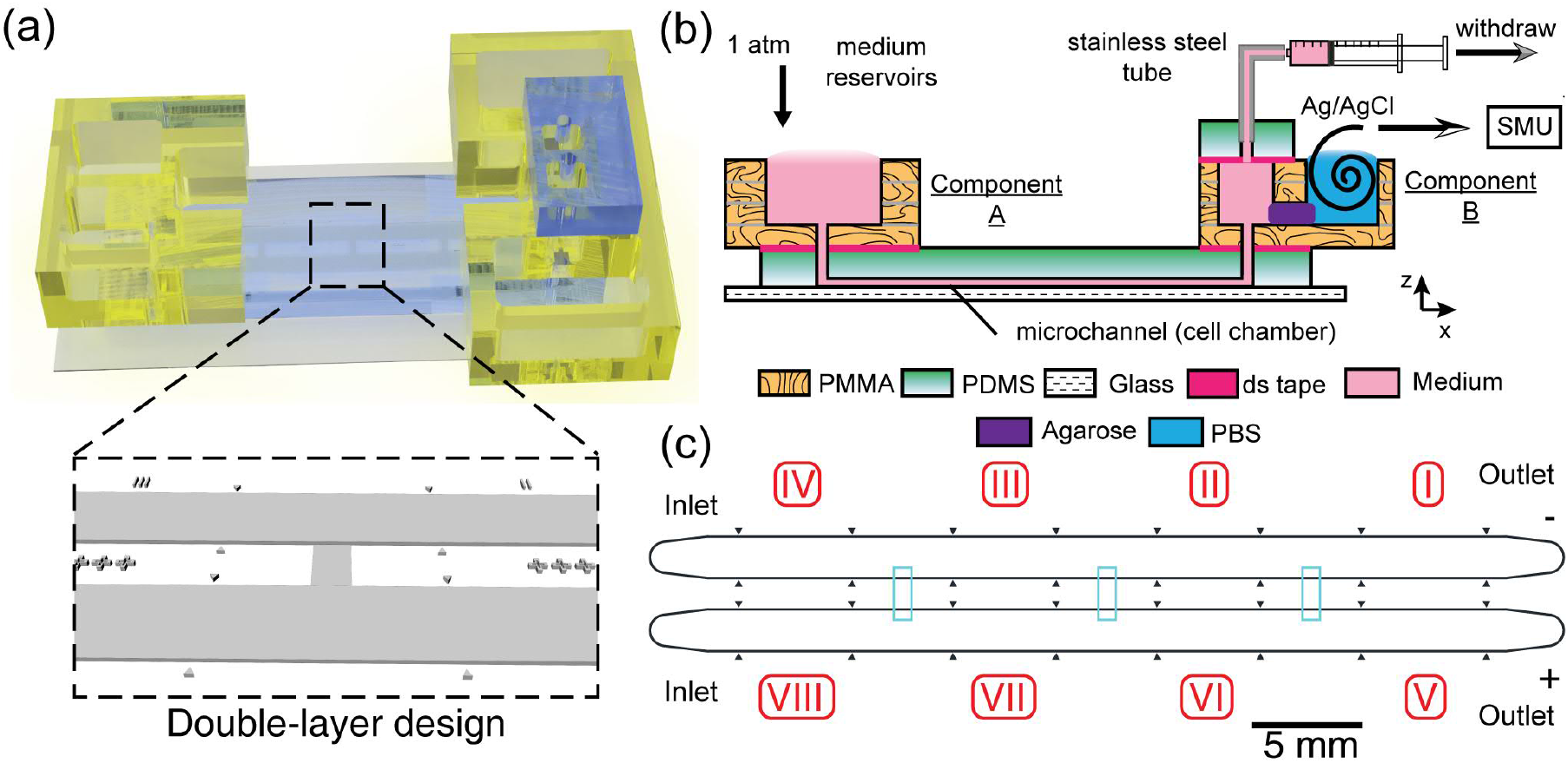
The design of hybrid multiple electric field chip (HMEFC). (a) The 3D rendered model of the final HMEFC. Double-layer microchannel design in the PDMS is shown; (b) The schematic diagram of using the HMEFC for electrotaxis experiments. Complex 3D microfluidic structures and world-to-chip interface are established in PMMA components A & B; (c) The channel design. The 10 *μ*m-high first layer structures are shown in cyan.

In HMEFC, two PMMA components for world-tochip interface and electrical application were adhered to a PDMS chip which contained a double-layer microchannel network where cells were cultured in and observed (FIG. 5.b). By double-layer microchannel design(FIG. 5.c), experiments with two different cell types or chemical treatments with four electric field strength (EFS) conditions could be performed in a single chip.

To create multiple dcEFs, the HMEFC was designed by R-2R resistor ladder configuration^17,51,136^ to create theoretical 5.25:2.5:1:0-ratio multiple EFs in section I to IV and section V to VIII (FIG. 5.c). The cells exposed to the highest dcEF were closest to the outlets to avoid paracrine signaling from electrically-stimulated cells to un-stimulated cells.

By using the double-layer design, the hydraulic resistances in the 10 *μ*m-high, 0.84 mm-wide interconnecting channels were much higher than the two 100 *μ*m-high, 2 mm-wide main channels where cells resided, limiting the advectional chemical transport and avoiding “cross-contamination” events that further increased the high experimental throughput (see detailed discussion on chip design in Supplementary Information).

Coupled numerical simulation of electric field, creeping flow, and chemical transport were carried out by finite element methods (COMSOL Multiphysics 5.3, COMSOL, USA). To correctly simulate the system, in-house measured material properties of the minimum essential media *α* (MEM*α*) supplemented with 10% FBS medium were measured and input in COMSOL. The liquid material properties of the 3D model was set as water with density of 1002.9 Kg m^3^, electrical conductivity of 1.536 S m^−1^, dynamic viscosity of 0.946 mPa s, and relative permittivity of 80. The numerically simulated electric field ratio was 4.99: 2.45:1:0 in section I to IV and section V to VIII with limited chemical transport across the interconnecting channels as designed (Supplementary FIG. S.13).

To fabricate HMEFC, first, the PDMS chip was fabricated by soft lithography technique^137^. The double-layer microfluidic design was fabricated into a mold using negative photoresists (10 *μ*m and 100 *μ*m) on a silicon wafer using direct-write lithographic writer and mask aligner (DL-1000, Nano Systems Solution, Japan and MA/BA6, SUSS MicroTec, Germany). After passiviation of the mold with perfluorosilane, mixed PDMS monomer (10:1 monomer:curing agent ratio, Sylgard 184, Corning, USA) was poured and cured on the master mold in a custom-made casting block which ensures the 4 mm thickness in finished PDMS devices. After degassing, a piece of 15 mm-thick PMMA block was placed on top of the casting block to ensure the surface flatness of PDMS. The PDMS was cured in oven and cut to yield individual devices. 1 mm-wide inlets and outlets were punched on cured PDMS devices and the PDMS devices were bonded to cover glasses (0.17 mm-thick) using O_2_ plasma, completing the PDMS chip.

Second, the PMMA components were fabricated by cutting the 4-layer design on a 2-mm thick casted PMMA substrate (Kanase, Japan) using a CO_2_ laser cutter (VLS3.50, Universal Laser Systems, USA). The layers were aligned and thermally bonded on a programmable automated hot press with temperature and pressure control (G-12RS2000, Orihara Industrial co., ltd, Japan). Third, dual energy double-sided tape (No.5302A, Nitto, Japan) was patterned using the CO_2_ laser cutter and used to join the PMMA components and the PDMS chip^17,135^. A PMMA top reservoir for submerged cell seeding was also fabricated by four layers of 2 mm-thick PMMA pieces using laser cutting and thermal bonding. The PMMA top reservoir was affixed on top of the two PMMA components through PDMS padding frames with dual energy double sided tapes, completing the assembly of HMEFC for cell seeding.

The detailed description for the HMEFC design, simulation, and fabrication can be found in the Supplementary Information.

### B. Cell culture and maintenance

Glioblastoma cell lines T98G (CRL-1690, ATCC, USA) and U-251MG (IFO50288, JCRB, Japan) were obtained from the respective tissue banks and thawed according to the received instructions. Ethics approval is not required. T98G and U-251MG were cultured in minimum essential media *α* (MEM*α*) supplemented with 10% FBS and 2.2 g L^−1^ under 37°C, 5% CO_2_ moist atmosphere (MCO-18AIC, Sanyo, Japan). The cells were subcultured every other day or whenever cells reached 80% confluency. Mycoplasma contamination was checked every three months using a mycoplasma species-specific PCR kit (e-Myco plus, iNtRON, Korea) on a thermocycler (C1000, Bio-Rad, USA).

Frozen stocks of the cells were prepared by resuspending 1 × 10^6^ log-phase cells in 1 mL CellBanker solution (Takara Bio, Japan) and cooled down in a freezing container (Mr. Frosty, Nunc, USA) in −80°C overnight. The frozen cells were then transferred into gaseous phase of liquid nitrogen for long term storage (Locator 6, Thermo Scientific, USA).

### C. Cell seeding and electrotaxis experiment

The cell experiment workflow included salt bridge preparation, priming of micsrochannels, coating substrate with ECM, seeding cells, assembly of world-to-chip interface, and electrotaxis experiment.

First, sterilized 1% molten agarose (Seakem LE agarose, Lonza, USA) dissolved in 1X phosphate buffered saline (PBS) was injected on the salt bridge junctions of the PMMA component B and allowed to gel (FIG. 5.b). The salt bridge served as a solid electroconductive separation in electrotaxis experiments between the cell culture media and electrode, avoiding formation of complex electrolysis products.

Second, 50 *μ*L 99.5% ethanol (Wako, Japan) loaded in 200 *μ*L pipet tips were used to wet the microchannels in the PDMS chip by capillary flow and gravity flow^17,138^. The microchannels were then washed with ultrapure water. The inlet/outlet ports were submerged under liquid solutions in all steps afterward to ensure bubble-free microchannels.

Third, 150 *μ*L of appropriate ECM solutions such as Geltrex™ was loaded in tips and inserted on one side of inlet/outlet ports. The ECM solutions were allowed to flow into chip passively and incubated for 1 hour at 37°C. After ECM coatings, the channels were washed once with PBS and MEM*α*. The top reservoir was then filled with MEM*α* until the inlets and outlets were under the same liquid level, balancing the inlet/outlet hydrostatic pressure.

Log-phase glioblastoma cells were washed with 1X PBS, trypsinized (TrypLE, Thermo Fisher Scientific, USA), counted on a benchtop flow cytometer (Muse counting and viability kit, Millipore, USA), centrifuged at 300×g for 5 min, and resuspended in MEM*α* media with 10% FBS at 10^6^ cells mL^−1^. Appropriate amount of cell suspension was injected into the microchannels using a 200 *μ*L micropipet. Due to the small volume in microchannels, only a minute amount of cell suspension was needed. The cells were allowed to adhere in the chip under 37°C, 5% CO_2_ moist atmosphere for 3 to 5 hours.

After cell seeding and adhesion, the top reservoir was removed for optical clarity. A piece of 4 mm-thick PDMS slab punched with two outlet holes (21G, Accu-punch MP, Syneo Corp, USA) was affixed to the top of component B through the first piece of patterned double-sided tape (Supplementary FIG. S.15.a), creating an air-tight seal.

To start electrotaxis experiment, fresh media were supplied in the reservoirs on component A of HMEFC. Two sets of tubings (06419-01, Cole Parmer, USA) with stainless tubes (21RW, New England Small Tubes, USA) on one ends and double Luer gel dispensing needles on the opposite ends (23G, Musashi, Japan) were used. The tubings were sterilized before priming with 1X PBS with 2.5 mL syringes (Terumo, Japan). The stainless tube ends were inserted into the 21G holes of the PDMS slab on the HMEFC. The syringes were mounted on a syringe pump (YSP-202, YMC, Japan) and set in withdrawal mode to perfuse the cells on HMEFC. The HMEFC was completed and ready for electrotaxis experiment (FIG. 5.b).

Two HMEFCs were prepared in one experiment that allowed 16 conditions to be screened simultaneously. However for data presentation, we only showed the ones with 300 V m^−1^ and 0 V m^−1^ dcEF. HMEFCs were affixed in a microscope on-stage incubator (WKSM, Tokaihit, Japan). A feedback thin-film K-type thermocouple was attached to the glass bottom of a HMEFC (60 *μ*mthick, Anbesmt, Japan) and regulate the incubator to maintain environmental temperature at 37°C.

A U-shaped PMMA salt bridge filled with 1% molten agarose in PBS was used to connect the two HMEFCs electrically by inserting into the ajoining PBS reservoirs on each HMEFC’s component B. 300 V m^−1^ dcEFs were established in section I and V (FIG. 5.c) through two home-made silver/silver chloride (Ag/AgCl) wire electrodes^51,139^ inserted in the PBS reservoirs on component B by a source measure unit (2410, Keithley, USA). Time lapse phase contrast images of each condition were taken on an automated phase contrast microscope (TiE, Nikon, Japan). The setup is shown in Supplementary FIG. S.10.

For electrotaxis of glioblastoma on different ECM coatings, different ECM coatings were coated in the microchannels and electrotaxis experiments were performed in MEM*α* with 10% FBS for 6 hours.

To test if the electrotaxis of gliblastoma cells require extracellular Ca^2+^, calcium-free Dulbecco’s minimum essential medium (CaFree DMEM, 21068, Gibco, USA) or cation chelators such as ethylenediaminetetraacetic acid (EDTA) (15575, Thermo Fisher Scientific, USA) and ethylene glycol-bis(2-aminoethylether)N,N,N’,N’-tetraacetic acid (EGTA) (08907-84, Nacalai, Japan) were used (Supplementary TABLE. S.3). Before dcEF stimulation, cells were incubated in CaFree DMEM or MEM*α* with 10% FBS mixed with calcium chelators at indicated molar concentrations for 2 hours at 20 *μ*L hr^−1^. Afterward a direct electric current was applied through Ag/AgCl wire electrodes in D-PBS buffer on the PMMA component B by the source measure unit and cell behaviors were observed for six hours.

To investigate if voltage-gated ion channels mediate the electrotaxis of glioblastoma cells, the inhibitors were added to fresh MEM*α* media with 10% FBS at appropriate working concentrations (Supplementary TABLE. S.3). The reagents were supplied in the reservoirs on PMMA component A of HMEFCs after cells were seeded. The inhibitor-containing media were infused into the channels at 20 *μ*L min^−1^ for 10 minutes before changing to 20 *μ*L hr^−1^ and incubated for 30 minutes. A direct electric current was applied through Ag/AgCl wire electrodes in PBS buffer on the PMMA component B by the source measure unit and cell behaviors were observed for six hours.

### D. Microscopy imaging and data processing

A Nikon Ti-E automated microscope with Perfect Focus System and motorized XYZ stage was used to perform all microscopy experiments. A 10X phase contrast objective and intermediate magnification of 1.5X were used for taking 10-minute interval phase contrast images with a scientific CMOS (sCMOS) camera with 2×2 binning (Orca Flash 4.0, Hamamatsu, Japan) in NIS Element AR software (Nikon, Japan). The spatial resolution at this setting was ≈ 0.87 *μ*m pixel^−1^. All experiments were performed in triplicate.

After each experiment, images were exported from NIS element software as tiff files and automatically organized by XY positions into folders using an in-house developed Python script. The cells in the raw images were segmented, tracked, and automatically analyzed using the *Usiigaci* software^30^. Only cells tracked throughout all the frames in one viewfield were analyzed. At least 100 cells in every condition were analyzed for cell-centric features such as the directedness, speed, and orientation changes before and after electric field or chemical stimulation (Supplementary TABLE. S.5).

Briefly, the definitions of key cell-centric features used to quantify cell electrotaxis (Supplementary FIG. S.11) were listed below:

- Directedness The directedness of cell electrotaxis was defined as the average cosine of Euclidean vector and EF vector, 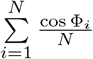, where Φ_i_ was the angle between the Euclidean vector of each cell migration and the vector of applied EF (from anode to cathode), and *N* was the total number of analyzed cells. A group of anodal moving cells held a directedness of −1; and a group of cathodal moving cells held a directedness of +1. For a group of randomly migrating cells, the directedness was zero.
- Speed The speed of cell electrotaxis was defined as the average of cell migration rate to travel the Euclidean distance, 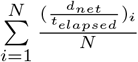, where the *d*_*net*_ was the Euclidean distance traveled by each cell, and *t_elapsed_* was the the time elapsed, and *N* was the total number of analyzed cells.
- Orientation The orientation was defined as the average cosine of two times angle between the EF vector and the cell long axis, 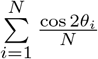, where *θ*_*i*_ was the angle between the applied EF vector and the long axis of a given cell; *N* is the total number of cells analyzed. A group of cells aligned perpendicular to the EF held an orientation of −1; and a group of cells aligned in parallel to the applied EF held an orientation of +1. For a group of randomly shaped cells, the average orientation was zero.

The cell-centric features were computed automatically and saved as Excel spreadsheets (Office365, Microsoft, USA) as part of the *Usiigaci* software analysis pipeline. The data was further statistically inferenced by inputting the data in a statistical software (Prism 7, GraphPad LLC, USA). All data were presented as the mean ± 95% confidence interval, which was 1.96 of standard error of mean, from triplicate experiments. Two-tailed Student’s t tests or one-way analyses of variance (ANOVA) with Tukey’s multiple-comparison post-hoc tests were performed for statistical testing between two groups or multiple groups. The results form one-way ANOVA were reported until noted otherwise. The confidence level to reject a null hypothesis between two datasets was set at 95%. A p-value (P, the probability for a true null hypothesis) less than 0.05 represented a statistical significance at 95% confidence.

## SUPPLEMENTARY MATERIAL

The supplementary figures, supplementary tables, supplementary videos, and detailed description of HMEFC chip design, simulation, and fabrication are collected in the supplementary material.

## DATA AVAILABILITY STATEMENT

The data that support the findings of this study are available from the corresponding author upon reasonable request.

## ACKNOWLEDGMENTS

H-.F. Tsai is a Research fellow of Japan Society for the Promotion of Science (JSPS Research Fellow) and this work is supported by JSPS KAKENHI (Grant Number JP1700362). The authors also thank Okinawa Institute of Science and Technology Graduate University (OIST) for its financial support with subsidy funding from the Cabinet Office, Government of Japan. Funders had no role in study design, data collection, the decision to publish, or preparation of the manuscript. The authors thank Professor Tomoyuki Takahashi (Cellular and Molecular Synaptic Function Unit, OIST) for his invaluable suggestion and discussion. The authors thank Ms. Yi-Ching Tsai for her assistance on illustration preparation.

H-.F. Tsai and A.Q. Shen declare potential conict of interest. A patent application has been submitted by Okinawa Institute of Science and Technology Graduate University based on these results with H-.F. Tsai and A.Q. Shen as co-inventors.

## Supplementary Figures

**FIG. S.1:**
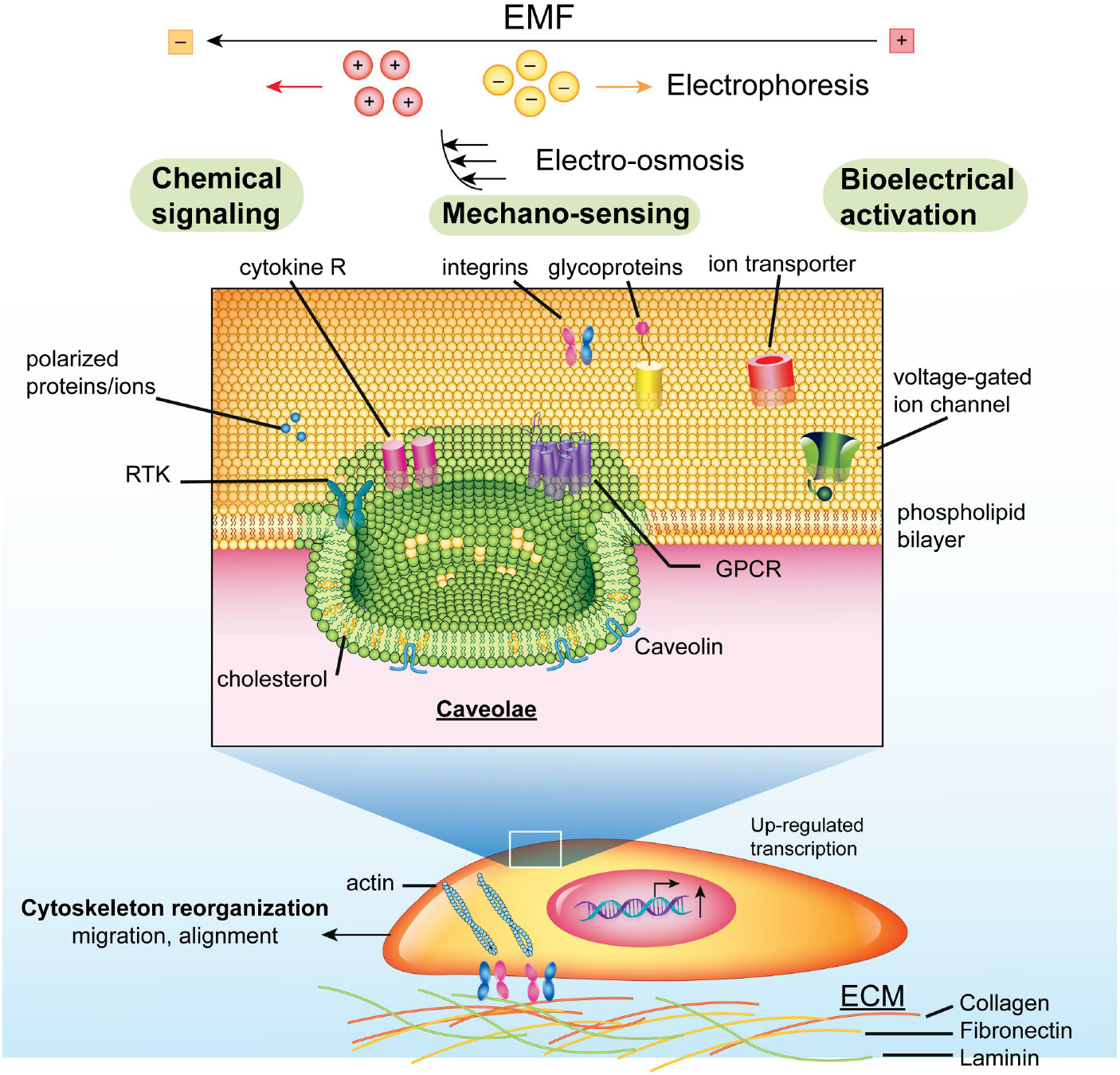
The complex mechanisms of cell response to electric fields. Adapted from McCaig et al. and Tsai et al^1,2^.

**FIG. S.2:**
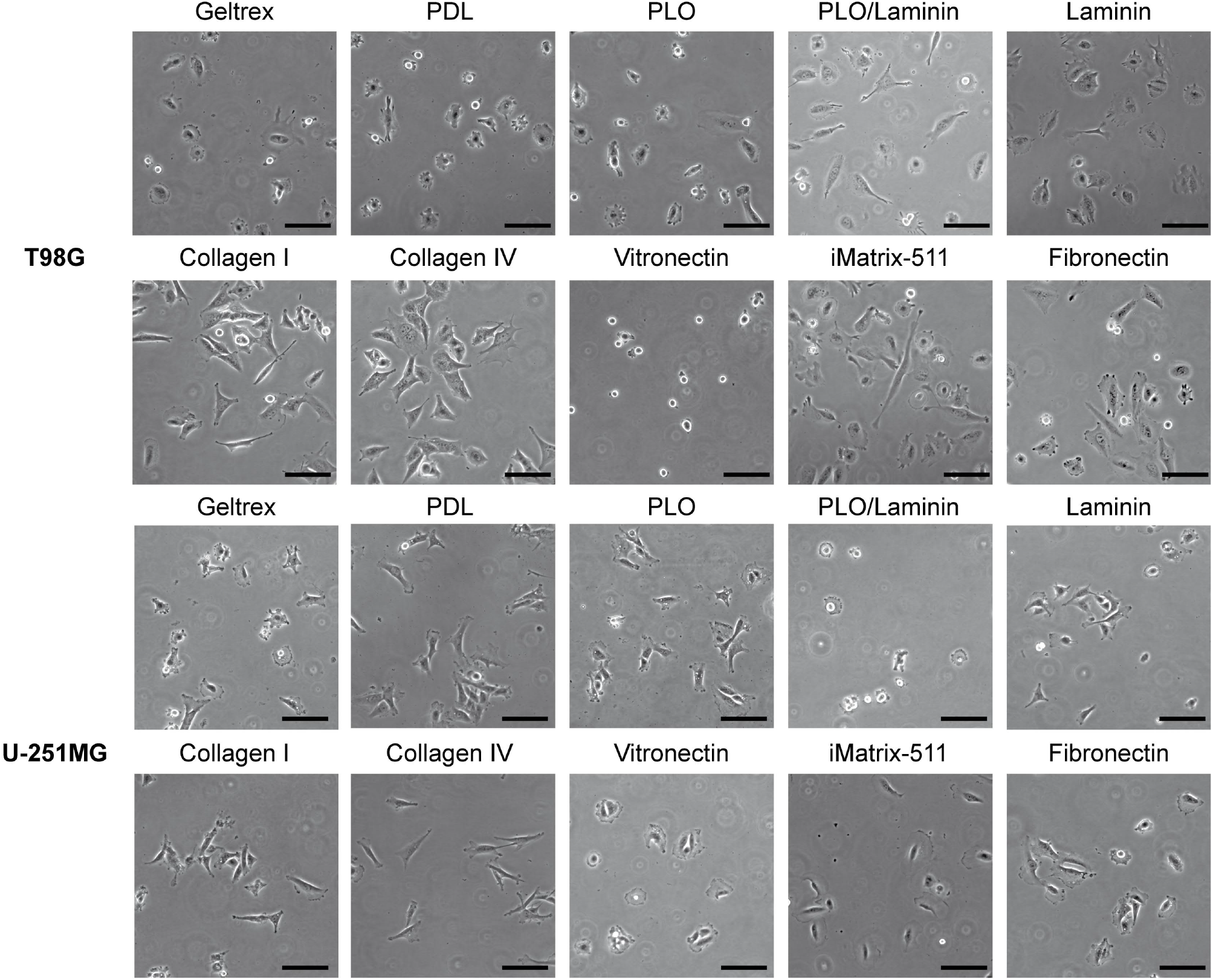
Phase contrast images of T98G & U-251MG cells on various ECMs. Scale bar: 100 *μ*m.

**FIG. S.3:**
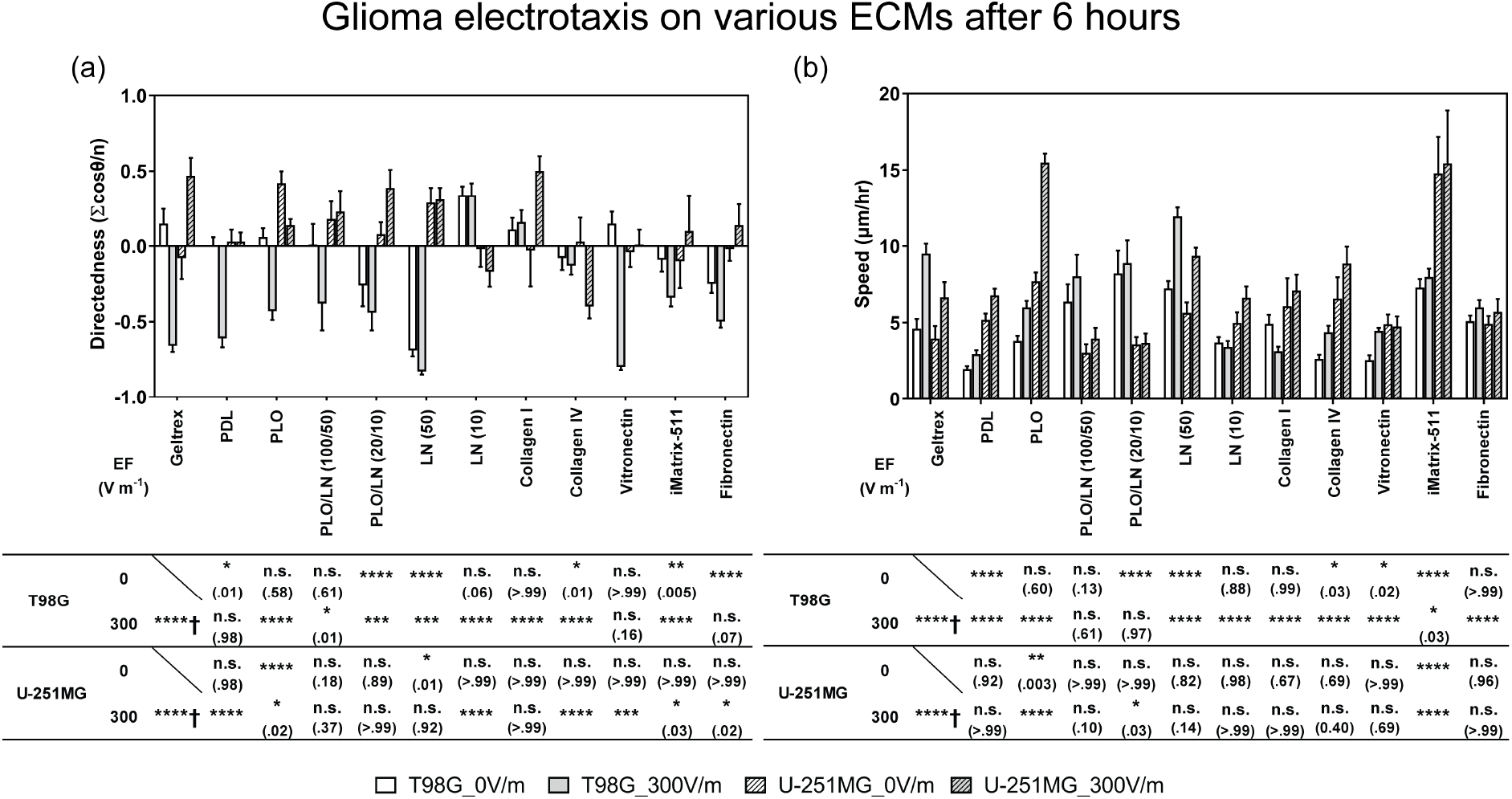
The electrotaxis of T98G & U-251MG glioblastoma cells on various ECMs after 6 hours under 300 V m ^−1^ EF stimulation. (a) The electrotaxis directedness; (b) The electrotaxis speed; † indicates the electrotaxis groups tested against those without EF stimulation; All other groups are statistically compared to their respective controls on Geltrex™ ECM; n.s. indicates not significant; * indicates P < .05; ** indicate P < .01; *** indicate P < .001; **** indicate P < .0001; The numbers in parentheses indicate actual P-values.

**FIG. S.4:**
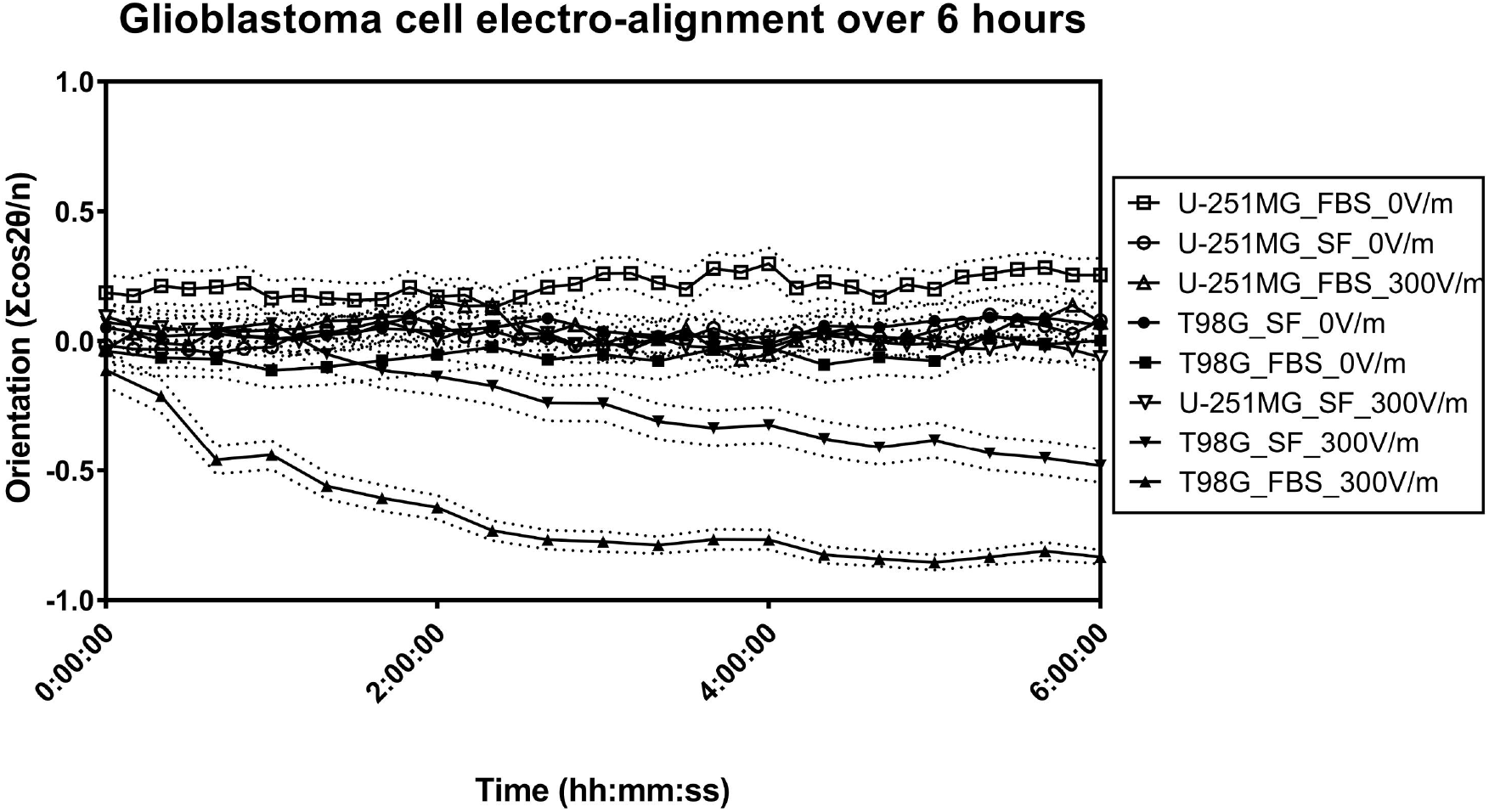
Time series plot of orientation in electrically stimulated glioblastoma cells. T98G cell results are indicated in closed symbols. U-251MG cell results are indicated in open symbols. Only T98G cells demonstrate prominent perpendicular alignment after electrical stimulation. The dashed lines indicate the 95% confidence interval for respective groups.

**FIG. S.5:**
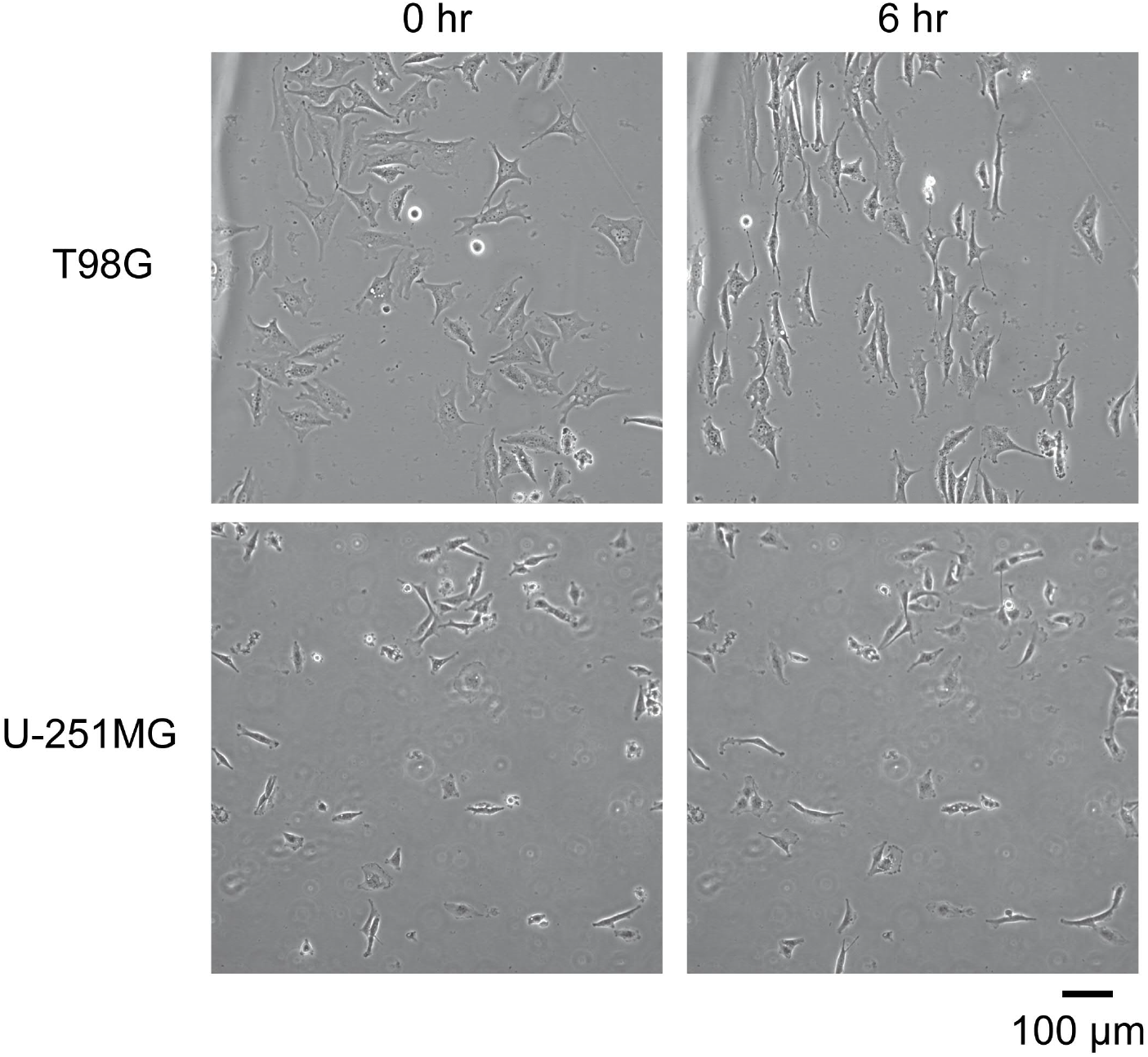
Phase contrast images of T98G and U-251MG cells before and after 6 hours 300 V m ^−1^ stimulation. The electric field is from left to right. Only T98G cells demonstrate prominent perpendicular allignment after electrical stimulation.

**FIG. S.6:**
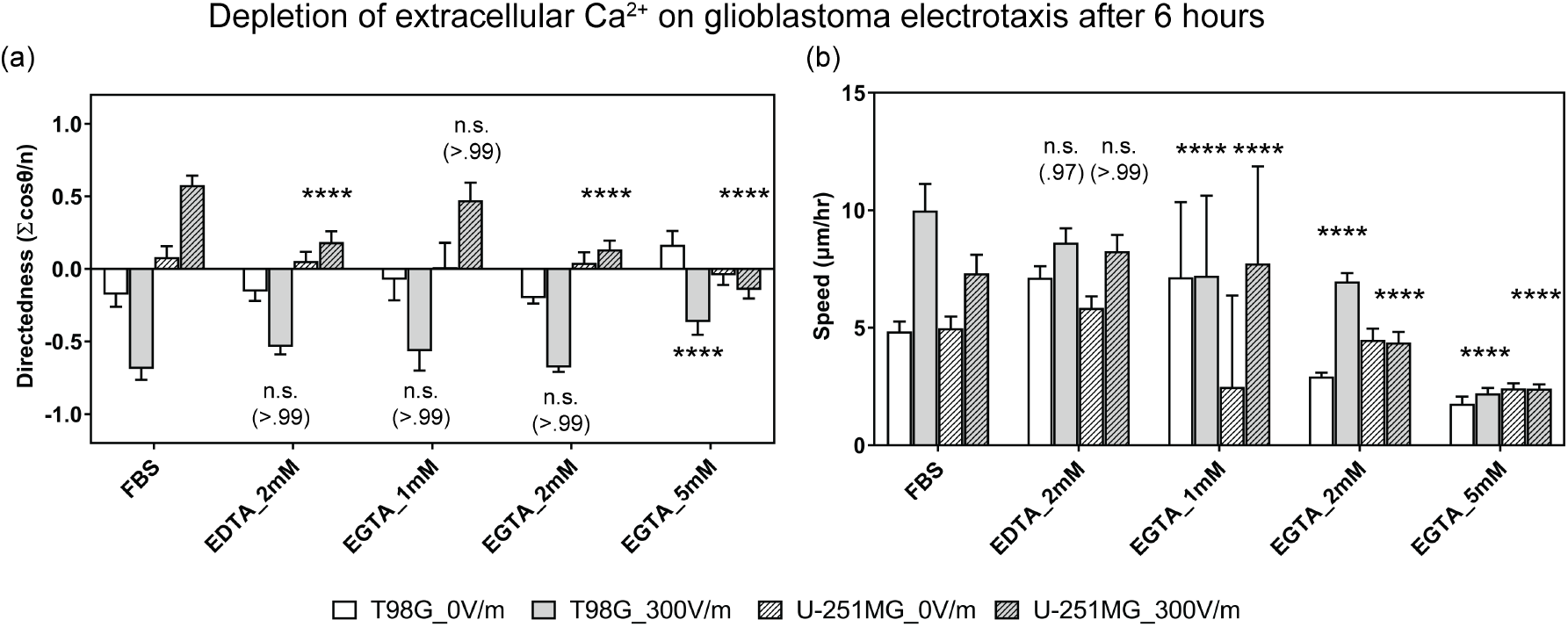
The electrotaxis of T98G & U-251MG glioblastoma cells in dependence of extracellular calcium by varying concentration of calcium chelators, EDTA and EGTA. (a) The electrotaxis directedness; (b) The electrotaxis speed; † indicate the electrotaxis groups tested against those without EF stimulation; All groups are statistically compared to controls in cell culture media with 10% FBS; n.s. indicates not significant; **** indicate P < .0001; The numbers in parentheses indicate actual P-values.

**FIG. S.7:**
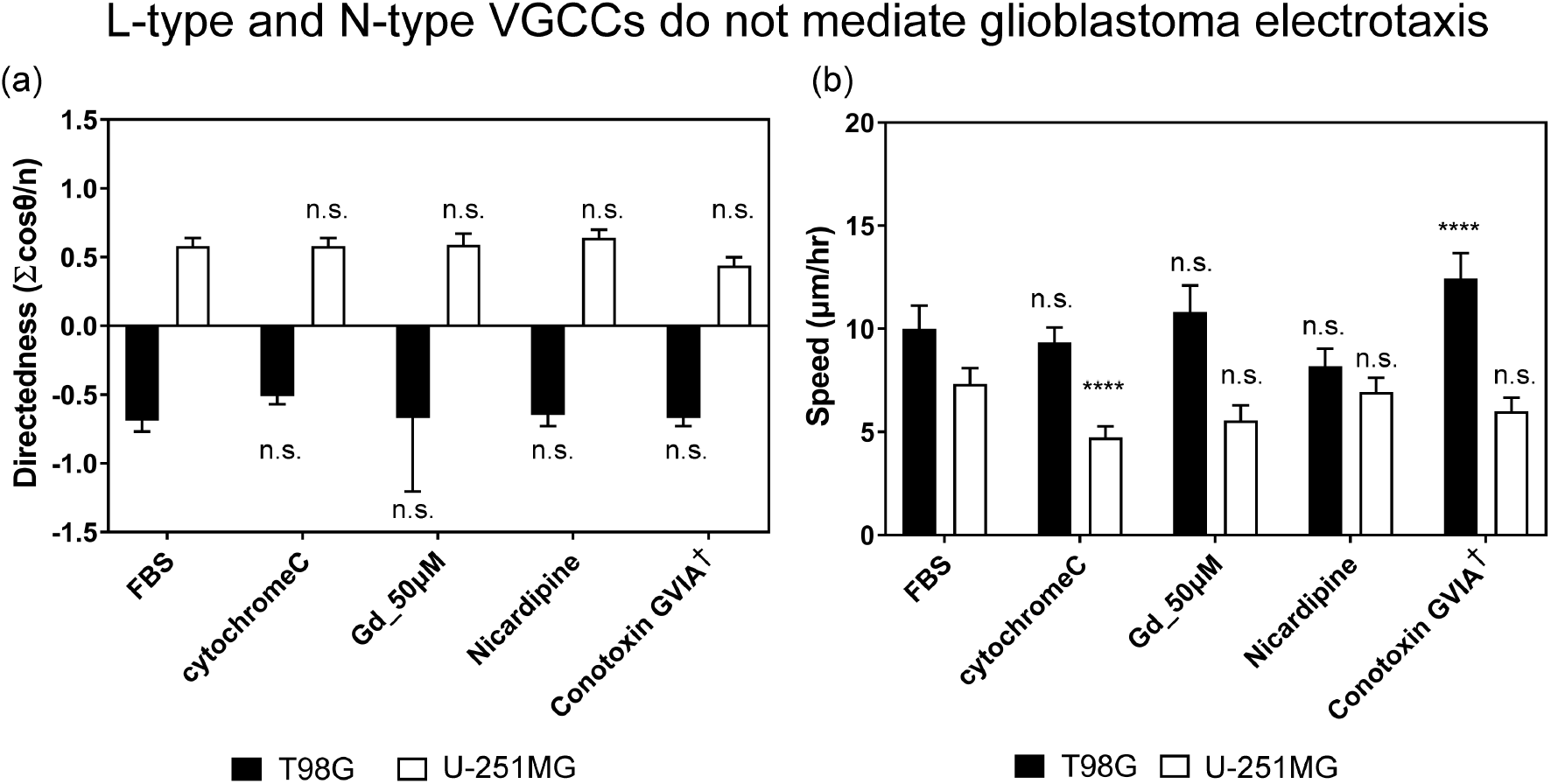
The electrotaxis of T98G & U-251MG glioblastoma cells under 300 V m ^−1^ dcEF after 6 hours with pharmacological inhibition on L-type (50 *μ*M Gadolinium [Gd] and Nicardipine) and N-type VGCCs (Conotoxin GVIA). a) The electrotactic directedness of the two cells. (b) The electrotactic speed of the two cells. ‡ indicate the electrotaxis group tested against those with cytochrome C which prevents adsorption of short peptides to experimental apparatus; All other groups are statistically compared to their respective controls in cell culture media with 10% FBS; n.s. indicates not significant; **** indicate P < .0001.

**FIG. S.8:**
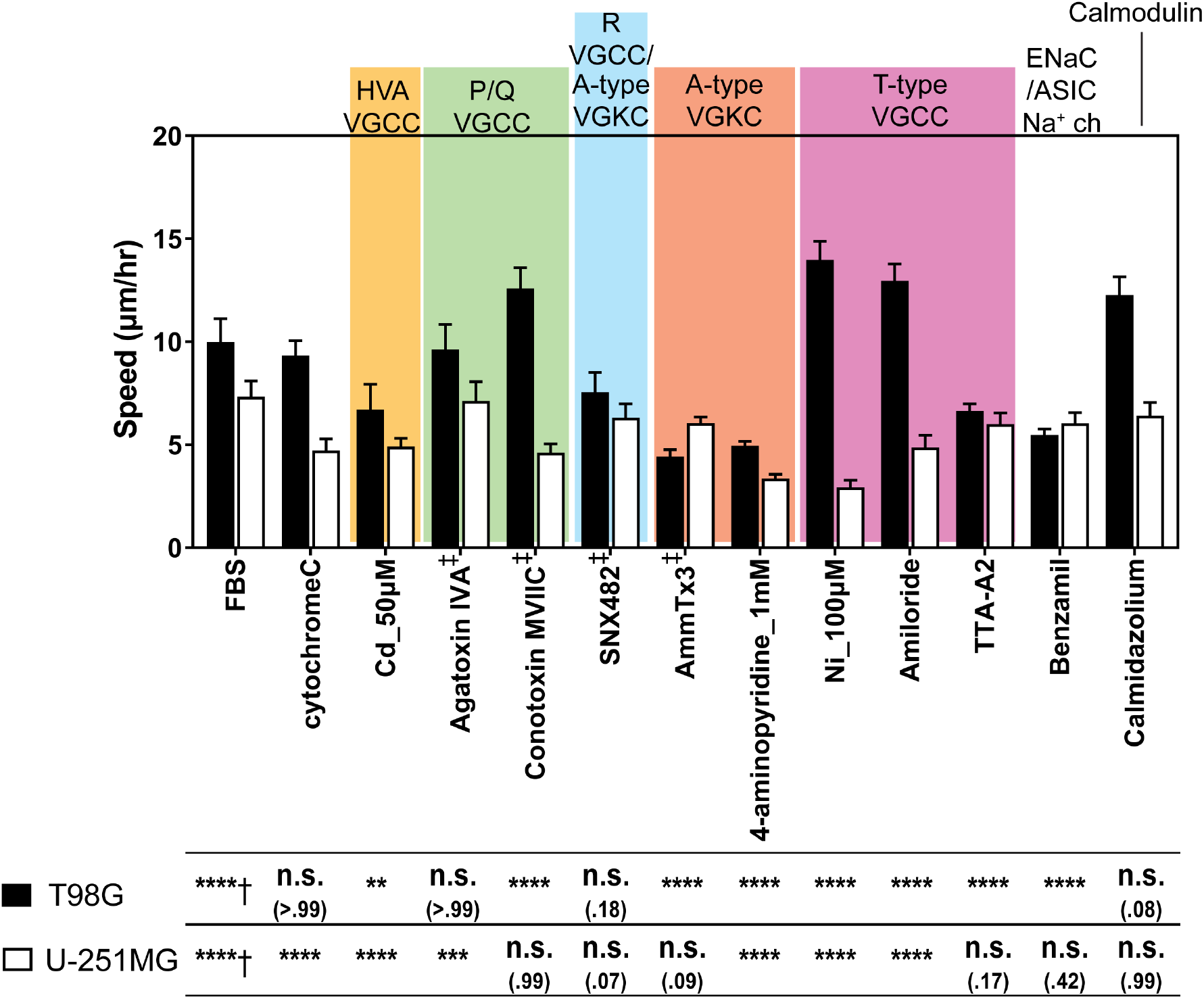
The electrotactic speed of T98G & U-251MG glioblastoma cells under 300 V m^−1^ dcEF after 6 hours with pharmacological inhibition on various ion channels. † indicates the electrotaxis tested against those without EF stimulation; † indicate the electrotaxis group tested against those with cytochrome C which prevents adsorption of short peptides to experimental apparatus; All other groups are statistically compared to their respective controls in cell culture media with 10% FBS; n.s. indicates not significant; ** indicate P < .01; *** indicate P < .001; **** indicate P < .0001; The numbers in parentheses indicate actual P-values.

**FIG. S.9:**
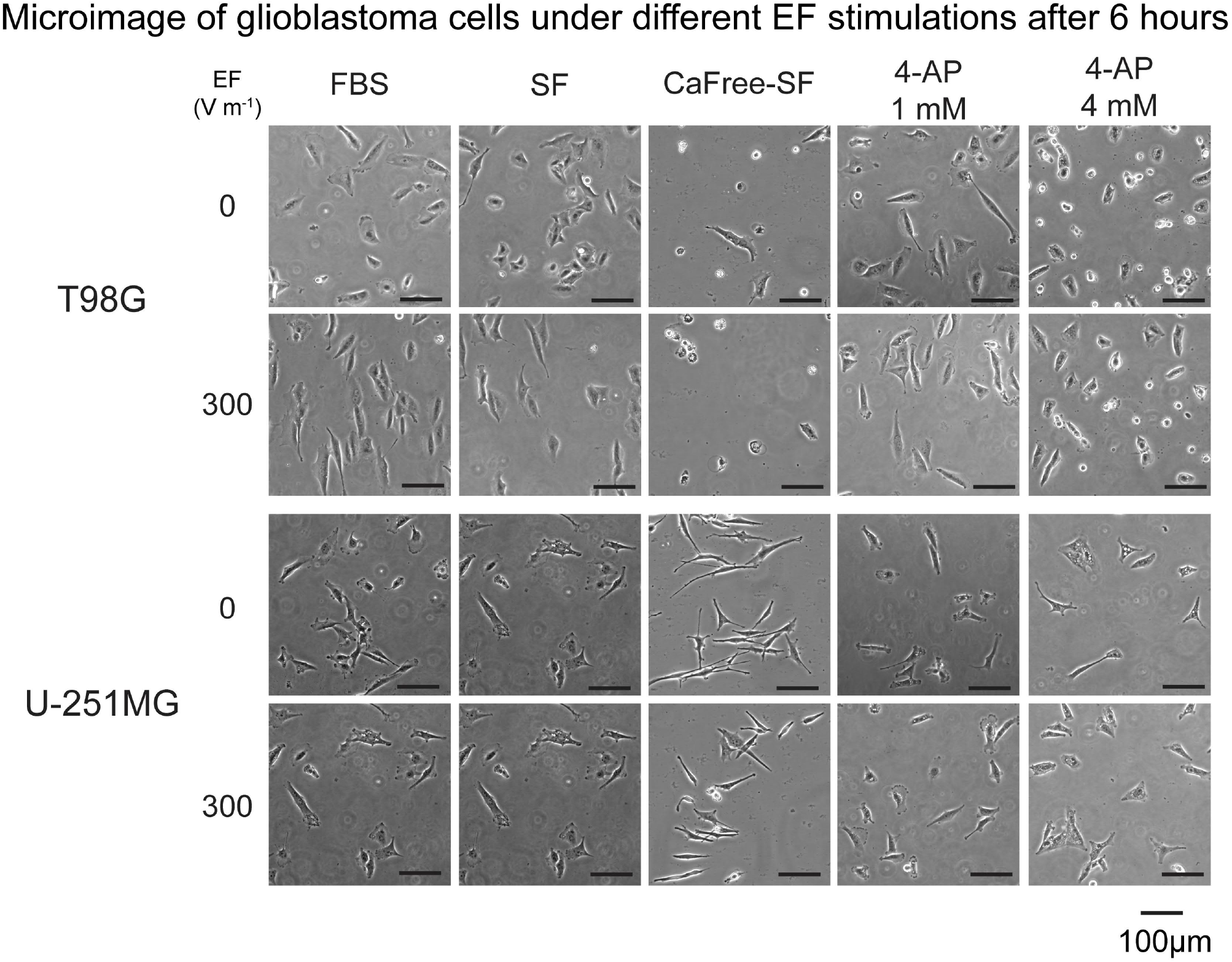
Phase contrast images of T98G & U-251MG cells under EF stimulations in different microenvironments after 6 hours. The scale bars represent 100 *μ*m. 4-AP: 4-aminopyridine. T98G cells demonstrate perpendicular alignment after electric field stimulation in cell culture media with or without FBS. However, without calcium and FBS in the cell culture medium, T98G cells’ viability decreases. Furthermore, in cell culture medium supplemented with 4 mM 4-AP, T98G also lose viability and start to detach from the substrate. These trends were not observed in the U-251MG cells, suggesting heterogeneity between cell lines.

**FIG. S.10:**
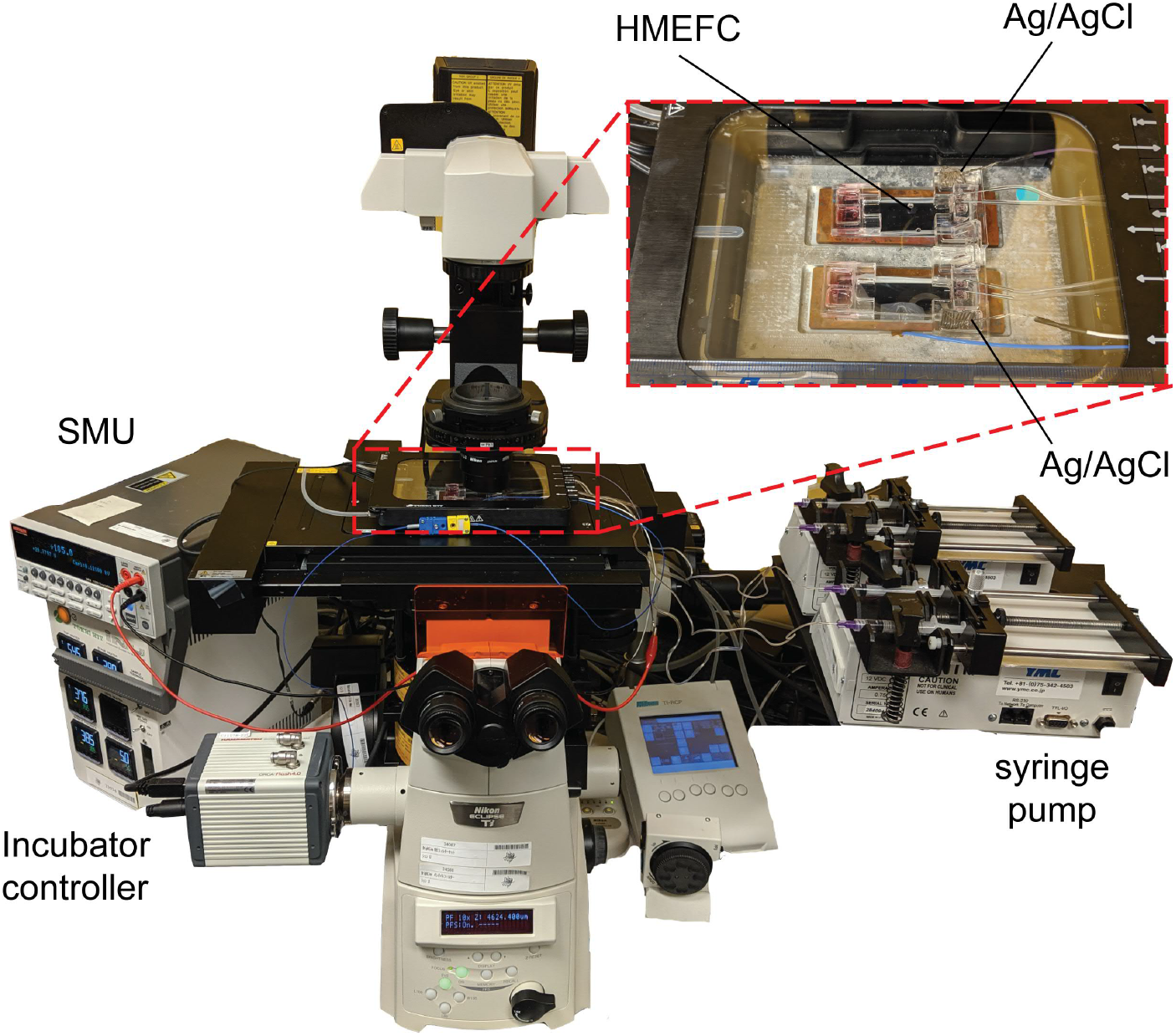
The photoimage of the experimental setup for high throughput experiments with the hybrid multiple electric field chip (HMEFC). The red dashed box indicates the magnified region with the hybrid PMMA/PDMS HMEFC in an on-stage incubator. SMU: source measure unit

**FIG. S.11:**
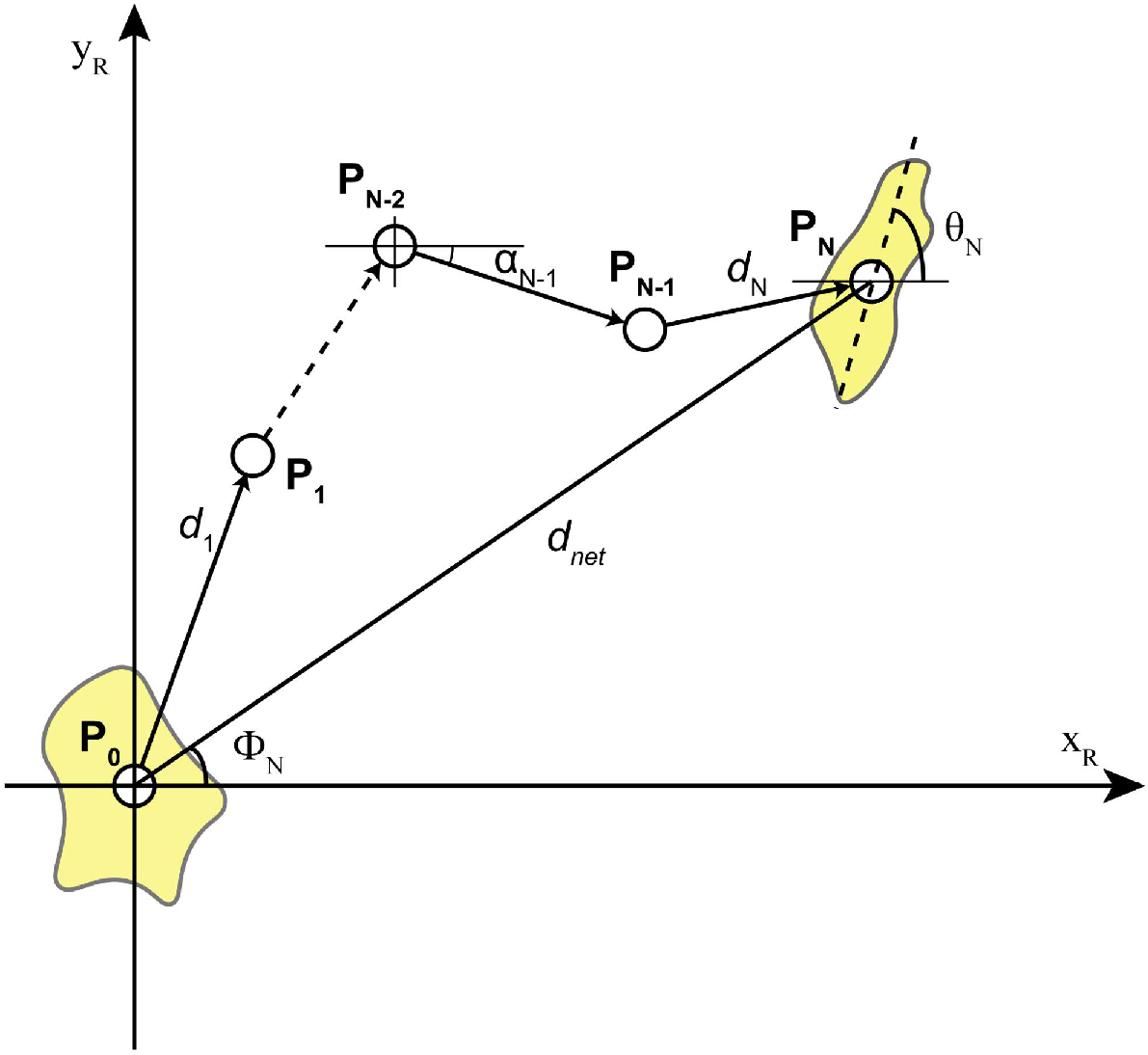
The single-cell migration parameters extracted from the electrotaxis experiments.

## Supplementary Tables

**TABLE.S.1:**
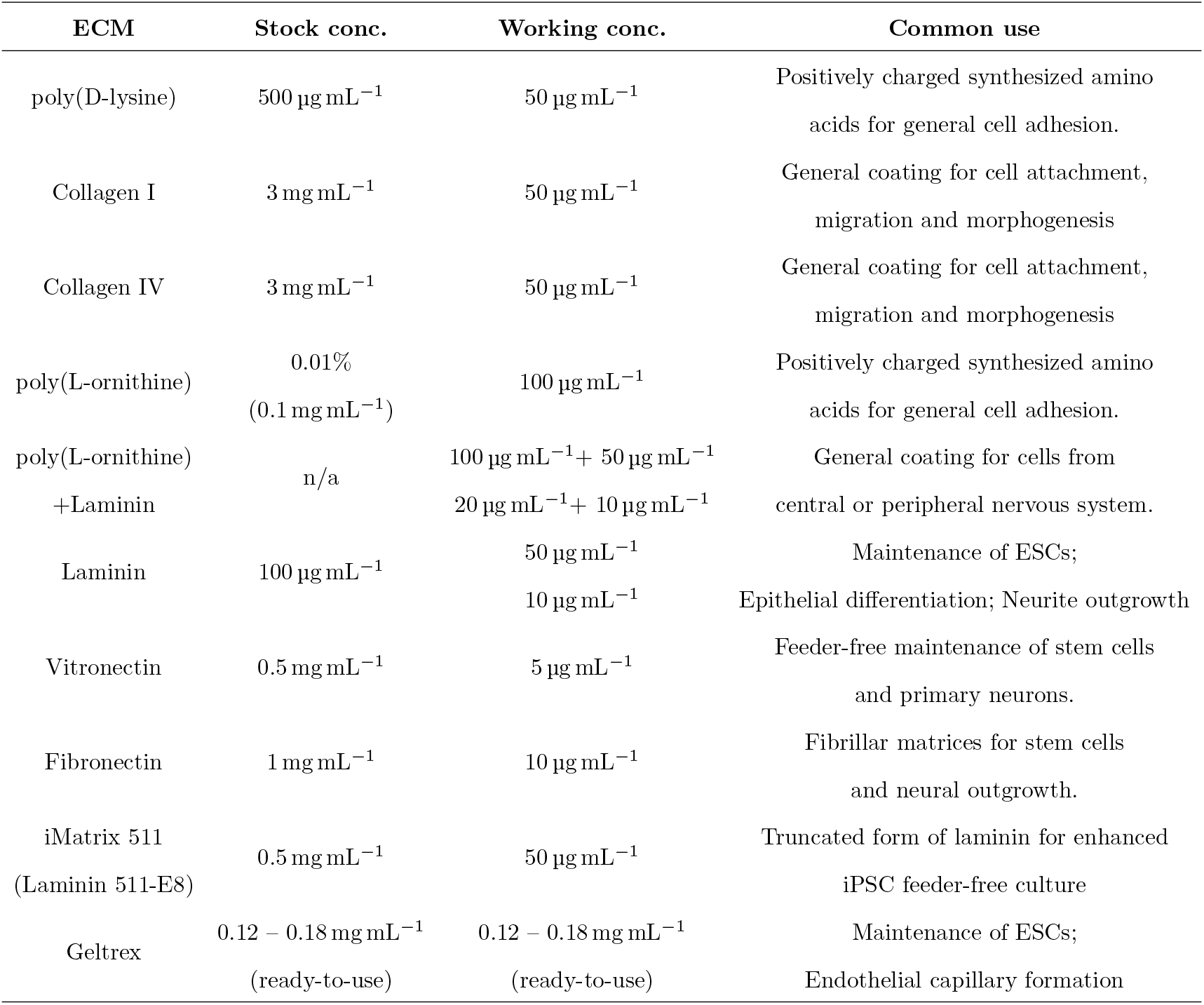
Commonly used extracellular matrix coatings for cell migration.

| ECM | Stock conc. | Working conc. | Common use |
| --- | --- | --- | --- |
| poly(D-lysine) | 500 $\mu\text{g mL}^{-1}$ | 50 $\mu\text{g mL}^{-1}$ | Positively charged synthesized amino acids for general cell adhesion. |
| Collagen I | 3 $\text{mg mL}^{-1}$ | 50 $\mu\text{g mL}^{-1}$ | General coating for cell attachment, migration and morphogenesis |
| Collagen IV | 3 $\text{mg mL}^{-1}$ | 50 $\mu\text{g mL}^{-1}$ | General coating for cell attachment, migration and morphogenesis |
| poly(L-ornithine) | 0.01%<br>(0.1 $\text{mg mL}^{-1}$) | 100 $\mu\text{g mL}^{-1}$ | Positively charged synthesized amino acids for general cell adhesion. |
| poly(L-ornithine)<br>+Laminin | n/a | 100 $\mu\text{g mL}^{-1}$ + 50 $\mu\text{g mL}^{-1}$<br>20 $\mu\text{g mL}^{-1}$ + 10 $\mu\text{g mL}^{-1}$ | General coating for cells from central or peripheral nervous system. |
| Laminin | 100 $\mu\text{g mL}^{-1}$ | 50 $\mu\text{g mL}^{-1}$<br>10 $\mu\text{g mL}^{-1}$ | Maintenance of ESCs;<br>Epithelial differentiation; Neurite outgrowth |
| Vitronectin | 0.5 $\text{mg mL}^{-1}$ | 5 $\mu\text{g mL}^{-1}$ | Feeder-free maintenance of stem cells and primary neurons. |
| Fibronectin | 1 $\text{mg mL}^{-1}$ | 10 $\mu\text{g mL}^{-1}$ | Fibrillar matrices for stem cells and neural outgrowth. |
| iMatrix 511<br>(Laminin 511-E8) | 0.5 $\text{mg mL}^{-1}$ | 50 $\mu\text{g mL}^{-1}$ | Truncated form of laminin for enhanced iPSC feeder-free culture |
| Geltrex | 0.12 – 0.18 $\text{mg mL}^{-1}$<br>(ready-to-use) | 0.12 – 0.18 $\text{mg mL}^{-1}$<br>(ready-to-use) | Maintenance of ESCs;<br>Endothelial capillary formation |

**TABLE.S.2:**
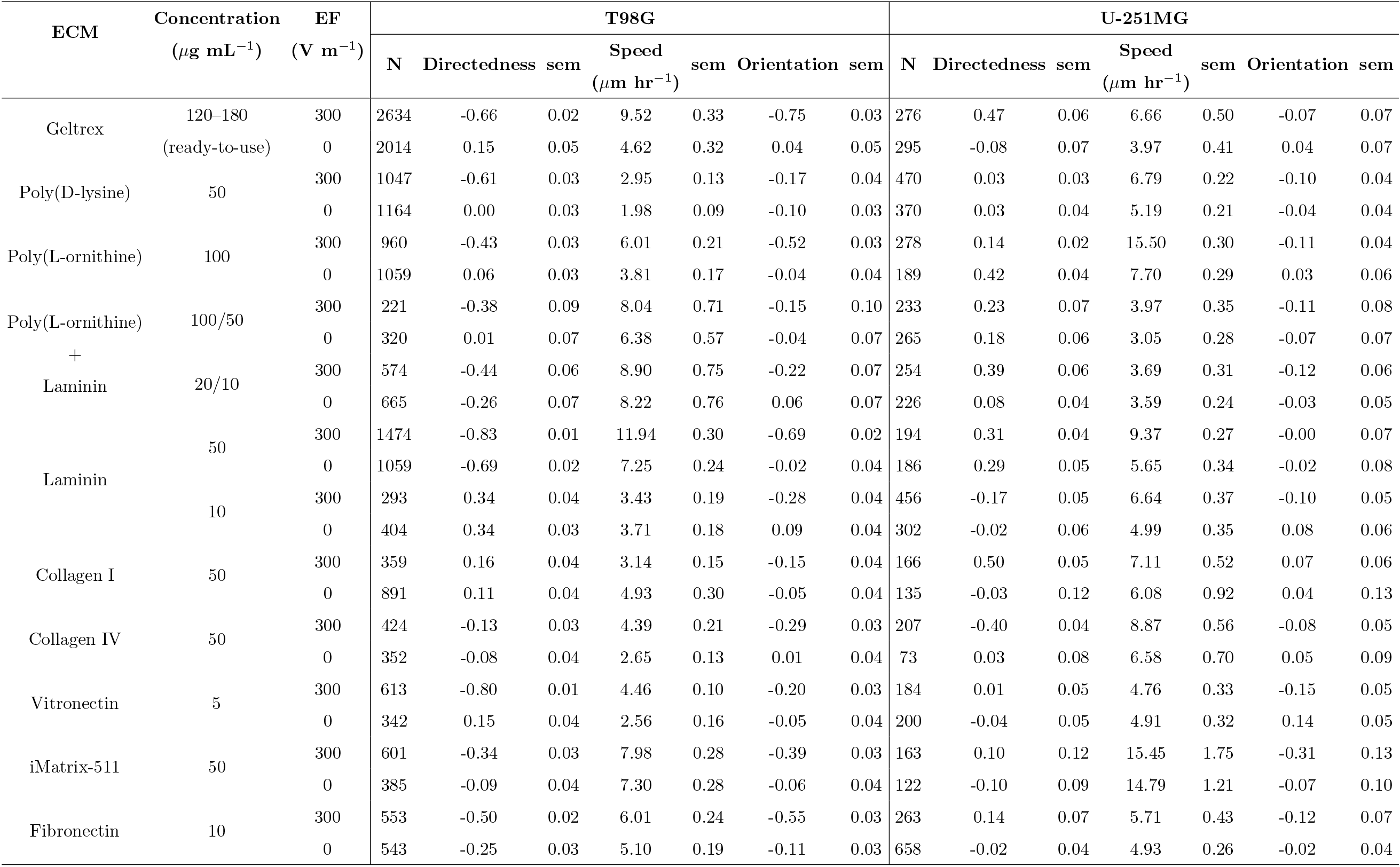
The results of glioblastoma electrotaxis on various ECM coatings in HMEFC after 6 hours. sem: standard error of mean.

**TABLE.S.3:**
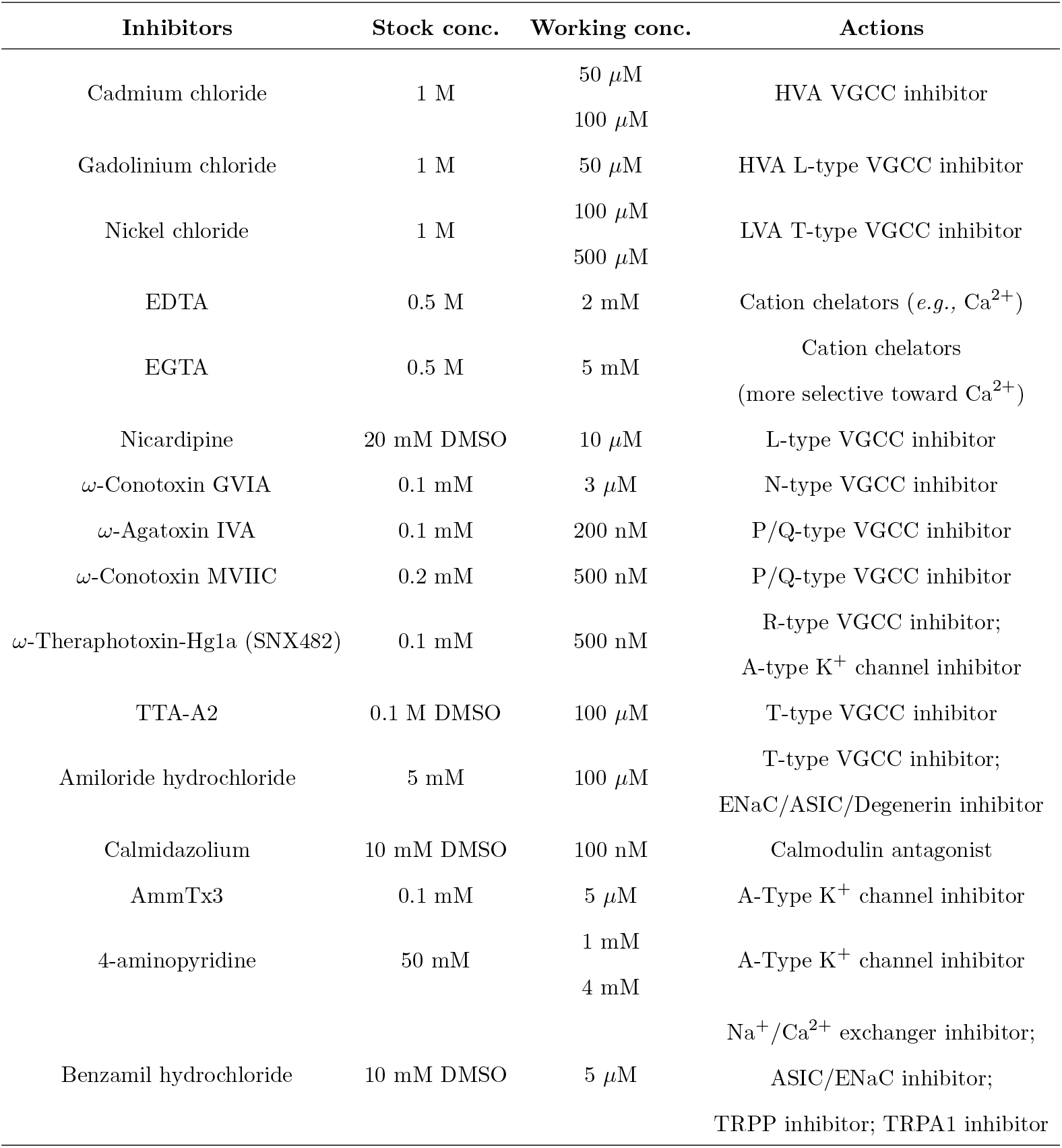
Pharmacological inhibitors to study various ion channels in glioma electrotaxis.

**TABLE.S.4:**
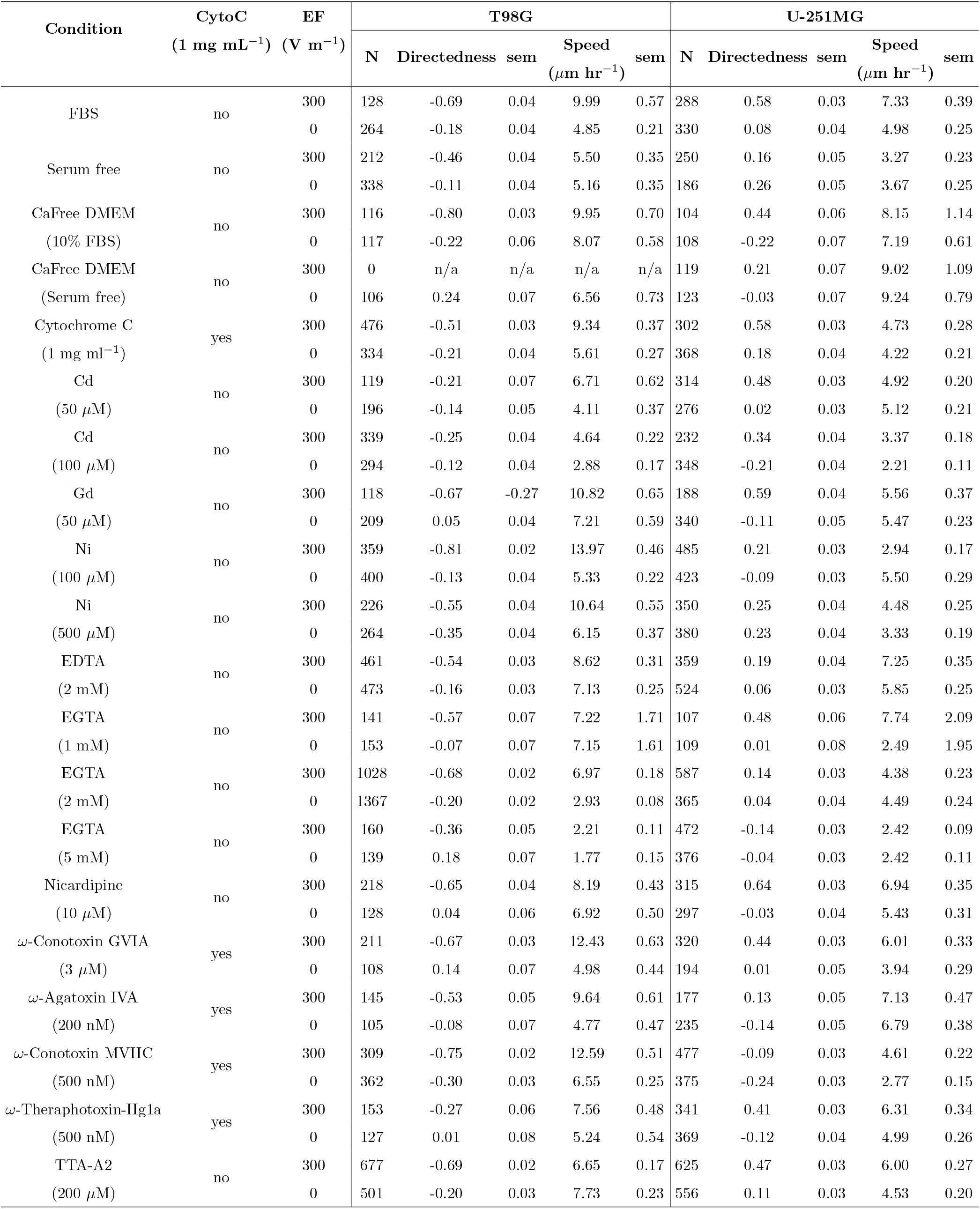
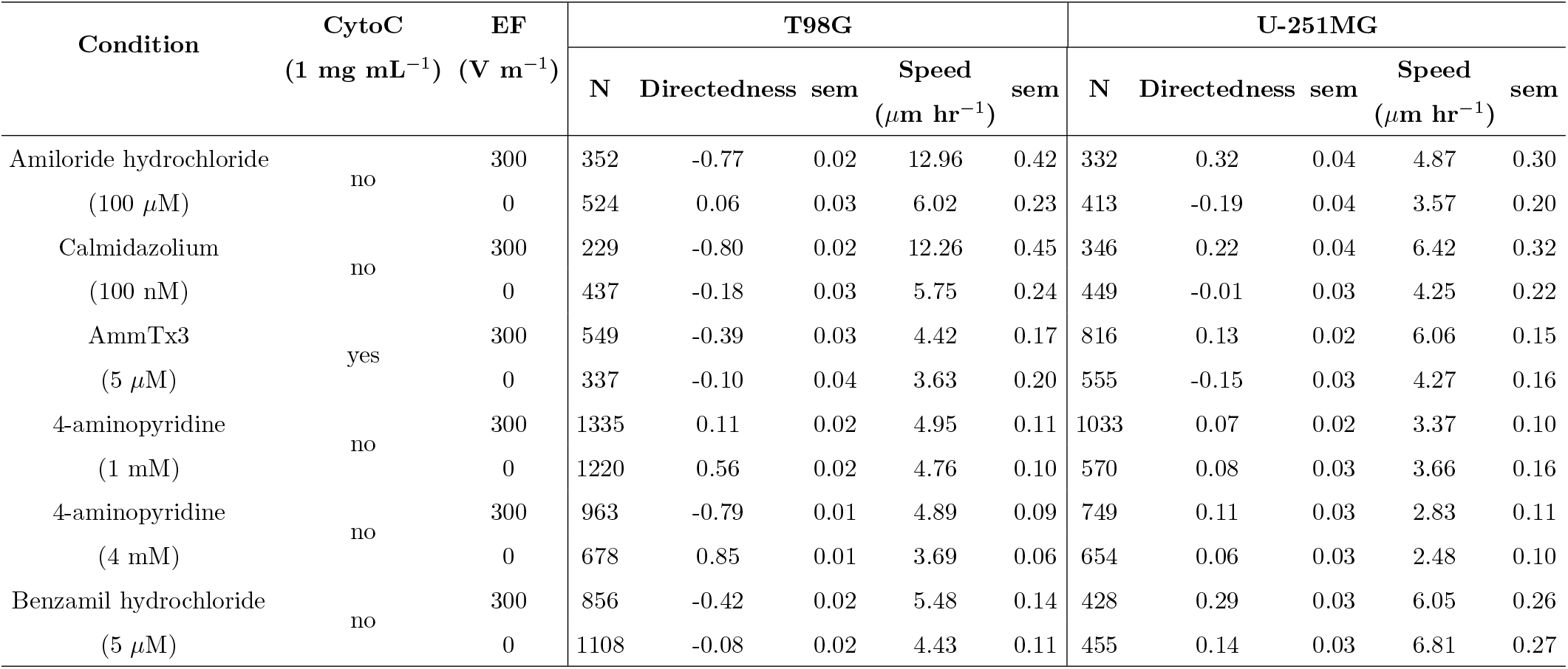
The detailed results of glioblastoma electrotaxis in the presence of antagonists against ion channels or calcium signaling after 6 hours. CytoC: Cytochrome C; sem: standard error of mean.

| Condition | CytoC<br>(1 mg mL<sup>-1</sup>) | EF<br>(V m<sup>-1</sup>) | T98G |  |  |  |  | U-251MG |  |  |  |  |
| --- | --- | --- | --- | --- | --- | --- | --- | --- | --- | --- | --- | --- |
| | | | N | Directedness | sem | Speed<br>($\mu\text{m hr}^{-1}$) | sem | N | Directedness | sem | Speed<br>($\mu\text{m hr}^{-1}$) | sem |
| FBS | no | 300 | 128 | -0.69 | 0.04 | 9.99 | 0.57 | 288 | 0.58 | 0.03 | 7.33 | 0.39 |
|  |  | 0 | 264 | -0.18 | 0.04 | 4.85 | 0.21 | 330 | 0.08 | 0.04 | 4.98 | 0.25 |
| Serum free | no | 300 | 212 | -0.46 | 0.04 | 5.50 | 0.35 | 250 | 0.16 | 0.05 | 3.27 | 0.23 |
|  |  | 0 | 338 | -0.11 | 0.04 | 5.16 | 0.35 | 186 | 0.26 | 0.05 | 3.67 | 0.25 |
| CaFree DMEM<br>(10% FBS) | no | 300 | 116 | -0.80 | 0.03 | 9.95 | 0.70 | 104 | 0.44 | 0.06 | 8.15 | 1.14 |
|  |  | 0 | 117 | -0.22 | 0.06 | 8.07 | 0.58 | 108 | -0.22 | 0.07 | 7.19 | 0.61 |
| CaFree DMEM<br>(Serum free) | no | 300 | 0 | n/a | n/a | n/a | n/a | 119 | 0.21 | 0.07 | 9.02 | 1.09 |
|  |  | 0 | 106 | 0.24 | 0.07 | 6.56 | 0.73 | 123 | -0.03 | 0.07 | 9.24 | 0.79 |
| Cytochrome C<br>(1 mg mL<sup>-1</sup>) | yes | 300 | 476 | -0.51 | 0.03 | 9.34 | 0.37 | 302 | 0.58 | 0.03 | 4.73 | 0.28 |
|  |  | 0 | 334 | -0.21 | 0.04 | 5.61 | 0.27 | 368 | 0.18 | 0.04 | 4.22 | 0.21 |
| Cd<br>(50 $\mu\text{M}$) | no | 300 | 119 | -0.21 | 0.07 | 6.71 | 0.62 | 314 | 0.48 | 0.03 | 4.92 | 0.20 |
|  |  | 0 | 196 | -0.14 | 0.05 | 4.11 | 0.37 | 276 | 0.02 | 0.03 | 5.12 | 0.21 |
| Cd<br>(100 $\mu\text{M}$) | no | 300 | 339 | -0.25 | 0.04 | 4.64 | 0.22 | 232 | 0.34 | 0.04 | 3.37 | 0.18 |
|  |  | 0 | 294 | -0.12 | 0.04 | 2.88 | 0.17 | 348 | -0.21 | 0.04 | 2.21 | 0.11 |
| Gd<br>(50 $\mu\text{M}$) | no | 300 | 118 | -0.67 | -0.27 | 10.82 | 0.65 | 188 | 0.59 | 0.04 | 5.56 | 0.37 |
|  |  | 0 | 209 | 0.05 | 0.04 | 7.21 | 0.59 | 340 | -0.11 | 0.05 | 5.47 | 0.23 |
| Ni<br>(100 $\mu\text{M}$) | no | 300 | 359 | -0.81 | 0.02 | 13.97 | 0.46 | 485 | 0.21 | 0.03 | 2.94 | 0.17 |
|  |  | 0 | 400 | -0.13 | 0.04 | 5.33 | 0.22 | 423 | -0.09 | 0.03 | 5.50 | 0.29 |
| Ni<br>(500 $\mu\text{M}$) | no | 300 | 226 | -0.55 | 0.04 | 10.64 | 0.55 | 350 | 0.25 | 0.04 | 4.48 | 0.25 |
|  |  | 0 | 264 | -0.35 | 0.04 | 6.15 | 0.37 | 380 | 0.23 | 0.04 | 3.33 | 0.19 |
| EDTA<br>(2 mM) | no | 300 | 461 | -0.54 | 0.03 | 8.62 | 0.31 | 359 | 0.19 | 0.04 | 7.25 | 0.35 |
|  |  | 0 | 473 | -0.16 | 0.03 | 7.13 | 0.25 | 524 | 0.06 | 0.03 | 5.85 | 0.25 |
| EGTA<br>(1 mM) | no | 300 | 141 | -0.57 | 0.07 | 7.22 | 1.71 | 107 | 0.48 | 0.06 | 7.74 | 2.09 |
|  |  | 0 | 153 | -0.07 | 0.07 | 7.15 | 1.61 | 109 | 0.01 | 0.08 | 2.49 | 1.95 |
| EGTA<br>(2 mM) | no | 300 | 1028 | -0.68 | 0.02 | 6.97 | 0.18 | 587 | 0.14 | 0.03 | 4.38 | 0.23 |
|  |  | 0 | 1367 | -0.20 | 0.02 | 2.93 | 0.08 | 365 | 0.04 | 0.04 | 4.49 | 0.24 |
| EGTA<br>(5 mM) | no | 300 | 160 | -0.36 | 0.05 | 2.21 | 0.11 | 472 | -0.14 | 0.03 | 2.42 | 0.09 |
|  |  | 0 | 139 | 0.18 | 0.07 | 1.77 | 0.15 | 376 | -0.04 | 0.03 | 2.42 | 0.11 |
| Nicardipine<br>(10 $\mu\text{M}$) | no | 300 | 218 | -0.65 | 0.04 | 8.19 | 0.43 | 315 | 0.64 | 0.03 | 6.94 | 0.35 |
|  |  | 0 | 128 | 0.04 | 0.06 | 6.92 | 0.50 | 297 | -0.03 | 0.04 | 5.43 | 0.31 |
| ω-Conotoxin GVIA<br>(3 $\mu\text{M}$) | yes | 300 | 211 | -0.67 | 0.03 | 12.43 | 0.63 | 320 | 0.44 | 0.03 | 6.01 | 0.33 |
|  |  | 0 | 108 | 0.14 | 0.07 | 4.98 | 0.44 | 194 | 0.01 | 0.05 | 3.94 | 0.29 |
| ω-Agatoxin IVA<br>(200 nM) | yes | 300 | 145 | -0.53 | 0.05 | 9.64 | 0.61 | 177 | 0.13 | 0.05 | 7.13 | 0.47 |
|  |  | 0 | 105 | -0.08 | 0.07 | 4.77 | 0.47 | 235 | -0.14 | 0.05 | 6.79 | 0.38 |
| ω-Conotoxin MVIIC<br>(500 nM) | yes | 300 | 309 | -0.75 | 0.02 | 12.59 | 0.51 | 477 | -0.09 | 0.03 | 4.61 | 0.22 |
|  |  | 0 | 362 | -0.30 | 0.03 | 6.55 | 0.25 | 375 | -0.24 | 0.03 | 2.77 | 0.15 |
| ω-Theraphotoxin-Hg1a<br>(500 nM) | yes | 300 | 153 | -0.27 | 0.06 | 7.56 | 0.48 | 341 | 0.41 | 0.03 | 6.31 | 0.34 |
|  |  | 0 | 127 | 0.01 | 0.08 | 5.24 | 0.54 | 369 | -0.12 | 0.04 | 4.99 | 0.26 |
| TTA-A2<br>(200 $\mu\text{M}$) | no | 300 | 677 | -0.69 | 0.02 | 6.65 | 0.17 | 625 | 0.47 | 0.03 | 6.00 | 0.27 |
|  |  | 0 | 501 | -0.20 | 0.03 | 7.73 | 0.23 | 556 | 0.11 | 0.03 | 4.53 | 0.20 |
|  |  |  | N | Directedness | sem | Speed<br>(μm hr<sup>-1</sup>) | sem | N | Directedness | sem | Speed<br>(μm hr<sup>-1</sup>) | sem |
| Amiloride hydrochloride | no | 300 | 352 | -0.77 | 0.02 | 12.96 | 0.42 | 332 | 0.32 | 0.04 | 4.87 | 0.30 |
| (100 μM) |  | 0 | 524 | 0.06 | 0.03 | 6.02 | 0.23 | 413 | -0.19 | 0.04 | 3.57 | 0.20 |
| Calmidazolium | no | 300 | 229 | -0.80 | 0.02 | 12.26 | 0.45 | 346 | 0.22 | 0.04 | 6.42 | 0.32 |
| (100 nM) |  | 0 | 437 | -0.18 | 0.03 | 5.75 | 0.24 | 449 | -0.01 | 0.03 | 4.25 | 0.22 |
| AmmTx3 | yes | 300 | 549 | -0.39 | 0.03 | 4.42 | 0.17 | 816 | 0.13 | 0.02 | 6.06 | 0.15 |
| (5 μM) |  | 0 | 337 | -0.10 | 0.04 | 3.63 | 0.20 | 555 | -0.15 | 0.03 | 4.27 | 0.16 |
| 4-aminopyridine | no | 300 | 1335 | 0.11 | 0.02 | 4.95 | 0.11 | 1033 | 0.07 | 0.02 | 3.37 | 0.10 |
| (1 mM) |  | 0 | 1220 | 0.56 | 0.02 | 4.76 | 0.10 | 570 | 0.08 | 0.03 | 3.66 | 0.16 |
| 4-aminopyridine | no | 300 | 963 | -0.79 | 0.01 | 4.89 | 0.09 | 749 | 0.11 | 0.03 | 2.83 | 0.11 |
| (4 mM) |  | 0 | 678 | 0.85 | 0.01 | 3.69 | 0.06 | 654 | 0.06 | 0.03 | 2.48 | 0.10 |
| Benzamil hydrochloride | no | 300 | 856 | -0.42 | 0.02 | 5.48 | 0.14 | 428 | 0.29 | 0.03 | 6.05 | 0.26 |
| (5 μM) |  | 0 | 1108 | -0.08 | 0.02 | 4.43 | 0.11 | 455 | 0.14 | 0.03 | 6.81 | 0.27 |

**TABLE.S.5:** Step-centric and cell-centric variables for describing single-cell migration.

| Step-centric features |  |
| --- | --- |
| Instantaneous displacement | $d_i = \sqrt{(x_i - x_{i-1})^2 + (y_i - y_{i-1})^2}$ |
| Instantaneous speed | $s_i = d(p_{i-1}, p_i) / \Delta t$ |
| Turning angle | $\alpha_i = \tan^{-1} \frac{(y_i - y_{i-1})}{(x_i - x_{i-1})}$ |
| Directional autocorrelation | $dir - aut_i = \cos(\alpha_i - \alpha_{i-1})$ |
| Instantaneous directedness | $Index_{directedness_{instant}} = \sum_{i=1}^N \frac{\cos \Phi_{instant_i}}{N} = \sum_{i=1}^N \frac{(x_i - x_{i-1})}{d_i \times N} = \frac{1}{N} \sum_{i=1}^N \frac{(x_i - x_{i-1})}{\sqrt{(x_i - x_{i-1})^2 + (y_i - y_{i-1})^2}}$ |
| Cell-centric features |  |
| Cumulative distance | $d_{total} = \sum_{i=1}^N d(p_{i-1}, p_i)$ |
| Net trigonometric distance<br>(Euclidean distance) | $d_{net} = d(p_0, p_N) = \sqrt{(x_N - x_0)^2 + (y_N - y_0)^2}$ |
| Euclidean velocity | $\bar{v}_{net} = \frac{d_{net}}{t_{elapsed}}$ |
| End-point directionality ratio | $ep\_dr = \frac{d_{net}}{d_{tot}}$ |
| Orientation | $Index_{orientation} = \sum_{i=1}^N \frac{\cos 2\theta_i}{N}$ |
| Directedness | $Index_{directedness} = \sum_{i=1}^N \frac{\cos \Phi_i}{N} = \sum_{i=1}^N \frac{(x_i - x_0)}{d_{net} \times N} = \frac{1}{N} \sum_{i=1}^N \frac{(x_i - x_0)}{\sqrt{(x_i - x_0)^2 + (y_i - y_0)^2}}$ |

## Supplementary Videos

**VIDEO. S.1:**
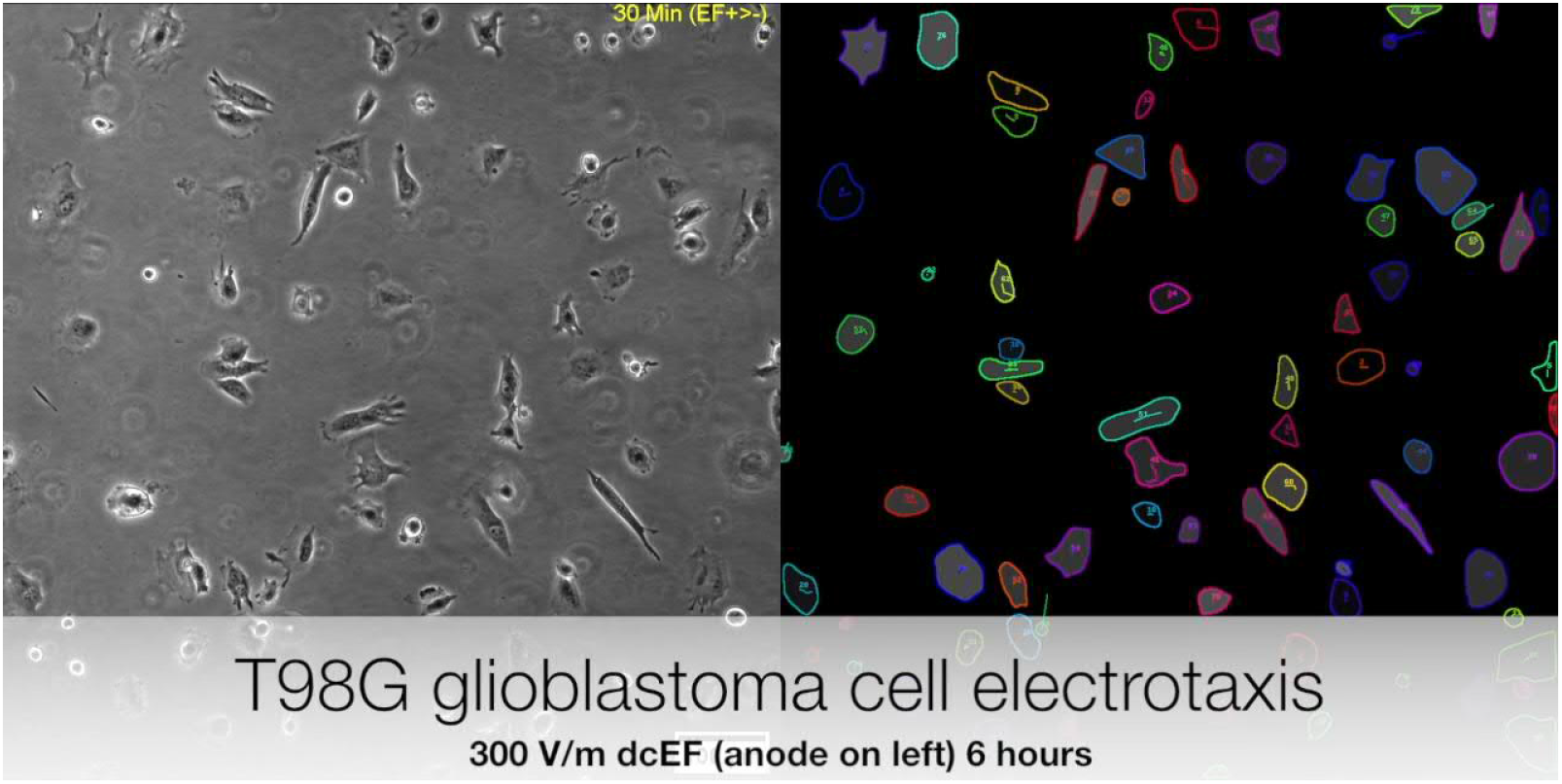
Video clip showing the electrotaxis of T98G glioblastoma cells under 300 V m ^−1^ dcEF for six hours and the respective tracking results using *Usiigaci* software.

**VIDEO. S.2:**
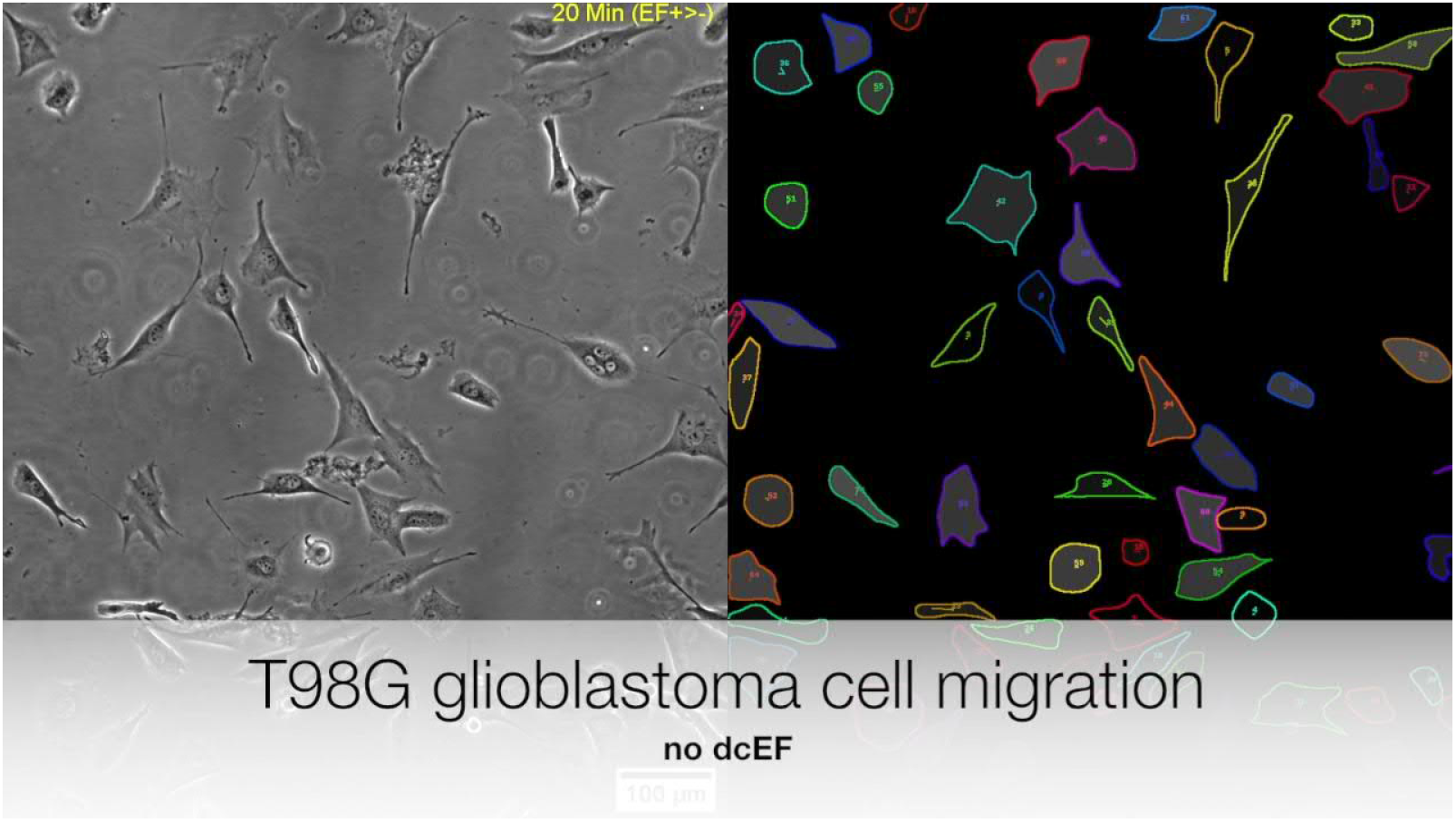
Video clip showing the random migration of T98G glioblastoma cells for six hours and the respective tracking results using *Usiigaci* software.

**VIDEO. S.3:**
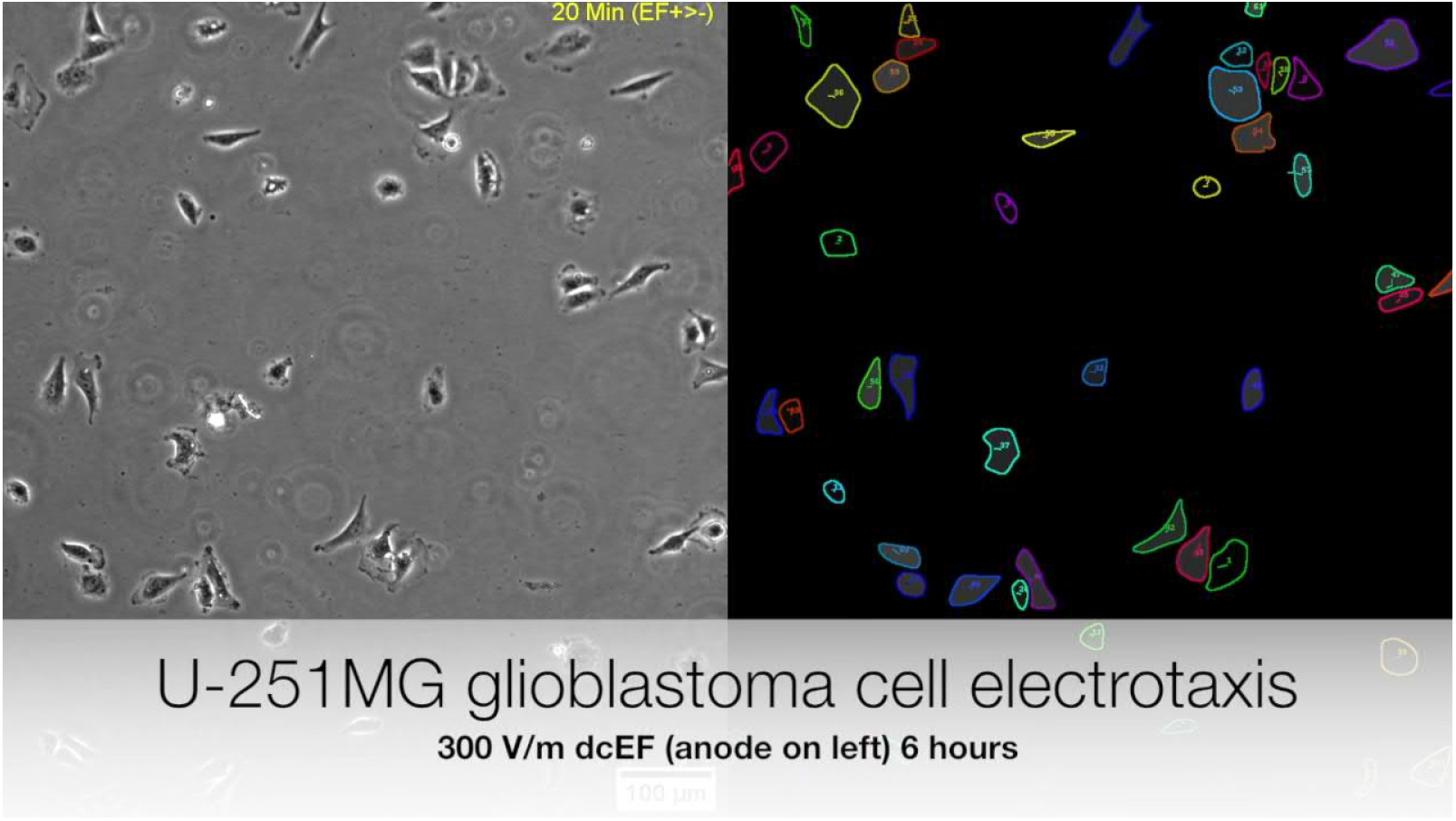
Video clip showing the electrotaxis of U-251MG glioblastoma cells under 300 V m ^−1^ dcEF for six hours and the respective tracking results using *Usiigaci* software.

**VIDEO. S.4:**
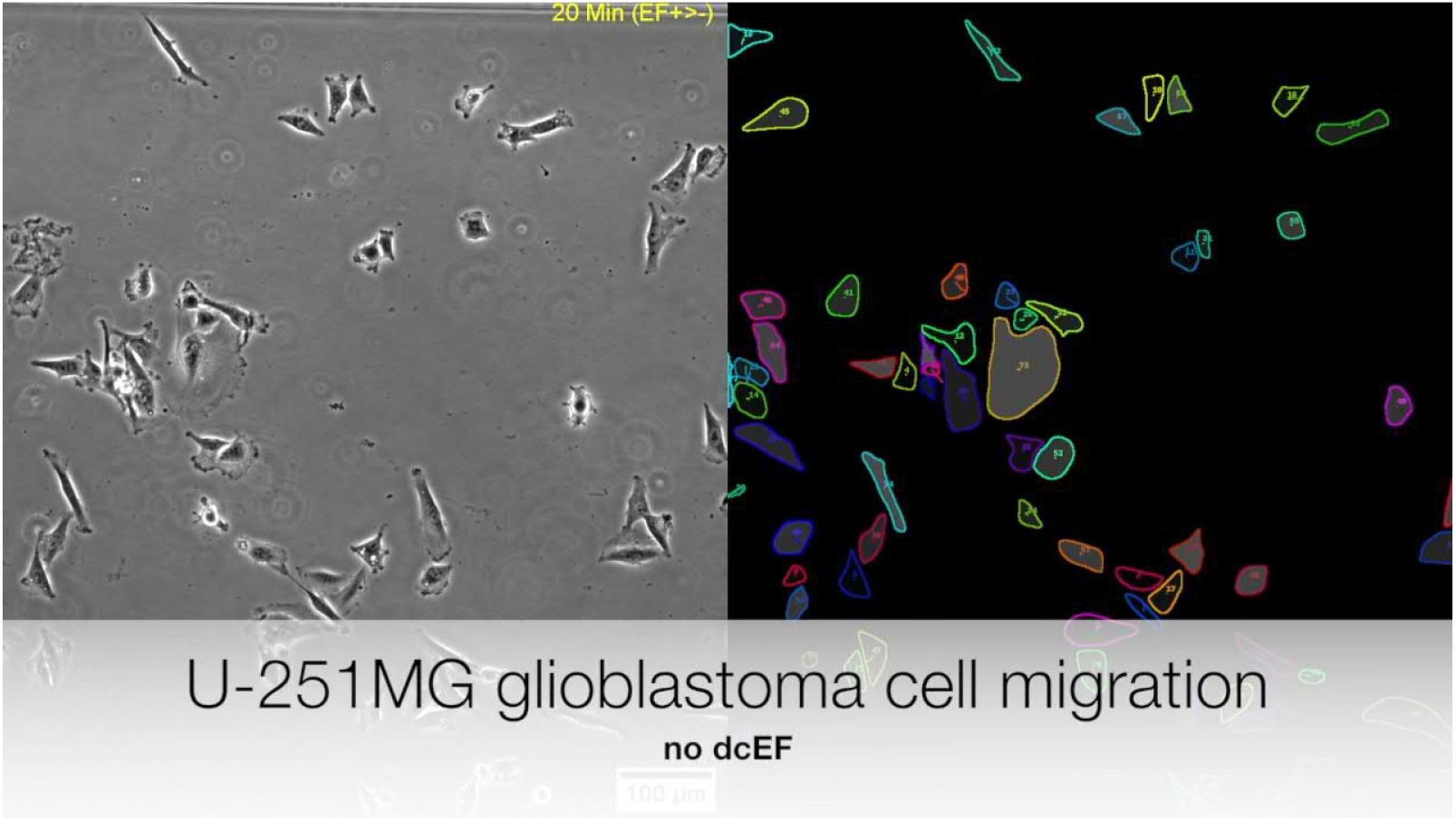
Video clip showing the random migration of U-251MG glioblastoma cells for six hours and the respective tracking results using *Usiigaci* software.

**VIDEO. S.5:**
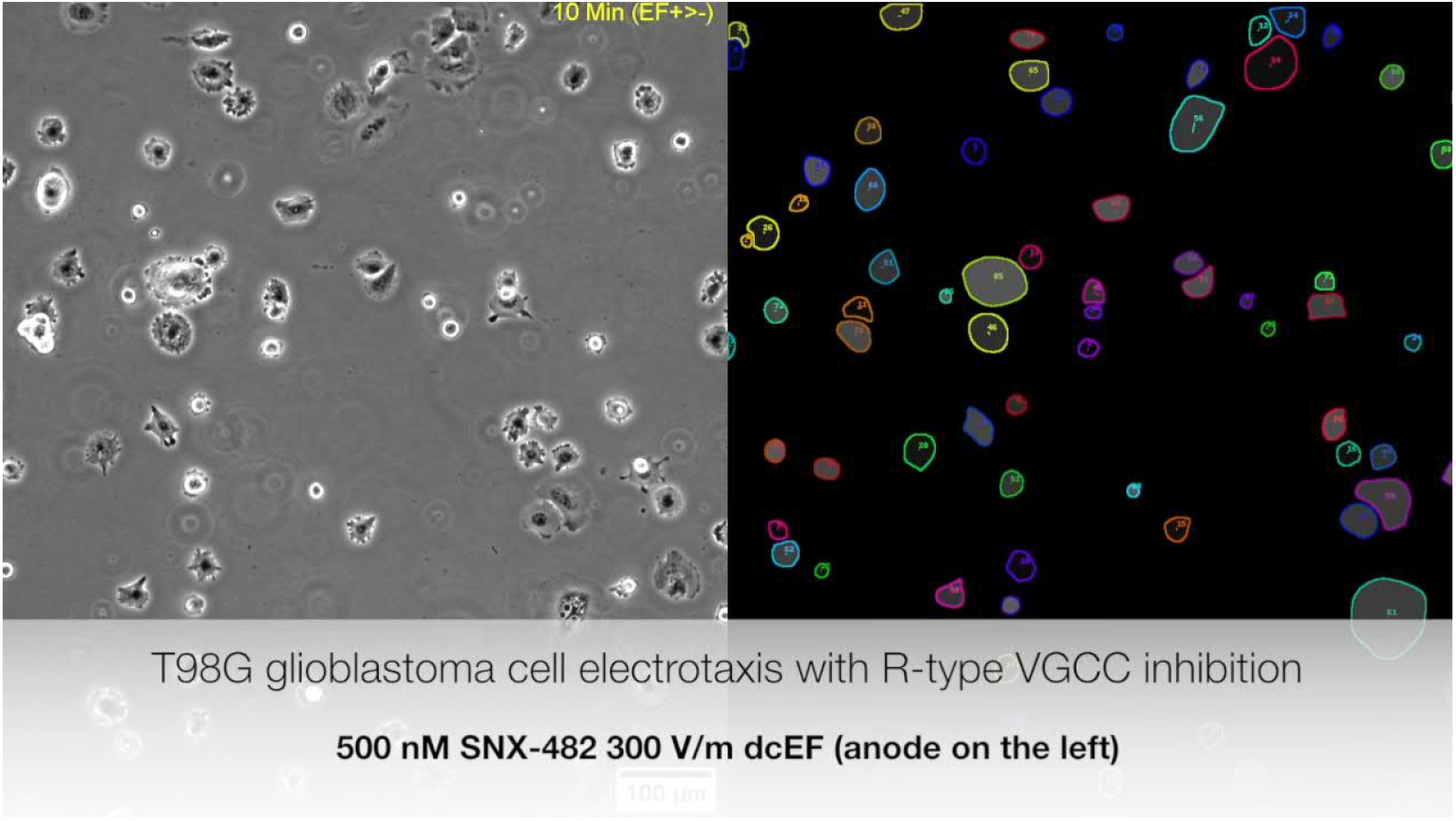
Video clip showing the electrotaxis of T98G glioblastoma cells suppressed with 500 nM SNX482 on R-type VGCC under 300 V m ^−1^ dcEF for six hours and the respective tracking results using *Usiigaci* software.

**VIDEO. S.6:**
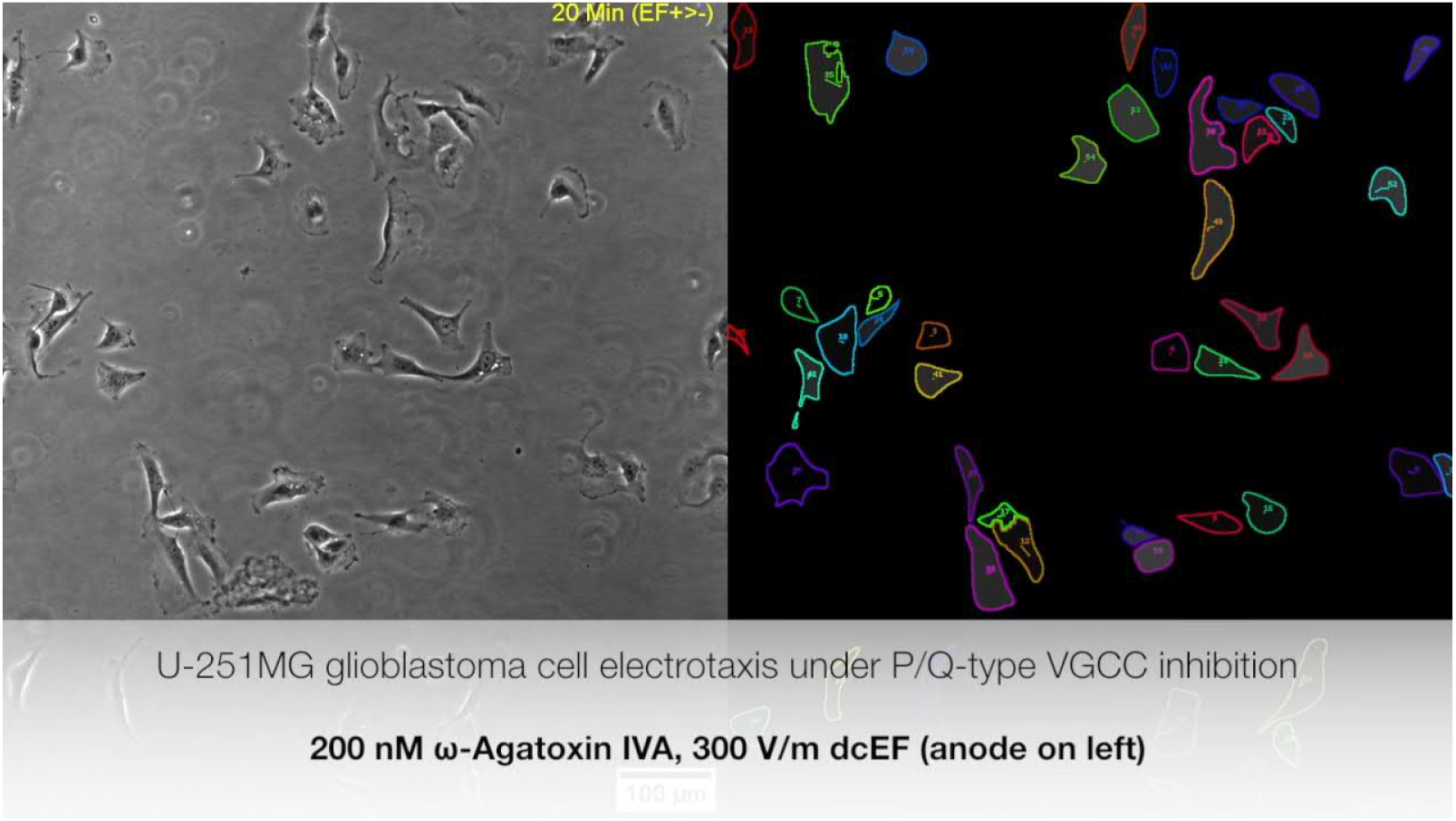
Video clip showing the electrotaxis of U-251MG glioblastoma cells suppressed with 200 nM Agatoxin IVA on P/Q-type VGCC under 300 V m ^−1^ dcEF for six hours and the respective tracking results using *Usiigaci* software.

**VIDEO. S.7:**
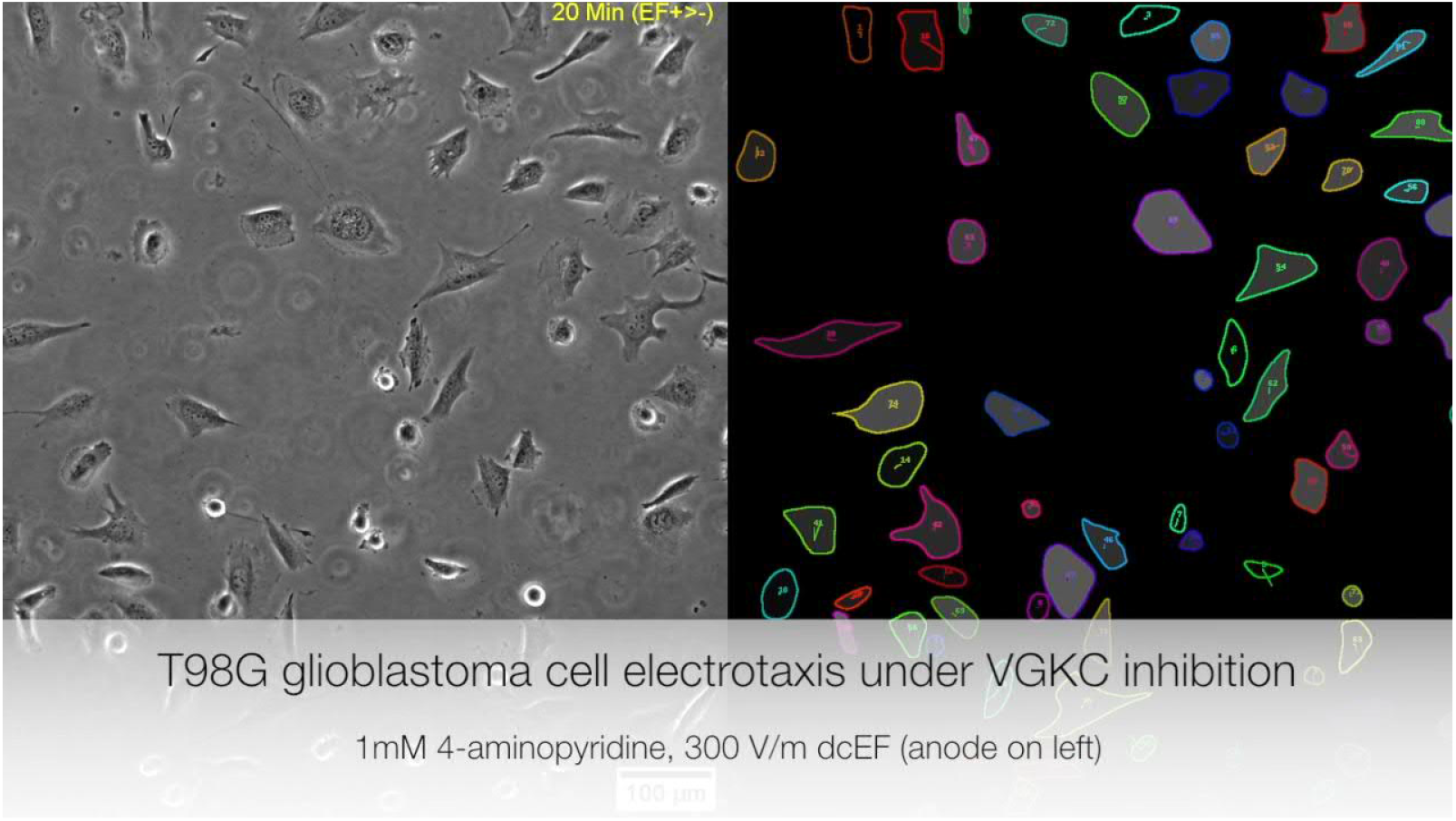
Video clip showing the electrotaxis of T98G glioblastoma cells suppressed with 4-AP on A-type VGKC under 300 V m ^−1^ dcEF for six hours and the respective tracking results using *Usiigaci* software.

**VIDEO. S.8:**
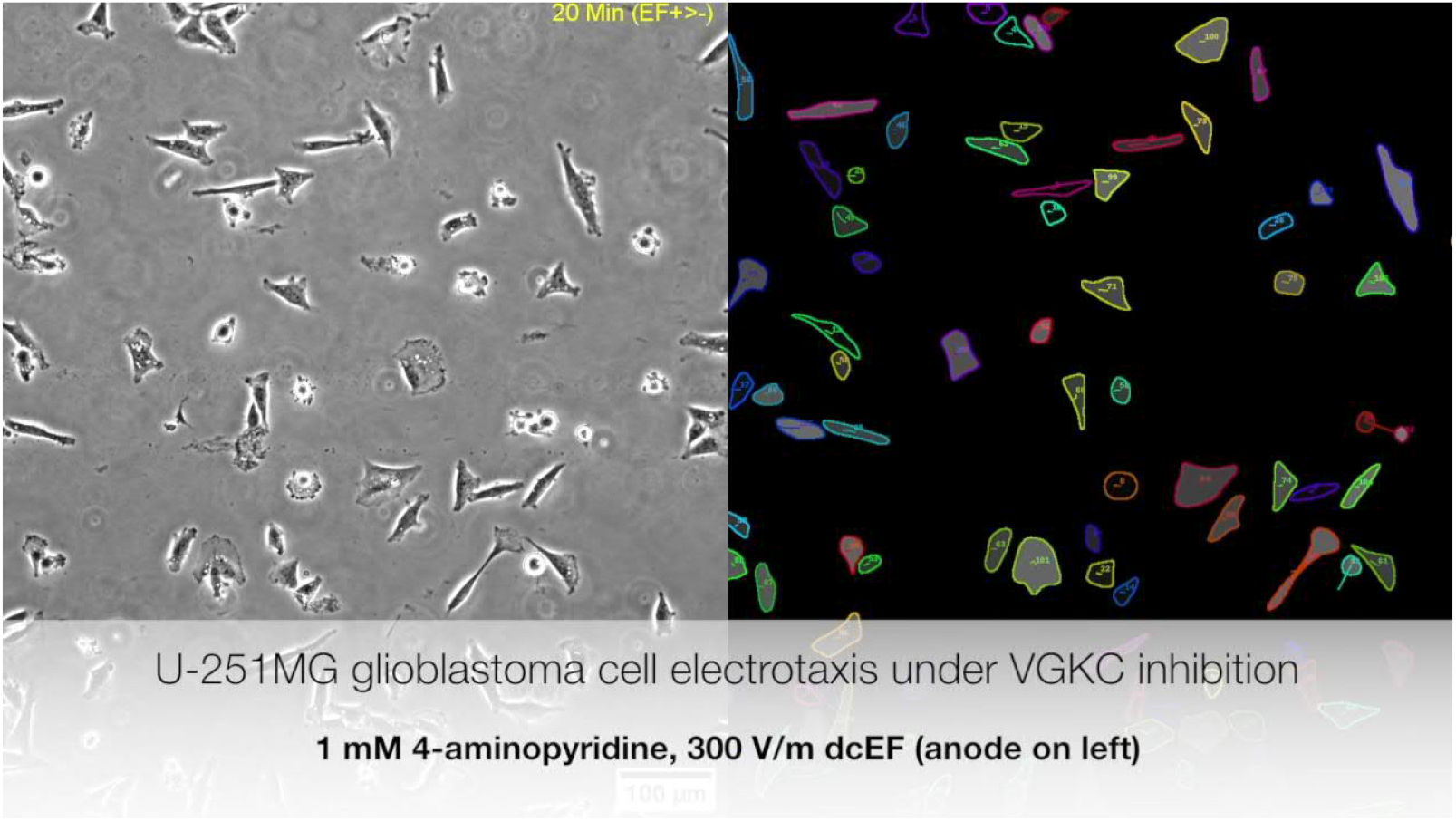
Video clip showing the electrotaxis of U-251MG glioblastoma cells suppressed with 4-AP on A-type VGKC under 300 V m^−1^ dcEF for six hours and the respective tracking results using *Usiigaci* software.

## Supplementary Information

### 1 Hybrid multiple electric field chip (HMEFC) design, simulation and fabrication

#### 1.1 HMEFC design

The HMEFC was designed by the hybrid PMMA/PDMS approach^3^ (FIG. 5), in which the prototyping disadvantages in both materials could be mitigated while the advantages can be combined. In PMMA, complex 3D structures for fluidic routing or reservoir and world-to-chip interface could be quickly prototyped by CO_2_ laser cutting and thermal bonding. However, spatial resolution using this approach was not high to create reliable microfluidic environments. In contrast, highly precise quasi-two dimensional microstructures could be fabricated using the soft lithography technique in PDMS microfluidic chip, however, standard soft lithogarphy was limited by 3D design complexity and world-to-chip interface. Using a dual-energy double-sided tape, PMMA and PDMS could be easily and reversibly bonded^3,4^, enabling broad flexibility in microfluidic design and experiments. In HMEFC, two PMMA components for world-to-chip interface and electrical application were adhered to a PDMS chip which contained a double-layer microchannel where cells were cultured in and observed (FIG. 5b). The electric field was applied from agrose interface next to the outlets that were connected to syringes.

By the double-layer microchannel design, four different strengths of direct current electric field (dcEF) were created in two main channels in a HMEFC and the chemical transport across them were limited by manipulating hydraulic resistances (FIG. 5.c). Two different cell types or chemical treatments could be used in the main channels. The microchannel characteristics were shown in TABLE. S.6. Two 43.8 mm-long, 2 mm-wide, 100 *μ*m-high main channels were connected with three 1.5 mm long, 0.84 mm wide, 10 *μ*m-high interconnecting channels at a spacing of 8.95 mm. The electrical equivalent circuit and hydraulic equivalent circuit were modeled as shown in FIG. S.12.

**TABLE.S.6:** Physical characteristics of the double-layer hybrid multiple electric field chip (HMEFC).

| Channel type | Double layer |  |
| --- | --- | --- |
|  | Interconnection | Main |
| Channel dimension (L×W×H) (mm<sup>3</sup>) | 1.5×0.84×0.01 | 8.95×2×0.1 |
| Hydraulic diameter ($\frac{2wh}{w+h}$) (mm) | 0.01976 | 0.1905 |
| Relative $R_{\Omega}$ (mm<sup>-1</sup>) | 178.99 | 44.75 |
| $R_{\text{hydraulic}}$ (Pa s m<sup>-3</sup>) ($\times 10^9$) | 21616.4 | 55.2 |
| Relative $R_{\text{hydraulic}}$ | 391.88 | 1 |
| Flow velocity simulated by PSPICE at 20 μL min<sup>-1</sup> (m s<sup>-1</sup>) | 2.02E-21 | 1.75E-3 |
| Reynolds number | 4.01E-20 | 0.333 |
| Péclet number | 8.97E-16 | 7460 |

To create multiple dcEFs, the HMEFS were designed by R-2R resistor ladder configuration^3,5,6^ creating different electric fields in section I to IV and section V to VIII (FIG. S.12.a). The cells exposed to the highest dcEF were nearest to the outlets to avoid paracrine signaling from electrically-stimulated cells to un-stimulated cells.

According to Ohm’s law, the electrical resistance of a resistor, *R*, was proportional to the length and inversely proportional to the cross-sectional area:

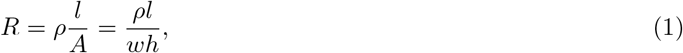

where *ρ*, *l*, *A*, *w*, and *h* were the electrical resistivity of the medium, the length, the cross-sectional area, the width, and the height of the microchannel, respectively. Assuming the electrical resistance of R_1_ being *r*, the relative electrical resistances of other segments (R_2_ : R_11_) could be calculated accordingly. The electric current flowing through each resistor was calculated by Kirchhoff’s circuit law and simulated using PSPICE in the electronic design automation software (OrCAD Lite, Cadence Design Systems, USA) by:

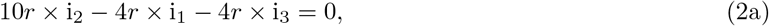

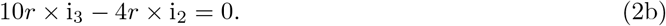

**FIG. S.12:**
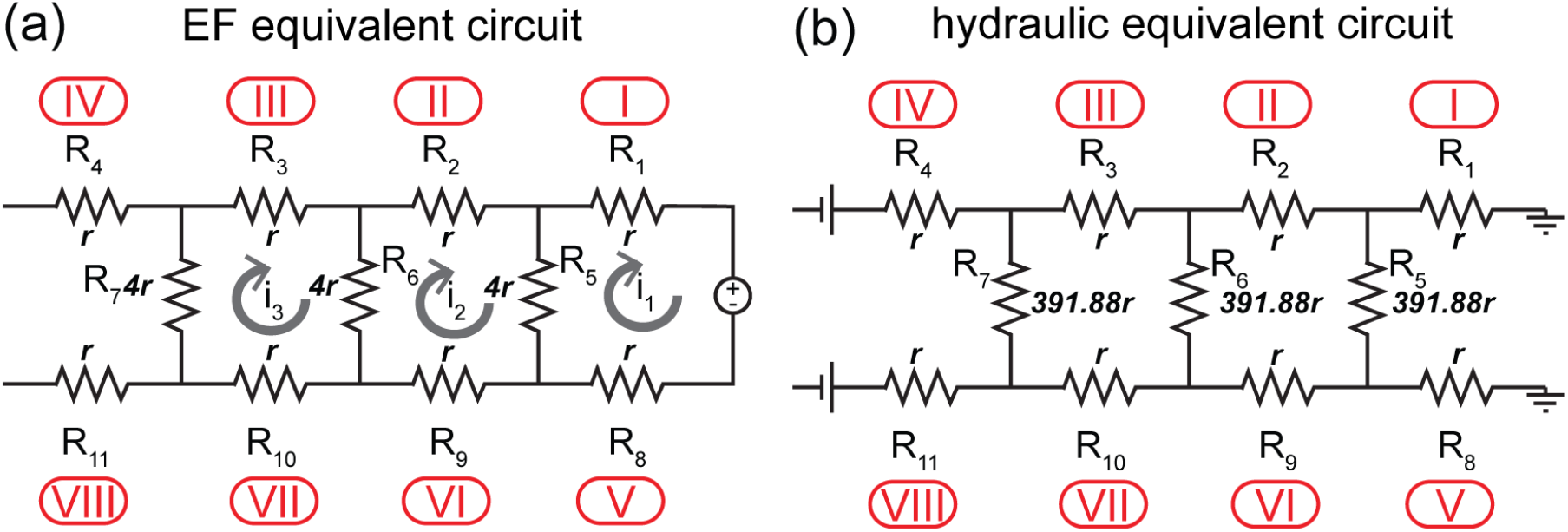
The circuit models of microfluidic design in the hybrid multiple electric field chip (HMEFC). (a) The electric field equivalent circuit; (b) The hydraulic equivalent circuit. The italic *r* depicts relative electrical resistance or relative hydraulic resistance in each circuit component.

By solving the system of equations in Equation 2b, the ratio of electric currents between each segment, hence the ratio of electric field strengths, was derived in Equation 3c:

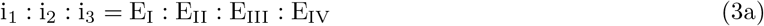

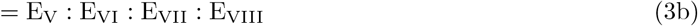

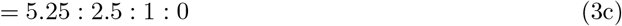

The similar analysis could be performed on the hydraulic equivalent circuit of HMEFC using the electricalhydraulic analogy^7^ (FIG. S.12.b). The interconnecting channels and main channels had width much bigger than height, thus, the hydraulic resistance can be calculated using the below equation^8^:

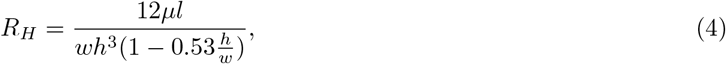

where *μ* is the fluid viscosity, *l* is the channel length, *w* is the channel width, and *h* is the channel height.

The hydraulic resistances in the 10 *μ*m-high interconnecting channel were much higher than the two 100 *μ*m-high, 2 mm-wide main channels were cells resided, limiting the advectional chemical transport. The flow rates in interconnecting channels and main channels differed by 18 orders as simulated by PSPICE analysis (TABLE. S.6).

Two dimensionless numbers such as Reynolds number and Péclet number can be used to characterize the fluid flow and chemical transport in the microfluidic system.

The Reynolds number, *ℜ*, is a dimensionless parameter to determine if the system is in the laminar or turbulent regime^9^. It is a measure of competition between the inertia force over the viscous force in the flow. In general, most microfluidic chips for cell studies involve Reynolds numbers much smaller than 1 and the flows in the lab-on-chip devices are observed to be laminar.

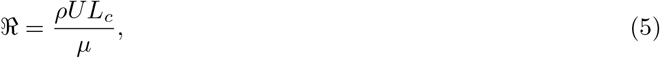

where *L_c_* is a characteristic length scale related to the flow field, *U* is the characteristic velocity, *ρ* is the density, and *μ* is the dynamic viscosity of the fluid. In HMEFC, the characteristic length was the hydrodynamic radius 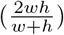, where *w* and *h* were width and height of microchannels.

The Péclet number describes the proportional relationship of chemical transport between convective fluxes and diffusive fluxes. In a system of high Péclet number, the diffusion is negligible whereas in low Péclet number system, the scalar (*i.e.,* solute concentration) distribution governed by diffusion follows the Fick’s Law^9^:

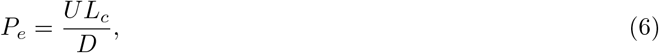

where *U* is the characteristic velocity, *L_c_* is the characteristic length, and *D* is the diffusion coefficient of the scalar.

In HMEFC, the Reynolds number and Péclet number were much lower in the interconnecting channels compared to those of the main channels, suggesting that the chemical transport through the interconnecting channels that would cause “cross-contamination” event was diffusion-limited.

#### 1.2 HMEFC simulation

The 3D design model HMEFC was exported from AutoCAD and imported into COMSOL Multiphysics 5.3 (COMSOL Inc, USA). Steady-state coupled simulations using three modules, creeping flow, chemical transport of diluted species, and electric currents, were performed. To correctly simulate the system, in-house measured material properties of the minimum essential media *α* (MEM*α*) for cell culture supplemented with 10% FBS medium were input. The material in the 3D model was set as water with density of 1002.9 Kg m^3^, electrical conductivity of 1.536 S m^−1^, dynamic shear viscosity of 0.946 mPa s, and relative permittivity of 80. The numerically simulated electric field ratio was 4.99: 2.45:1:0 in section I to IV and section V to VIII with limited chemical transport across the interconnecting channels as designed.

The boundary conditions were input accordingly for creeping flow, electric current, and chemical transport of diluted species as shown in FIG. 5. For creeping flow, 20 *μ*L min^−1^ flow rate was used. For chemical transport, 100 mol mm^−3^ of 40 kDa dextran was set to inflow in the top main channel while no dextran was flew in the bottom channel. The 40 kDa had a diffusion coefficient of 44.7 *μ*m^2^ s^−110^. For establishment of 300 V m^−1^ electric field, 485.5 *μ*A m^−2^ was set at the anode and electric potential of 0 V was set at the cathode.

The dcEFs in section I to IV and V to VIII were simulated as 4.99:2.45:1:0 (FIG. S.13.a) which were comparable to the theoretical calculation. The chemical distribution of the 40 kDa dextran was shown in FIG. S.13.b. By adjusting the scale of the colormap, the “cross-contamination” events where chemical leaches from one main channel to another through the interconnecting channels could be seen more clearly (FIG. S.13.c). In the double-layer design, it was more diffusion-dominant and the cross-contamination was limited. These results shows the HMEFCs could create multiple dcEFs without cross-contamination of chemicals, increasing the experimental throughput.

#### 1.3 Diffusion-limited chemical transport validation in HMEFC

The diffusion-limited chemical transport in HMEFC was visualized by flowing 40 kDa fluorescein-dextran (FD40, Sigma-Aldrich, USA) with U-251MG cells or 40 kDa tetramethylrhodamine-dextran (D1842, Thermo Fisher Scientific, USA) in Fluorobrite DMEM (Gibco, USA). The dyed media were loaded separately in two 2.5 mL syringes (Terumo, Japan) and the channels was primed at 20 *μ*L min^−1^ for 10 minutes before reducing to 20 *μ*L hr^−1^ (cell culture experiment flow rate), followed by capturing microscopy images with 10 minute time lapse epifluorescence microscopy over ten hours (Ti-E, Nikon, Japan).

As shown in FIG. S.14, the chemical transport was stable over the course of 10 hours at cell culture-relevant flow rates. Furthermore, when cells were injected in the top channel of the double-layer microfluidic chip, the cells were not able to pass through 10 *μ*m-height interconnecting channels and were restricted to its main channel. When different cell types were seeded in separate channels, the double-layer microchannel design could further increase the experimental throughput.

**FIG. S.13:**
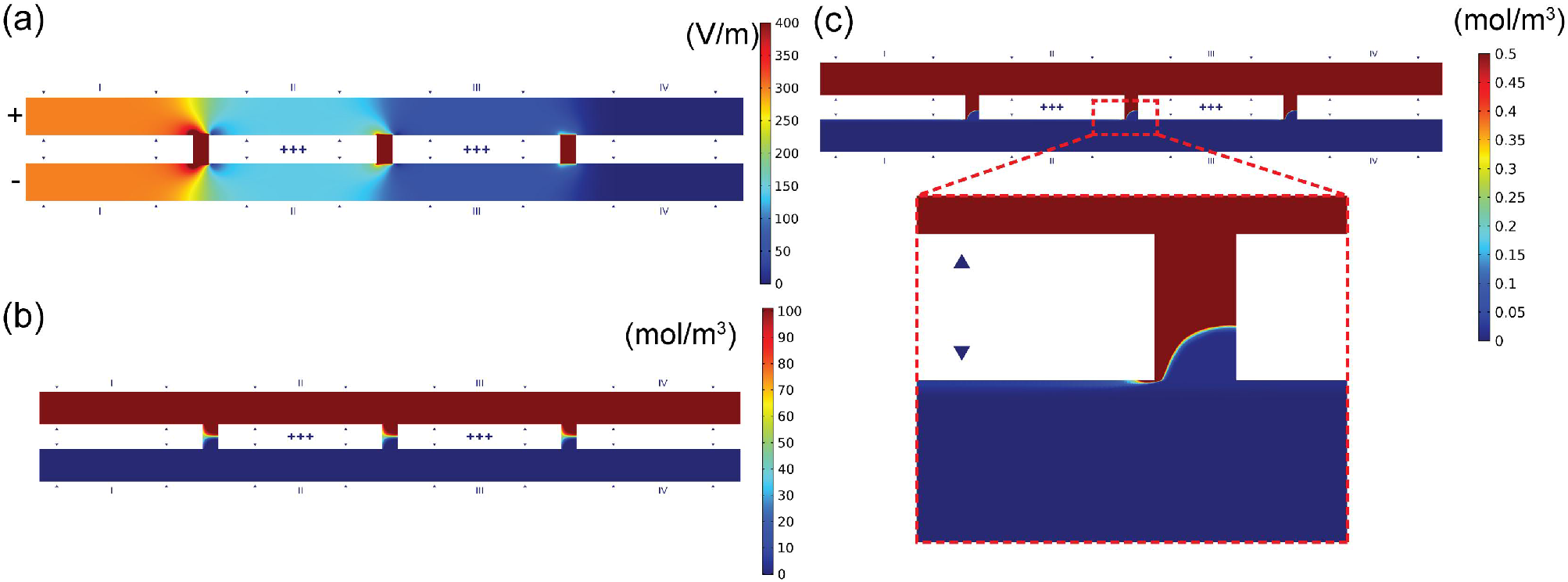
The numerical simulation results of the hybrid multiple electric field chip (HMEFC). (a) The electric field distribution in HMEFC. Application of a direct current electric field from the two outlets (nearest to section I) creates 4.99:2.45:1:0 stepwise electric fields for high throughout electrotaxis study; (b) The coupled chemical transport (convection and electrokinesis) simulated in HMEFC. 100 mol m^−3^ of 40 kDa dextran was flowing in the top channel and 0 mol m^−3^ was flowing in the bottom channel; (c) The coupled chemical transport simulated in HMEFC with the color scaled adjusted to show the “cross-contamination” event. The red dashed box shows the magnified region of an interconnecting channel. Only a minute amount of chemical diffuse through the interconnecting channel.

#### 1.4 HMEFC fabrication

A HMEFC was composed of a PMMA top reservoir, two PMMA components, and a PDMS chip (FIG. S.15). For high throughput experiments, another U-shaped PMMA agarose salt bridge made electrical conduction between two HMEFCs.

##### PDMS chip fabrication

In the double-layer HMEFC, the first layer was 10 *μ*m-high and the second layer was 100 *μ*m. The design was designed in AutoCAD and exported to KLayout. Several cross-shaped alignment markers were included in the design to assist alignment during making of the mold for double-layer microfluidic chip.

A 10 *μ*m-thick layer of photoresist (DWL-5, Micro Resist Technology, Germany) was spin-coated on a 100 mm silicon wafer and soft baked. The first design layer was directly written by the maskless lithographic writer (80 mJ cm^−2^, DL-1000, Nano Systems Solutions, Japan) and subsequently hard-baked. Next, without development, a layer of 100*μ*m-thick photoresist (SU-8 3050, MicroChem Corp, USA) was spin-coated on the wafer and soft baked. A chrome mask (CBL5006Du-AZPFS, Clean Surface Technology, Japan) for the second layer was fabricated using the maskless lithographic writer and the pattern was etched away by etchant (651826, Sigma Aldrich, USA). Using the alignment markers, the silicon wafer with the first-layer structures was aligned to the second layer structures on the chrome mask on the mask aligner (MA/BA6, SUSS MicroTec, Germany) and subsequently exposed (45 s of i-line UV irradiation). The unexposed photoresist was dissolved away in propylene glycol methyl ether acetate (PGMEA, Sigma Aldrich, USA) and the wafer was washed thoroughly with isopropanol, water, and dried with nitrogen gas. The wafer with microstructures was passivated by 1H, 1H, 2H, 2H-perfluorooctyltriethoxysilane (667420, Sigma-Aldrich, USA) in a vacuum dessicator.

To fabricate the PDMS devices, PDMS (Sylgard 184, Dow Corning, USA) monomer was mixed with curing agent at 10:1 ratio, degassed, and poured on the mold in a in-house made casing block made of polytetrafluo-roethylene. After degassing, the PDMS in the casting block was flanked with a piece of 15 mm-thick PMMA block to ensure the exact 4 mm-high final device with surface parallelism. The PDMS was cured under 60°C for at least 2 hours.

**FIG. S.14:**
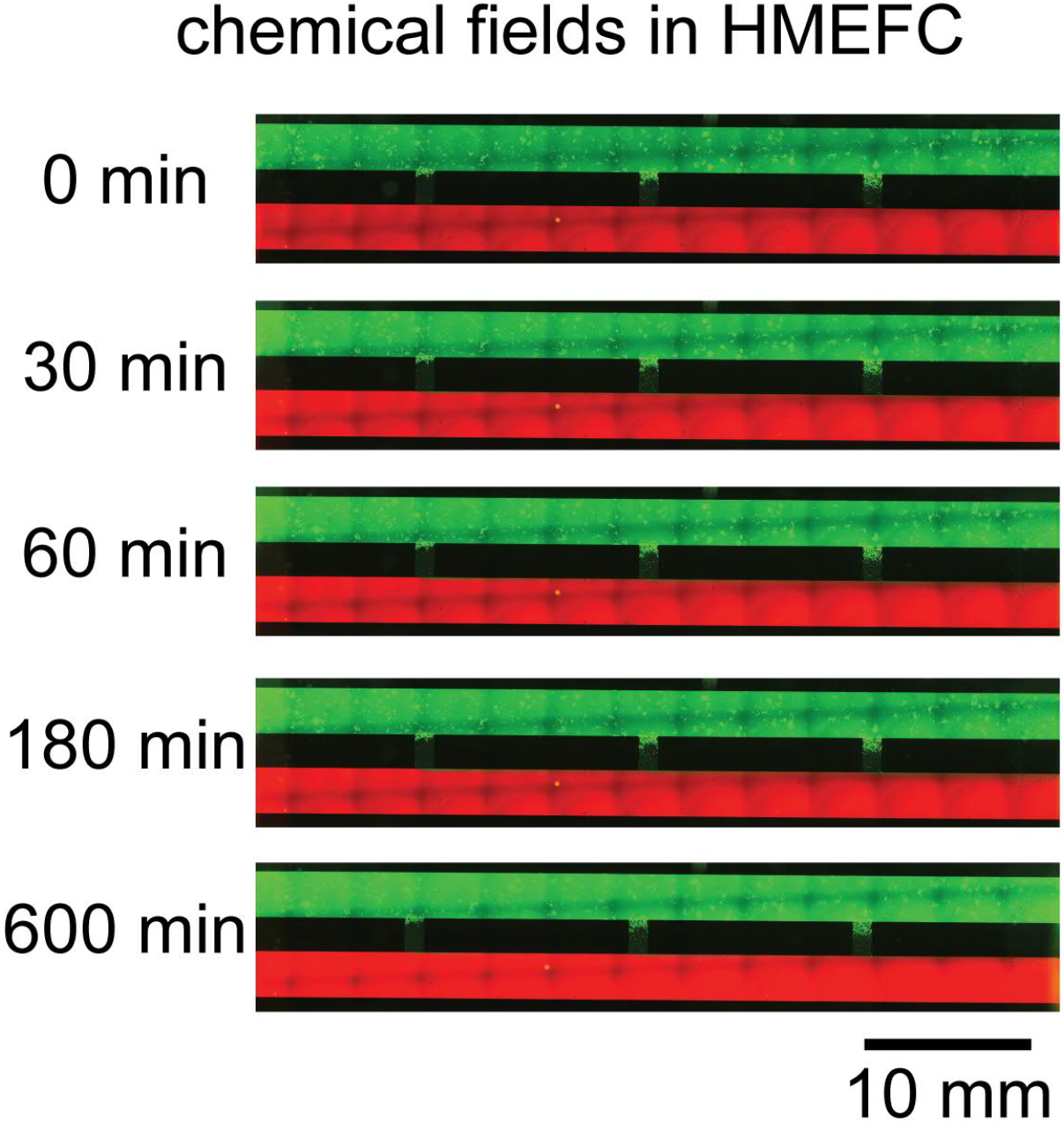
Chemical distribution in the hybrid multiple electric field chip (HMEFC) over 10 hours at 20 *μ* L hr ^−1^. 1 mM 40 kDa fluorescein-dextran with cells was flown in the top channel and 1 mM 40 kDa tetramethylrhodamine-dextran was flown in the bottom channel at 20 *μ*L hr^−1^. The chemical distribution was stable over a period of 10 hours.

After curing, the PDMS devices with the negative impression of the microstructures on the master mold was delaminated and cut into single devices appropriately. Inlet/outlet ports were punched with a 1 mm-diameter biopsy punch (BPP-10F, KAI Group, Japan). The port size around 1 mm was appropriate for biologist-friendly fluid manipulation using a 200 *μ*L pipet tip.

Each PDMS device was bonded to a piece of coverglass cleaned to complete the PDMS chip. To ensure surface quality and optical performance, high-precision borosilicate coverglasses (24×60 mm, 170 *±*5 *μ*mthick, No.1.5H, Marienfeld-Superior, Germany) were cleaned in a washing solution with ultrasound (1% TFD4, Franklab, France). The coverglasses were washed thoroughly with ultrapure water, dried, and disinfected with ultraviolet irradiation prior to bonding to PDMS devices using O_2_ plasma (AP-300, Nordson MARCH, USA).

###### PMMA component fabrication

3D microfluidic components could be easily and rapidly fabricated using PMMA. In HMEFC, 4 PMMA components were used including the medium inlet reservoir (component A, FIG. S.15.a), outlet/electrical stimulation interface (component B, FIG. S.15.a), top reservoir, and salt bridge connector. The medium inlet reservoir, outlet/electrical stimulation interface, and top reservoir components are composed of four layers of 2 mm PMMA substrates.

The designs were created in AutoCAD software and the patterns were cut on a piece of 2 mm casted PMMA substrate (casted acrylic sheet K, Kanase, Japan) with a CO_2_ laser cutter (VLS3.50, Universal Laser Systems, USA).

3D microfluidic components can be easily fabricated by stacking the laser-cut PMMA pieces and bonding them under high temperature and pressure. The layers were aligned by hand and flanked between two pieces of 2 mm-thick, 100 mm Tempax float glass wafers (Schott AG, Germany) on a force-controlled programmable hot press (G-12RS2000, Orihara Industrial co., ltd, Japan). The PMMA pieces were heated above its glass transition temperature with pressure to form leakage free 3D microfluidic components (118°C, 500 N, 30 min).

**FIG. S.15:**
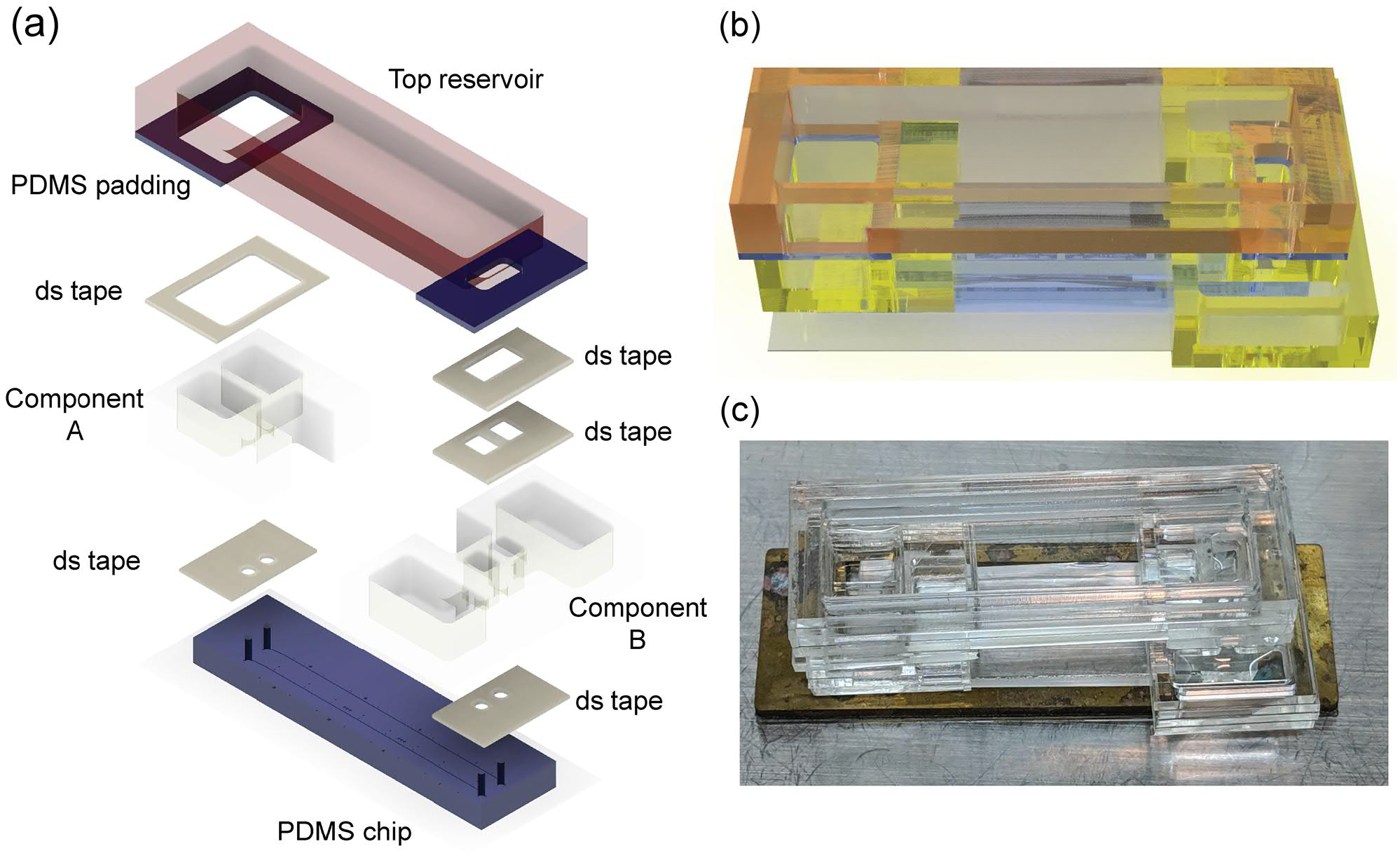
The assembled hybrid multiple electric field chip (HMEFC). (a) The exploded view of the components for HMEFC. ds tape: double-sided tape; (b) The 3D rendered model of the the assembled device; (c) The photoimage of the assembled device for experiment. A copper based holder was used to prevent damage to the coverglass.

###### Facile reversible-bonding between PMMA components and PDMS chip

A dual energy double-sided tape (85 *μ*m, No. 5302A, Nitto, Japan) was patterned using the CO_2_ laser scriber. On no.5302A double-sided tapes, a silicone-type adhesive and an acrylic-type adhesive were overlaid on the two sides of a poly(ethylene terephthalate) (PET) substrate. The silicone-type pressure sensitive adhesive adhered to silicone rubber, while the acrylic-type adhesive affixed to plastic, glass, and metal surfaces^3,4^. The double-sided tape provided an easy and facile way to bond between PDMS and PMMA, an interface typically hard to join. Thus, double-sided tape provides the advantage of high spatial precision of PDMS microfluidics and rapid 3D prototyping of PMMA microfluidics. Using two pieces of patterned double-sided tape, the PMMA component A and B were aligned and affixed to the PDMS chip (FIG. S.15.a).

In addition, to avoid breakage and flexing of thin coverglass, a copper holder was used to provide solid support (FIG. S.15.c). The 2 mm-thick copper holder was made of wire electrical discharge machining (wireEDM) and backed with 0.5 mm PDMS sheet. The PDMS was attached reversibly to the coverglass of the PDMS microfluidic chip.

###### PMMA top reservoir fabrication

To balance the hydrostatic pressure between the inlet and outlet to prevent Poiseuille flow that could displace cells due to hydrostatic pressure, a top reservoir was placed to connect component A and component B. The top reservoir was fabricated by 4 layers of 2 mm-thick PMMA fabricated also with laser cutting and thermal bonding. The top reservoir was made to reversibly bond to PMMA components by affixing 2 mm-thick PDMS paddings using double sided tapes. Openings were cut by a utility knife. Afterward the PDMS part of the top reservoir was affixed to the top of the component A and B using patterned double-sided tapes, completing the assembly of the device. The zoomed-in view of the components and 3D render of the hybrid HMEFC were shown in FIG. S.15.b & FIG. S.15.c.

## References

1 O. Gallego, “Nonsurgical treatment of recurrent glioblastoma,” Current oncology 22, e273 (2015).

2 S. Mallick, R. Benson, A. Hakim, and G. K. Rath, “Management of glioblastoma after recurrence: A changing paradigm,” Journal of the Egyptian National Cancer Institute 28, 199–210 (2016).

3 A. Hara, T. Kanayama, K. Noguchi, A. Niwa, M. Miyai, M. Kawaguchi, K. Ishida, Y. Hatano, M. Niwa, and H. Tomita, “Treatment strategies based on histological targets against invasive and resistant glioblastoma.” Journal of oncology 2019, 2964783 (2019).

4 M. Alieva, V. Leidgens, M. J. Riemenschneider, C. A. Klein, P. Hau, and J. van Rheenen, “Intravital imaging of glioma border morphology reveals distinctive cellular dynamics and contribution to tumor cell invasion,” Scientific reports 9, 2054 (2019).

5 A. Dirkse, A. Golebiewska, T. Buder, P. V. Nazarov, A. Muller, S. Poovathingal, N. H. Brons, S. Leite, N. Sauvageot, D. Sarkisjan, et al., “Stem cell-associated heterogeneity in glioblastoma results from intrinsic tumor plasticity shaped by the microenvironment,” Nature communications 10, 1787 (2019).

6 D. Hambardzumyan and G. Bergers, “Glioblastoma: Defining tumor niches.” Trends in cancer 1, 252–265 (2015).

7 H.-F. Tsai, A. Trubelja, A. Q. Shen, and G. Bao, “Tumour-ona-chip: microfluidic models of tumour morphology, growth and microenvironment.” Journal of the Royal Society, Interface 14, 20170137 (2017).

8 I. Manini, F. Caponnetto, A. Bartolini, T. Ius, L. Mariuzzi, C. Di Loreto, A. P. Beltrami, and D. Cesselli, “Role of microenvironment in glioma invasion: what we learned from in vitro models,” International journal of molecular sciences 19, 147 (2018).

9 W. Tomaszewski, L. Sanchez-Perez, T. F. Gajewski, and J. H. Sampson, “Brain tumor microenvironment and host state: Implications for immunotherapy,” Clinical Cancer Research 25, 4202–4210 (2019).

10 M. Diksin, S. J. Smith, and R. Rahman, “The molecular and phenotypic basis of the glioma invasive perivascular niche,” International journal of molecular sciences 18, 2342 (2017).

11 A. Hatoum, R. Mohammed, and O. Zakieh, “The unique invasiveness of glioblastoma and possible drug targets on extracellular matrix,” Cancer Management and Research 11, 1843 (2019).

12 C. G. Hales and S. Pockett, “The relationship between local field potentials (lfps) and the electromagnetic fields that give rise to them,” Frontiers in Systems Neuroscience 8, 233 (2014).

13 L. Cao, D. Wei, B. Reid, S. Zhao, J. Pu, T. Pan, E. N. Yamoah, and M. Zhao, “Endogenous electric currents might guide rostral migration of neuroblasts,” EMBO reports 14, 184–190 (2013).

14 J. H. Lee, J. E. Lee, J. Y. Kahng, S. H. Kim, J. S. Park, S. J. Yoon, J.-Y. Um, W. K. Kim, J.-K. Lee, J. Park, et al., “Human glioblastoma arises from subventricular zone cells with low-level driver mutations,” Nature 560, 243–247 (2018).

15 Y.-J. Huang, G. Hoffmann, B. Wheeler, P. Schiapparelli, A. Quinones-Hinojosa, and P. Searson, “Cellular microenvironment modulates the galvanotaxis of brain tumor initiating cells.” Scientific reports 6, 21583 (2016).

16 Y.-J. Huang, P. Schiapparelli, K. Kozielski, J. Green, E. Lavell, H. Guerrero-Cazares, A. Quinones-Hinojosa, and P. Searson, “Electrophoresis of cell membrane heparan sulfate regulates galvanotaxis in glial cells.” Journal of cell science 130, 2459–2467 (2017).

17 H.-F. Tsai, K. Toda-Peters, and A. Q. Shen, “Glioblastoma adhesion in a quick-fit hybrid microdevice,” Biomedical microdevices 21, 30 (2019).

18 Y.-S. Sun, “Direct-current electric field distribution in the brain for tumor treating field applications: A simulation study.” Computational and mathematical methods in medicine 2018, 3829768 (2018).

19 J. G. Lyon, S. L. Carroll, N. Mokarram, and R. V. Bellamkonda, “Electrotaxis of glioblastoma and medulloblastoma spheroidal aggregates.” Scientific reports 9, 5309 (2019).

20 B. E. Goodman, “Channels active in the excitability of nerves and skeletal muscles across the neuromuscular junction: basic function and pathophysiology,” Advances in physiology education 32, 127–135 (2008).

21 W. Taft and R. DeLorenzo, “Regulation of calcium channels in brain: implications for the clinical neurosciences.” The Yale journal of biology and medicine 60, 99 (1987).

22 E. M. Kawamoto, C. Vivar, and S. Camandola, “Physiology and pathology of calcium signaling in the brain,” Frontiers in pharmacology 3, 61 (2012).

23 N. Robil, F. Petel, M.-C. Kilhoffer, and J. Haiech, “Glioblas-toma and calcium signaling-analysis of calcium toolbox expression,” Int. J. Dev. Biol 59, 407–415 (2015).

24 F. B. Morrone, M. P. Gehring, and N. F. Nicoletti, “Calcium channels and associated receptors in malignant brain tumor therapy.” Molecular pharmacology 90, 403–409 (2016).

25 J. Pollak, K. G. Rai, C. C. Funk, S. Arora, E. Lee, J. Zhu, N. D. Price, P. J. Paddison, J.-M. Ramirez, and R. C. Rostomily, “Ion channel expression patterns in glioblastoma stem cells with functional and therapeutic implications for malignancy.” PloS one 12, e0172884 (2017).

26 N. N. Phan, C.-Y. Wang, C.-F. Chen, Z. Sun, M.-D. Lai, and Y.-C. Lin, “Voltage-gated calcium channels: Novel targets for cancer therapy.” Oncology letters 14, 2059–2074 (2017).

27 A. Maklad, A. Sharma, and I. Azimi, “Calcium signaling in brain cancers: Roles and therapeutic targeting.” Cancers 11 (2019), 10.3390/cancers11020145.

28 B. Song, Y. Gu, J. Pu, B. Reid, Z. Zhao, and M. Zhao, “Application of direct current electric fields to cells and tissues in vitro and modulation of wound electric field in vivo,” Nature protocols 2, 1479 (2007).

29 N. Tandon, C. Cannizzaro, P.-H. G. Chao, R. Maidhof, A. Marsano, H. T. H. Au, M. Radisic, and G. Vunjak-Novakovic, “Electrical stimulation systems for cardiac tissue engineering,” Nature protocols 4, 155 (2009).

30 H.-F. Tsai, J. Gajda, T. F. Sloan, A. Rares, and A. Q. Shen, “Usiigaci: Instance-aware cell tracking in stain-free phase contrast microscopy enabled by machine learning,” SoftwareX 9, 230–237 (2019).

31 L. Kim, Y.-C. Toh, J. Voldman, and H. Yu, “A practical guide to microfluidic perfusion culture of adherent mammalian cells.” Lab on a chip 7, 681–694 (2007).

32 G. A. Villalona, B. Udelsman, D. R. Duncan, E. McGillicuddy, R. F. Sawh-Martinez, N. Hibino, C. Painter, T. Mirensky, B. Erickson, T. Shinoka, et al., “Cell-seeding techniques in vascular tissue engineering,” Tissue Engineering Part B: Reviews 16, 341–350 (2010).

33 P. M. Reynolds, C. H. Rasmussen, M. Hansson, M. Dufva, M. O. Riehle, and N. Gadegaard, “Controlling fluid flow to improve cell seeding uniformity,” PloS one 13, e0207211 (2018).

34 J. T. Parsons, A. R. Horwitz, and M. A. Schwartz, “Cell adhesion: integrating cytoskeletal dynamics and cellular tension,” Nature reviews Molecular cell biology 11, 633 (2010).

35 C. De Pascalis and S. Etienne-Manneville, “Single and collective cell migration: the mechanics of adhesions,” Molecular biology of the cell 28, 1833–1846 (2017).

36 C. W. Brennan, R. G. W. Verhaak, A. McKenna, B. Cam-pos, H. Noushmehr, S. R. Salama, S. Zheng, D. Chakravarty, J. Z. Sanborn, S. H. Berman, R. Beroukhim, B. Bernard, C.-J. Wu, G. Genovese, I. Shmulevich, J. Barnholtz-Sloan, L. Zou, R. Vegesna, S. A. Shukla, G. Ciriello, W. K. Yung, W. Zhang, C. Sougnez, T. Mikkelsen, K. Aldape, D. D. Bigner, E. G. Van Meir, M. Prados, A. Sloan, K. L. Black, J. Eschbacher, G. Finocchiaro, W. Friedman, D. W. Andrews, A. Guha, M. Iacocca, B. P. O’Neill, G. Foltz, J. Myers, D. J. Weisenberger, R. Penny, R. Kucherlapati, C. M. Perou, D. N. Hayes, R. Gibbs, M. Marra, G. B. Mills, E. Lander, P. Spellman, R. Wilson, C. Sander, J. Weinstein, M. Meyerson, S. Gabriel, P. W. Laird, D. Haussler, G. Getz, L. Chin, and T. R. Network, “The somatic genomic landscape of glioblastoma.” Cell 155, 462–477 (2013).

37 L. N. Kiseleva, A. V. Kartashev, N. L. Vartanyan, A. A. Pinevich, and M. P. Samoilovich, “A172 and t98g cell lines characteristics,” Cell and Tissue Biology 10, 341–348 (2016).

38 B. W. Stringer, J. Bunt, B. W. Day, G. Barry, P. R. Jamieson, K. S. Ensbey, Z. C. Bruce, K. Goasdoué, H. Vidal, S. Charmsaz, F. M. Smith, L. T. Cooper, M. Piper, A. W. Boyd, and L. J. Richards, “Nuclear factor one b (nfib) encodes a subtype-specific tumour suppressor in glioblastoma.” Oncotarget 7, 29306–29320 (2016).

39 F. Li, T. Chen, S. Hu, J. Lin, R. Hu, and H. Feng, “Superoxide mediates direct current electric field-induced directional migration of glioma cells through the activation of akt and erk.” PloS one 8, e61195 (2013).

40 R. Fridman, M. C. Kibbey, L. S. Royce, M. Zain, M. Sweeney, D. L. Jicha, J. R. Yannelli, G. R. Martin, and H. K. Kleinman, “Enhanced tumor growth of both primary and established human and murine tumor cells in athymic mice after coinjection with matrigel.” Journal of the National Cancer Institute 83, 769–774 (1991).

41 R. O. Hynes, “Integrins: versatility, modulation, and signaling in cell adhesion.” Cell 69, 11–25 (1992).

42 M. S. Diamond and T. A. Springer, “The dynamic regulation of integrin adhesiveness.” Current biology: CB 4, 506–517 (1994).

43 S. McLaughlin and M. M. Poo, “The role of electro-osmosis in the electric-field-induced movement of charged macromolecules on the surfaces of cells,” Biophysical Journal 34, 85–93 (1981).

44 F. X. Hart, “Integrins may serve as mechanical transducers for low-frequency electric fields,” Bioelectromagnetics 27, 505–508 (2006).

45 K. Zhu, Y. Takada, K. Nakajima, Y. Sun, J. Jiang, Y. Zhang, Q. Zeng, Y. Takada, and M. Zhao, “Expression of integrins to control migration direction of electrotaxis,” The FASEB Journal, fj–201802657R (2019).

46 Y. Taniguchi, H. Ido, N. Sanzen, M. Hayashi, R. Sato-Nishiuchi, S. Futaki, and K. Sekiguchi, “The c-terminal region of laminin beta chains modulates the integrin binding affinities of laminins.” The Journal of biological chemistry 284, 7820–7831 (2009).

47 T. Miyazaki, S. Futaki, H. Suemori, Y. Taniguchi, M. Yamada, M. Kawasaki, M. Hayashi, H. Kumagai, N. Nakatsuji, K. Sekiguchi, and E. Kawase, “Laminin e8 fragments support efficient adhesion and expansion of dissociated human pluripotent stem cells.” Nature communications 3, 1236 (2012).

48 J. Cha and P. Kim, “Biomimetic strategies for the glioblastoma microenvironment,” Frontiers in Materials 4, 45 (2017).

49 L. W. Lau, R. Cua, M. B. Keough, S. Haylock-Jacobs, and V.W. Yong, “Pathophysiology of the brain extracellular matrix: a new target for remyelination,” Nature Reviews Neuroscience 14, 722–729 (2013).

50 G. J. Baker, V. N. Yadav, S. Motsch, C. Koschmann, A.-A. Calinescu, Y. Mineharu, S. I. Camelo-Piragua, D. Orringer, S. Bannykh, W. S. Nichols, et al., “Mechanisms of glioma formation: iterative perivascular glioma growth and invasion leads to tumor progression, vegf-independent vascularization, and resistance to antiangiogenic therapy,” Neoplasia 16, 543–561 (2014).

51 H.-F. Tsai, S.-W. Peng, C.-Y. Wu, H.-F. Chang, and J.-Y. Cheng, “Electrotaxis of oral squamous cell carcinoma cells in a multiple-electric-field chip with uniform flow field.” Biomicrofluidics 6, 34116 (2012).

52 A. Soeda, A. Hara, T. Kunisada, S.-i. Yoshimura, T. Iwama, and D. M. Park, “The evidence of glioblastoma heterogeneity.” Scientific reports 5, 7979 (2015).

53 Z. Zakaria, A. Tivnan, L. Flanagan, D. W. Murray, M. Salvucci, B. W. Stringer, B. W. Day, A. W. Boyd, D. Kögel, M. Rehm, D. F. O’Brien, A. T. Byrne, and J. H. M. Prehn, “Patientderived glioblastoma cells show significant heterogeneity in treatment responses to the inhibitor-of-apoptosis-protein antagonist birinapant.” British journal of cancer 114, 188–198 (2016).

54 S. Chen, T. Le, B. A. C. Harley, and P. I. Imoukhuede, “Characterizing glioblastoma heterogeneity via single-cell receptor quantification.” Frontiers in bioengineering and biotechnology 6, 92 (2018).

55 S. L. Perrin, M. S. Samuel, B. Koszyca, M. P. Brown, L. M. Ebert, M. Oksdath, and G. A. Gomez, “Glioblastoma heterogeneity and the tumour microenvironment: implications for preclinical research and development of new treatments.” Biochemical Society transactions 47, 625–638 (2019).

56 N. Schäfer, G. H. Gielen, L. Rauschenbach, S. Kebir, A. Till, R. Reinartz, M. Simon, P. Niehusmann, C. Kleinschnitz, U. Herrlinger, T. Pietsch, B. Scheffler, and M. Glas, “Longitudinal heterogeneity in glioblastoma: moving targets in recurrent versus primary tumors.” Journal of translational medicine 17, 96 (2019).

57 E. K. Onuma and S. W. Hui, “A calcium requirement for electric field-induced cell shape changes and preferential orientation.” Cell calcium 6, 281–292 (1985).

58 L. Tung, N. Sliz, and M. R. Mulligan, “Influence of electrical axis of stimulation on excitation of cardiac muscle cells.” Circulation research 69, 722–730 (1991).

59 M. Zhao, J. V. Forrester, and C. D. McCaig, “A small, physiological electric field orients cell division,” Proceedings of the National Academy of Sciences of the United States of America 96, 4942–4946 (1999).

60 T. A. Banks, P. S. B. Luckman, J. E. Frith, and J. J. Cooper-White, “Effects of electric fields on human mesenchymal stem cell behaviour and morphology using a novel multichannel device.” Integrative biology: quantitative biosciences from nano to macro 7, 693–712 (2015).

61 G. H. Markx, “The use of electric fields in tissue engineering: A review.” Organogenesis 4, 11–17 (2008).

62 L. P. da Silva, S. C. Kundu, R. L. Reis, and V. M. Correlo, “Electric phenomenon: A disregarded tool in tissue engineering and regenerative medicine.” Trends in biotechnology (2019), 10.1016/j.tibtech.2019.07.002.

63 I. Minoura and E. Muto, “Dielectric measurement of individual microtubules using the electroorientation method.” Biophysical journal 90, 3739–3748 (2006).

64 T. Kim, M.-T. Kao, E. F. Hasselbrink, and E. Meyhöfer, “Active alignment of microtubules with electric fields.” Nano letters 7, 211–217 (2007).

65 R. Nuccitelli and T. Smart, “Extracellular calcium levels strongly influence neural crest cell galvanotaxis,” The Biological Bulletin 176, 130–135 (1989).

66 L. J. Shanley, P. Walczysko, M. Bain, D. J. MacEwan, and M. Zhao, “Influx of extracellular ca2+ is necessary for electrotaxis in dictyostelium,” J Cell Sci 119, 4741–4748 (2006).

67 P. Borys, “The role of passive calcium influx through the cell membrane in galvanotaxis,” Cellular & molecular biology letters 18, 187 (2013).

68 L. Guo, C. Xu, D. Li, X. Zheng, J. Tang, J. Bu, H. Sun, Z. Yang, W. Sun, and X. Yu, “Calcium ion flow permeates cells through socs to promote cathode-directed galvanotaxis,” PloS one 10, e0139865 (2015).

69 R. Babona-Pilipos, N. Liu, A. Pritchard-Oh, A. Mok, D. Badawi, M. Popovic, and C. Morshead, “Calcium influx differentially regulates migration velocity and directedness in response to electric field application,” Experimental cell research 368, 202–214 (2018).

70 M. B. Mcferrin and H. Sontheimer, “A role for ion channels in glioma cell invasion,” Neuron glia biology 2, 39–49 (2006).

71 H. Sontheimer, “An unexpected role for ion channels in brain tumor metastasis,” Experimental biology and medicine 233, 779–791 (2008).

72 R. J. Molenaar, “Ion channels in glioblastoma,” ISRN neurology 2011(2011).

73 R. Wang, C. I. Gurguis, W. Gu, E. A. Ko, I. Lim, H. Bang, T. Zhou, and J.-H. Ko, “Ion channel gene expression predicts survival in glioma patients.” Scientific reports 5, 11593 (2015).

74 H.-Y. Wang, J.-Y. Li, X. Liu, X.-Y. Yan, W. Wang, F. Wu, T.-Y. Liang, F. Yang, H.-M. Hu, H.-X. Mao, Y.-W. Liu, and S.-Z. Zhang, “A three ion channel genes-based signature predicts prognosis of primary glioblastoma patients and reveals a chemotherapy sensitive subtype.” Oncotarget 7, 74895–74903 (2016).

75 N. C. K. Valerie, B. Dziegielewska, A. S. Hosing, E. Augustin, L. S. Gray, D. L. Brautigan, J. M. Larner, and J. Dziegielewski, “Inhibition of t-type calcium channels disrupts akt signaling and promotes apoptosis in glioblastoma cells.” Biochemical pharmacology 85, 888–897 (2013).

76 Y. Zhang, N. Cruickshanks, F. Yuan, B. Wang, M. Pahuski, J. Wulfkuhle, I. Gallagher, A. F. Koeppel, S. Hatef, C. Papanicolas, J. Lee, E. E. Bar, D. Schiff, S. D. Turner, E. F. Petricoin, L. S. Gray, and R. Abounader, “Targetable t-type calcium channels drive glioblastoma.” Cancer research 77, 3479–3490 (2017).

77 B. Hille, Ion Channels of Excitable Membranes, Third Edition, 3rd ed. (Sinauer Associates, 2011).

78 G. R. Tibbs, D. J. Posson, and P. A. Goldstein, “Voltage-gated ion channels in the pns: Novel therapies for neuropathic pain?” Trends in pharmacological sciences 37, 522–542 (2016).

79 M. Strickland, B. Yacoubi-Loueslati, B. Bouhaouala-Zahar, S. L. F. Pender, and A. Larbi, “Relationships between ion channels, mitochondrial functions and inflammation in human aging.” Frontiers in physiology 10, 158 (2019).

80 D. R. Trollinger, R. R. Isseroff, and R. Nuccitelli, “Calcium channel blockers inhibit galvanotaxis in human keratinocytes.” Journal of cellular physiology 193, 1–9 (2002).

81 M. Aonuma, T. Kadono, and T. Kawano, “Inhibition of anodic galvanotaxis of green paramecia by t-type calcium channel inhibitors.” Zeitschrift fur Naturforschung. C, Journal of biosciences 62, 93–102 (2007).

82 Y. Atsuta, R. R. Tomizawa, M. Levin, and C. J. Tabin, “L-type voltage-gated ca2+ channel cav1.2 regulates chondrogenesis during limb development,” Proceedings of the National Academy of Sciences of the United States of America 116, 21592–21601 (2019).

83 L. A. Pardo, C. Contreras-Jurado, M. Zientkowska, F. Alves, and W. Stühmer, “Role of voltage-gated potassium channels in cancer,” The Journal of membrane biology 205, 115–124 (2005).

84 H. Wulff, N. A. Castle, and L. A. Pardo, “Voltage-gated potassium channels as therapeutic targets,” Nature reviews Drug discovery 8, 982 (2009).

85 L. Y. Jan and Y. N. Jan, “Voltage-gated potassium channels and the diversity of electrical signalling,” The Journal of physiology 590, 2591–2599 (2012).

86 X. Huang and L. Y. Jan, “Targeting potassium channels in cancer,” J Cell Biol 206, 151–162 (2014).

87 A. Grizel, G. Glukhov, and O. Sokolova, “Mechanisms of activation of voltage-gated potassium channels,” Acta Naturae 6(2014).

88 N. Comes, A. Serrano-Albarras, J. Capera, C. Serrano-Novillo, E. Condom, S. R. y Cajal, J. C. Ferreres, and A. Felipe, “Involvement of potassium channels in the progression of cancer to a more malignant phenotype,” Biochimica et Biophysica Acta (BBA)-Biomembranes 1848, 2477–2492 (2015).

89 D. M. Kim and C. M. Nimigean, “Voltage-gated potassium channels: a structural examination of selectivity and gating,” Cold Spring Harbor perspectives in biology 8, a029231 (2016).

90 G. Zhang, M. Edmundson, V. Telezhkin, Y. Gu, X. Wei, P. J. Kemp, and B. Song, “The role of kv1.2 channel in electrotaxis cell migration.” Journal of cellular physiology 231, 1375–1384 (2016).

91 K.-I. Nakajima, K. Zhu, Y.-H. Sun, B. Hegyi, Q. Zeng, C. J. Murphy, J. V. Small, Y. Chen-Izu, Y. Izumiya, J. M. Penninger, and M. Zhao, “Kcnj15/kir4.2 couples with polyamines to sense weak extracellular electric fields in galvanotaxis.” Nature communications 6, 8532 (2015).

92 J. Morokuma, D. Blackiston, D. S. Adams, G. Seebohm, B. Trimmer, and M. Levin, “Modulation of potassium channel function confers a hyperproliferative invasive phenotype on embryonic stem cells.” Proceedings of the National Academy of Sciences of the United States of America 105, 16608–16613 (2008).

93 M. E. Laniado, S. P. Fraser, and M. B. Djamgoz, “Voltagegated k+ channel activity in human prostate cancer cell lines of markedly different metastatic potential: Distinguishing characteristics of pc-3 and lncap cells,” The Prostate 46, 262–274 (2001).

94 E. Venturini, L. Leanza, M. Azzolini, S. Kadow, A. Mattarei, M. Weller, G. Tabatabai, M. J. Edwards, M. Zoratti, C. Paradisi, et al., “Targeting the potassium channel kv1. 3 kills glioblastoma cells,” Neurosignals 25, 26–38 (2017).

95 S. I. McDonough, L. M. Boland, I. M. Mintz, and B. P. Bean, “Interactions among toxins that inhibit n-type and p-type calcium channels.” The Journal of general physiology 119, 313–328 (2002).

96 A. Lacampagne, F. Gannier, J. Argibay, D. Garnier, and J. Y. Le Guennec, “The stretch-activated ion channel blocker gadolinium also blocks l-type calcium channels in isolated ventricular myocytes of the guinea-pig.” Biochimica et biophysica acta 1191, 205–208 (1994).

97 P. Turlapaty, R. Vary, and J. A. Kaplan, “Nicardipine, a new intravenous calcium antagonist: a review of its pharmacology, pharmacokinetics, and perioperative applications.” Journal of cardiothoracic anesthesia 3, 344–355 (1989).

98 J. M. McIntosh, B. M. Olivera, L. J. Cruz, and W. R. Gray, “Gamma-carboxyglutamate in a neuroactive toxin.” The Journal of biological chemistry 259, 14343–14346 (1984).

99 B. M. Olivera, G. P. Miljanich, J. Ramachandran, and M. E. Adams, “Calcium channel diversity and neurotransmitter release: the omega-conotoxins and omega-agatoxins.” Annual review of biochemistry 63, 823–867 (1994).

100 P. M. Hinkle, P. A. Kinsella, and K. C. Osterhoudt, “Cadmium uptake and toxicity via voltage-sensitive calcium channels.” The Journal of biological chemistry 262, 16333–16337 (1987).

101 B. Xu, S. Chen, Y. Luo, Z. Chen, L. Liu, H. Zhou, W. Chen, T. Shen, X. Han, L. Chen, and S. Huang, “Calcium signaling is involved in cadmium-induced neuronal apoptosis via induction of reactive oxygen species and activation of mapk/mtor network.” PloS one 6, e19052 (2011).

102 R. C. Foehring, P. G. Mermelstein, W. J. Song, S. Ulrich, and D. J. Surmeier, “Unique properties of r-type calcium currents in neocortical and neostriatal neurons.” Journal of neurophysiology 84, 2225–2236 (2000).

103 C. Wormuth, A. Lundt, C. Henseler, R. Müller, K. Broich, A. Papazoglou, and M. Weiergräber, “Review: Cav2.3 r-type voltage-gated ca2+ channels functional implications in convulsive and non-convulsive seizure activity,” The open neurology journal 10, 99–126 (2016).

104 N. Nejatbakhsh and Z.-p. Feng, “Calcium binding protein-mediated regulation of voltage-gated calcium channels linked to human diseases.” Acta pharmacologica Sinica 32, 741–748 (2011).

105 M. Ben-Johny and D. T. Yue, “Calmodulin regulation (calmodulation) of voltage-gated calcium channels,” The Journal of general physiology 143, 679–692 (2014).

106 B. A. Simms and G. W. Zamponi, “Neuronal voltage-gated calcium channels: structure, function, and dysfunction,” Neuron 82, 24–45 (2014).

107 C.-M. Tang, F. Presser, and M. Morad, “Amiloride selectively blocks the low threshold (t) calcium channel,” Science 240, 213–215 (1988).

108 R. L. Kraus, Y. Li, Y. Gregan, A. L. Gotter, V. N. Uebele, S. V. Fox, S. M. Doran, J. C. Barrow, Z.-Q. Yang, T. S. Reger, K. S. Koblan, and J. J. Renger, “In vitro characterization of t-type calcium channel antagonist tta-a2 and in vivo effects on arousal in mice.” The Journal of pharmacology and experimental therapeutics 335, 409–417 (2010).

109 E. Chavez-Colorado, Z. Herrera-Carrillo, and J. C. Gomora, “Blocking of t-type calcium channels by tta-a2 reveals a conservative binding site for state-dependent antagonists,” Biophysical Journal 110, 439a–440a (2016).

110 J. F. Faivre, T. P. Calmels, S. Rouanet, J. L. Javré, B. Cheval, and A. Bril, “Characterisation of kv4.3 in hek293 cells: comparison with the rat ventricular transient outward potassium current.” Cardiovascular research 41, 188–199 (1999).

111 N. Hatano, S. Ohya, K. Muraki, W. Giles, and Y. Imaizumi, “Dihydropyridine ca2+ channel antagonists and agonists block kv4.2, kv4.3 and kv1.4 k+ channels expressed in hek293 cells.” British journal of pharmacology 139, 533–544 (2003).

112 T. Kimm and B. P. Bean, “Inhibition of a-type potassium current by the peptide toxin snx-482.” The Journal of neuroscience: the official journal of the Society for Neuroscience 34, 9182–9189 (2014).

113 H. Vacher, M. Alami, M. Crest, L. D. Possani, P. E. Bougis, and M.-F. Martin-Eauclaire, “Expanding the scorpion toxin alphaktx 15 family with ammtx3 from androctonus mauretanicus.” European journal of biochemistry 269, 6037–6041 (2002).

114 J. K. Maffie, E. Dvoretskova, P. E. Bougis, M.-F. Martin-Eauclaire, and B. Rudy, “Dipeptidyl-peptidase-like-proteins confer high sensitivity to the scorpion toxin ammtx3 to kv4mediated a-type k+ channels.” The Journal of physiology 591, 2419–2427 (2013).

115 P. E. Bougis and M.-F. Martin-Eauclaire, “Shal-type (kv4.x) potassium channel pore blockers from scorpion venoms.” Sheng li xue bao: [Acta physiologica Sinica] 67, 248–254 (2015).

116 S. Thompson, “Aminopyridine block of transient potassium current.” The Journal of general physiology 80, 1–18 (1982).

117 T. Takahashi, “Inward rectification in neonatal rat spinal motoneurones.” The Journal of physiology 423, 47–62 (1990).

118 T. Ishikawa, Y. Nakamura, N. Saitoh, W.-B. Li, S. Iwasaki, and T. Takahashi, “Distinct roles of kv1 and kv3 potassium channels at the calyx of held presynaptic terminal.” The Journal of neuroscience: the official journal of the Society for Neuroscience 23, 10445–10453 (2003).

119 S. Gründer and X. Chen, “Structure, function, and pharmacology of acid-sensing ion channels (asics): focus on asic1a,” International journal of physiology, pathophysiology and pharmacology 2, 73 (2010).

120 I. Hanukoglu, “Asic and enac type sodium channels: conformational states and the structures of the ion selectivity filters.” The FEBS journal 284, 525–545 (2017).

121 B. K. Berdiev, J. Xia, L. A. McLean, J. M. Markert, G. Y. Gillespie, T. B. Mapstone, A. P. Naren, B. Jovov, J. K. Bubien, H.-L. Ji, C. M. Fuller, K. L. Kirk, and D. J. Benos, “Acid-sensing ion channels in malignant gliomas.” The Journal of biological chemistry 278, 15023–15034 (2003).

122 Y. Tian, P. Bresenitz, A. Reska, L. El Moussaoui, C. P. Beier, and S. Gründer, “Glioblastoma cancer stem cell lines express functional acid sensing ion channels asic1a and asic3.” Scientific reports 7, 13674 (2017).

123 H.-Y. Yang, R.-P. Charles, E. Hummler, D. L. Baines, and R. R. Isseroff, “The epithelial sodium channel mediates the directionality of galvanotaxis in human keratinocytes.” Journal of cell science 126, 1942–1951 (2013).

124 D. Mba, M. Mycielska, Z. Madeja, S. P. Fraser, and W. Korohoda, “Directional movement of rat prostate cancer cells in direct-current electric field: involvement of voltagegated na+ channel activity.” Journal of cell science 114, 2697–2705 (2001).

125 M. B. A. Djamgoz and R. Onkal, “Persistent current blockers of voltage-gated sodium channels: a clinical opportunity for controlling metastatic disease.” Recent patents on anti-cancer drug discovery 8, 66–84 (2013).

126 M. B. A. Djamgoz, S. P. Fraser, and W. J. Brackenbury, “In vivo evidence for voltage-gated sodium channel expression in carcinomas and potentiation of metastasis.” Cancers 11(2019), 10.3390/cancers11111675.

127 K.-G. Fischer, N. Jonas, F. Poschenrieder, C. Cohen, M. Kretzler, S. Greiber, and H. Pavenstädt, “Characterization of a na(+)-ca(2+) exchanger in podocytes.” Nephrology, dialysis, transplantation: official publication of the European Dialysis and Transplant Association European Renal Association 17, 1742–1750 (2002).

128 X.-Q. Dai, A. Ramji, Y. Liu, Q. Li, E. Karpinski, and X.-Z. Chen, “Inhibition of trpp3 channel by amiloride and analogs,” Molecular pharmacology 72, 1576–1585 (2007).

129 S. Jeong, S. H. Lee, Y. O. Kim, and M. H. Yoon, “Antinociceptive effects of amiloride and benzamil in neuropathic pain model rats.” Journal of Korean medical science 28, 1238–1243 (2013).

130 A. Litan and S. A. Langhans, “Cancer as a channelopathy: ion channels and pumps in tumor development and progression,” Frontiers in cellular neuroscience 9, 86 (2015).

131 L. Leanza, A. Manago, M. Zoratti, E. Gulbins, and I. Szabo, “Pharmacological targeting of ion channels for cancer therapy: in vivo evidences,” Biochimica et Biophysica Acta (BBA)Molecular Cell Research 1863, 1385–1397 (2016).

132 A. Müller-Längle, H. Lutz, S. Hehlgans, F. Rödel, K. Rau, and B. Laube, “Nmda receptor-mediated signaling pathways enhance radiation resistance, survival and migration in glioblas-toma cells—a potential target for adjuvant radiotherapy,” Cancers 11, 503 (2019).

133 H. S. Venkatesh, W. Morishita, A. C. Geraghty, D. Silverbush, R. M. Gillespie, M. Arzt, L. T. Tam, C. Espenel, A. Ponnuswami, L. Ni, et al., “Electrical and synaptic integration of glioma into neural circuits,” Nature 573, 539–545 (2019).

134 V. Venkataramani, D. I. Tanev, C. Strahle, A. Studier-Fischer, L. Fankhauser, T. Kessler, C. Körber, M. Kardorff, M. Ratliff, R. Xie, et al., “Glutamatergic synaptic input to glioma cells drives brain tumour progression,” Nature 573, 532–538 (2019).

135 C. F. Carlborg, K. B. Gylfason, A. Kaźmierczak, F. Dortu, M. B. Polo, A. M. Catala, G. M. Kresbach, H. Sohlström, T. Moh, L. Vivien, et al., “A packaged optical slot-waveguide ring resonator sensor array for multiplex label-free assays in labs-on-chips,” Lab on a Chip 10, 281–290 (2010).

136 S. Zhao, K. Zhu, Y. Zhang, Z. Zhu, Z. Xu, M. Zhao, and T. Pan, “Electrotaxis-on-a-chip (etc): an integrated quantitative highthroughput screening platform for electrical field-directed cell migration,” Lab on a Chip 14, 4398–4405 (2014).

137 Y. Xia and G. M. Whitesides, “Soft lithography,” Angewandte Chemie International Edition 37, 550–575 (1998).

138 Y. Wang, D. Lee, L. Zhang, H. Jeon, J. E. Mendoza-Elias, T. A. Harvat, S. Z. Hassan, A. Zhou, D. T. Eddington, and J. Oberholzer, “Systematic prevention of bubble formation and accumulation for long-term culture of pancreatic islet cells in microfluidic device.” Biomedical microdevices 14, 419–426 (2012).

139 H.-F. Tsai, C.-W. Huang, H.-F. Chang, J. J. Chen, C.-H. Lee, and J.-Y. Cheng, “Evaluation of egfr and rtk signaling in the electrotaxis of lung adenocarcinoma cells under direct-current electric field stimulation,” PLoS One 8, e73418 (2013).

## References

1 C. D. McCaig, B. Song, and A. M. Rajnicek, “Electrical dimensions in cell science”, Journal of Cell Science 122, 4267–4276 (2009).

2 H.-F. Tsai, C.-W. Huang, H.-F. Chang, J. J. Chen, C.-H. Lee, and J.-Y. Cheng, “Evaluation of egfr and rtk signaling in the electrotaxis of lung adenocarcinoma cells under direct-current electric field stimulation”, PLoS One 8, e73418 (2013).

3 H.-F. Tsai, K. Toda-Peters, and A. Q. Shen, “Glioblastoma adhesion in a quick-fit hybrid microdevice”, Biomedical microdevices 21, 30 (2019).

4 C. F. Carlborg, K. B. Gylfason, A. Kaźmierczak, F. Dortu, M. B. Polo, A. M. Catala, G. M. Kresbach, H. Sohlström, T. Moh, L. Vivien, et al., “A packaged optical slot-waveguide ring resonator sensor array for multiplex label-free assays in labs-on-chips”, Lab on a Chip 10, 281–290 (2010).

5 H.-F. Tsai, S.-W. Peng, C.-Y. Wu, H.-F. Chang, and J.-Y. Cheng, “Electrotaxis of oral squamous cell carcinoma cells in a multiple-electric-field chip with uniform flow field.”, Biomicrofluidics 6, 34116 (2012).

6 S. Zhao, K. Zhu, Y. Zhang, Z. Zhu, Z. Xu, M. Zhao, and T. Pan, “Electrotaxis-on-a-chip (etc): an integrated quantitative high-throughput screening platform for electrical field-directed cell migration”, Lab on a Chip 14, 4398–4405 (2014).

7 K. W. Oh, K. Lee, B. Ahn, and E. P. Furlani, “Design of pressure-driven microfluidic networks using electric circuit analogy”, Lab on a Chip 12, 515–545 (2012).

8 D. J. Beebe, G. A. Mensing, and G. M. Walker, “Physics and applications of microfluidics in biology”, Annual Review of Biomedical Engineering 4, 261–286 (2002).

9 B. J. Kirby, Micro-and nanoscale fluid mechanics: transport in microfluidic devices (Cambridge university press, 2010).

10 T. Kihara, J. Ito, and J. Miyake, “Measurement of biomolecular diffusion in extracellular matrix condensed by fibroblasts using fluorescence correlation spectroscopy.”, PloS one 8, e82382 (2013).

